# Pseudoreplication bias in single-cell studies; a practical solution

**DOI:** 10.1101/2020.01.15.906248

**Authors:** Kip D. Zimmerman, Mark A. Espeland, Carl D. Langefeld

## Abstract

Cells from the same individual share a common genetic and environmental background and are not independent, therefore they are subsamples or pseudoreplicates. Thus, single-cell data have a hierarchical structure that many current single-cell methods do not address, leading to biased inference, highly inflated type 1 error rates, and reduced robustness and reproducibility. This includes methods that use a batch effect correction for individual as a means of accounting for within sample correlation. Here, we document this dependence across a range of cell types and show that ‘pseudo-bulk’ aggregation methods are overly conservative and underpowered relative to mixed models. We propose applying two-part hurdle generalized linear mixed models with a random effect for individual to properly account for both zero inflation and the correlation structure among measures from cells within an individual. Finally, we provide power estimates across a range of experimental conditions to assist researchers in designing appropriately powered studies.

## Introduction

The rapid evolution of single-cell technologies will enable novel interrogation of fundamental questions in biology, dramatically accelerating discoveries across many biological disciplines. Thus, researchers are developing methods that leverage or account for the unique properties of single-cell RNA sequencing (scRNA-seq) data, particularly its increased sparseness and heterogeneity compared to its bulk sequencing counterpart^1–3^. An important characteristic of single-cell experiments is that they result in many cells from the same individual, and therefore the same genetic and environmental background. Here we empirically document the correlation among measures from cells within an individual and demonstrate how testing for differential expression analysis in scRNA-seq data within a cell type across conditions without considering this correlation, the current common practice, violates fundamental assumptions and leads to false conclusions. Proper identification of the experimental unit (i.e., the smallest observation for which independence can be assumed) for the hypothesis is critical for proper inference.

Observations nested within an experimental unit are referred to as subsamples, technical replicates, or pseudoreplicates. Pseudoreplication, or subsampling, is formally defined as “the use of inferential statistics where replicates are not statistically independent”^4^. There are two types of pseudoreplication commonly occurring in single-cell experiments: simple and sacrificial. Simple pseudoreplication occurs when “samples from a single experimental unit are treated as replicates representing multiple experimental units”^4–6^. Sacrificial pseudoreplication occurs when “data for replicates are pooled prior to statistical analysis or the samples taken from each experimental unit are treated as independent replicates”^4–6^. Pseudoreplication has been addressed repeatedly in the fields of ecology, agriculture, psychology, and neuroscience and has been acknowledged as one of the most common statistical mistakes in scientific literature^4–9^.

New technologies are particularly prone to this error. Thus, it is not surprising that pseudoreplication is ubiquitous in the single-cell literature. Properly identifying the right experimental unit, and analyzing the data accordingly, needs to be urgently addressed in single-cell studies before a lack of reproducibility tarnishes single-cell’s reputation and single-cell technology itself is potentially thought of as unreliable.

In this study we simulate hierarchical single-cell expression data and evaluate the type 1 error rates and power of mixed models relative to some of the most frequently applied differential expression methods. We assess a number of commonly applied differential expression methods, but we primarily focus on the computation of a two-part hurdle model. This model explicitly accounts for the common problem of zero-inflation in scRNA-seq data by simultaneously modeling the rate of expression and the positive expression mean^10^. Using the two-part hurdle model, we compute differential expression as it is most typically applied in the literature: without a random effect for individual. We then re-evaluate the two-part hurdle model’s performance when computing differential expression with a random effect for individual, and after applying a batch effect correction for individual. Additionally, we examine the type 1 error control and power of aggregation (i.e., “pseudo-bulk”) methods, where gene expression values are averaged across all of the cells within an individual and the test statistic is computed on the individual means^11–13^. Aggregation methods are implemented to control for both zero-inflation and inter-individual heterogeneity, but represent the classic definition of sacrificial pseudoreplication. In total, these simulations inform how properly accounting for the correlation structure among measures from cells within an individual will greatly increase both robustness and reproducibility, thereby leveraging the very features that make single-cell methods powerful.

## Results

### Intra-individual Correlation

Measures from cells from the same individual should be more (positively) correlated with each other than cells from unrelated individuals. Empirically, this appears true across a range of cell types (Fig. 1). Thus, single-cell data have a hierarchical structure in which the single-cells may not be mutually independent and have a study-specific correlation (e.g., exchangeable correlation within an individual). We note, that, across a cell type, cells appear to also exhibit some correlation across individuals (Fig. 1). We hypothesize this is primarily due to both zero-inflation and the stability in functional gene expression that is needed for a cell to classify as a specific cell type (e.g., T-cells need to have some consistent signals of gene expression related to their function as T-cells).

**Fig. 1.**
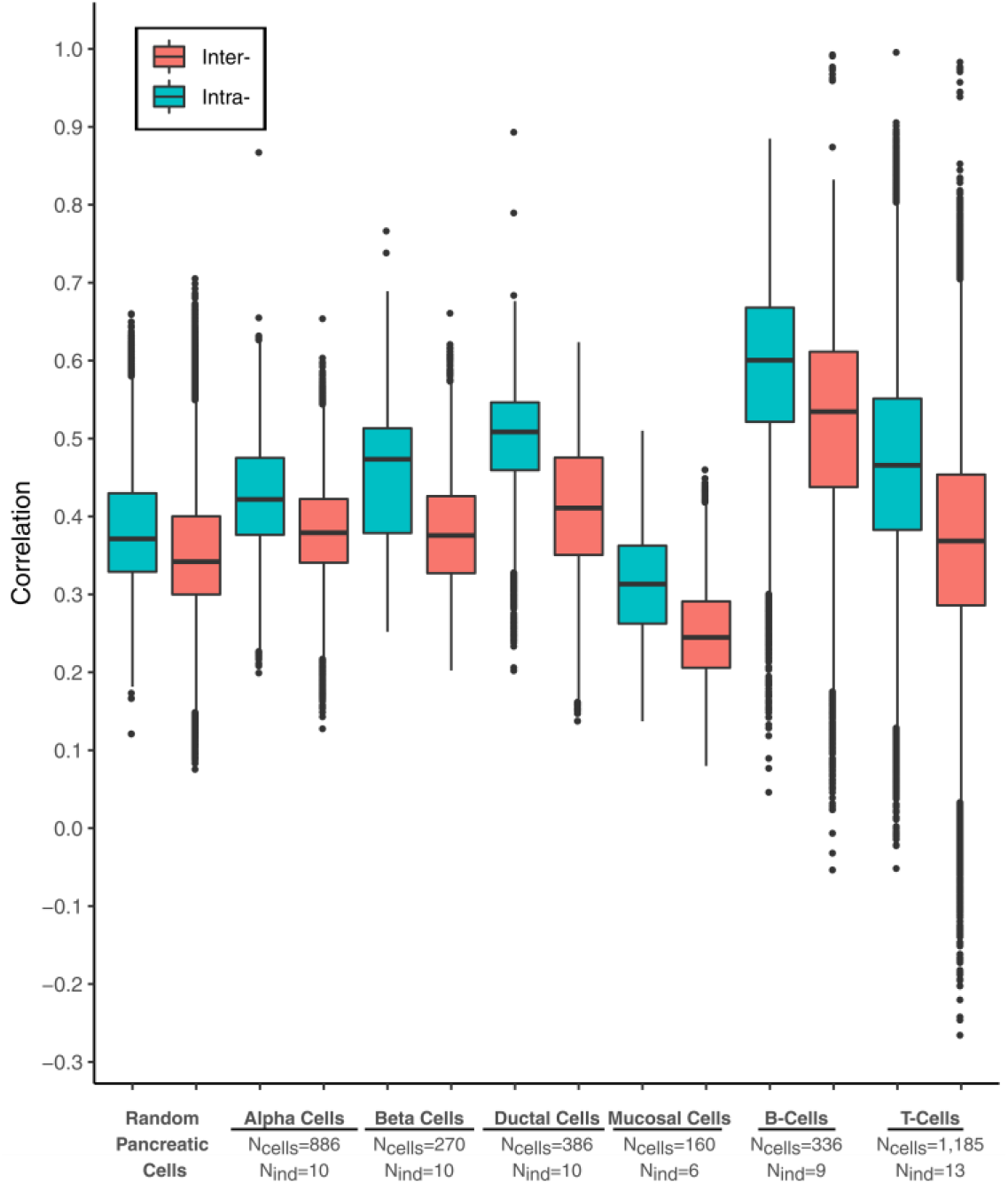
Intra-individual correlation. Box plot of the intra- and inter-individual Spearman’s correlations for gene expression values across six different cell types as well as across multiple cell types in the pancreas. Cell types, along with their respective numbers of cells (Ncells) and individuals (Nindividuals), are labeled on the x axis. Mean correlation among a donor’s own cells (intra-individual) is always greater than the mean correlation across individuals (inter-individual). Some cell types may be more correlated than others. Cell types were designated by previous authors. The center line represents the median. The lower and upper box limits represent the 25% and 75% quantiles, respectively. The whiskers extend to the largest observation within the box limit plus or minus one and a half multiplied by the interquartile range.

### Simulation

We completed a simulation study that reproduces both the inter- and intra-individual variance structures estimated from real data and documented the effect of intra-individual correlation on the type 1 error rates of the most frequently used single-cell analysis tools (Fig. 2, fig. S1-4). Our simulation captures some of the most important aspects of single-cell data (Fig. S1-4) and was used to compare methods that do and do not account for the repeated observations within an experimental unit (see Methods). We varied the number of individuals and cells within an individual and all methods considered use asymptotic approximations and admit covariates.

**Fig. 2.**
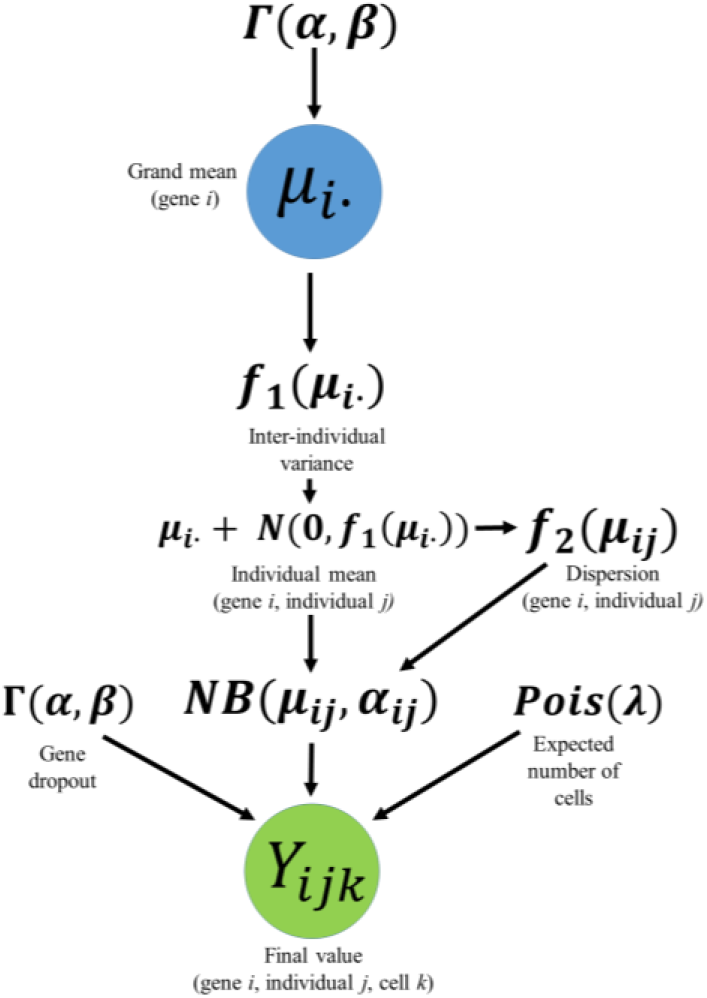
Simulation workflow. A gamma [**Γ(*α*, *β*)**] distribution was fit to the global mean normalized read counts of each gene and used to obtain a grand mean, ***μ*_*i*_**. The variance of the individual-specific means (inter-individual variance) was modeled as a linear function of the grand mean, ***f*_1_(*μ*_*i*_)**. Using a normal ***N*(*μ*, *σ*^2^)** distribution with an expected value of zero and a variance computed by the linear relationship, ***f*_1_(*μ*_*i*_)**, a difference in means was drawn for each individual in the simulation. This difference was summed with the grand mean to obtain an individual mean, *μ*_*ij*_. Within-sample dispersion was simulated as a logarithmic function of the inter-individual mean, ***f*_2_(*μ*_*ij*_)**. A Poisson (***λ***) distribution with a λ equal to the expected number of cells desired for each individual was then used to obtain the count of cells per individual. The probability of dropout was estimated as a gamma distribution. For each cell assigned to an individual, a count, ***Y*_*ijk*_**, was drawn from a negative binomial distribution.

### Type 1 Error Evaluations

We observed that the generalized linear mixed model (GLMM), either employing a tweedie distribution or a two-part hurdle model with a random effect (RE) for individual, outperformed other methods across a variety of conditions (Table 1, tables S1-S4).

**Table 1.**
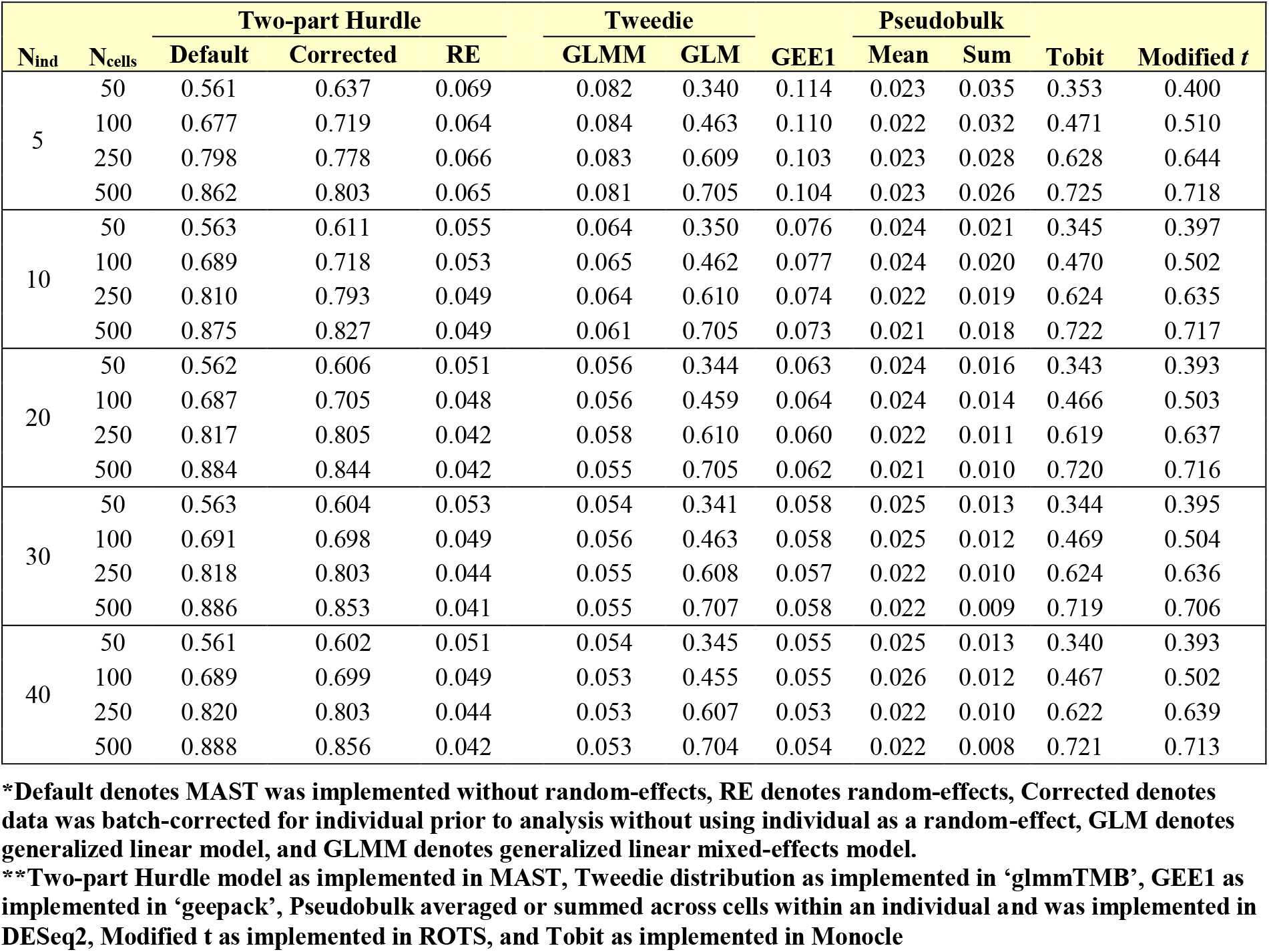
Type I error rates of some of the currently applied tools in single-cell analysis. Type I error rates of ten different methods under twenty different conditions and a significance threshold of p<0.05. 250,000 iterations were computed to obtain an error rate for each method. The inflated type I error rates computed with mixed models at the lower numbers of individuals per group are a consequence of the two-part hurdle model simultaneously testing two hypotheses and an overabundance of sub-sampling with small sample sizes. Type I error rates are well controlled for with mixed models and pseudo-bulk methods, while type I error rates inflate with other methods as additional independent samples or more cells are added. Pseudo-bulk methods are overly conservative.

Among the methods that explicitly model the correlation structure, GLMM consistently had better type 1 error rate control than generalized estimating equations (GEE1) models, where the latter performed poorly for all numbers of subsamples until the number of independent experimental units approached 30. However, all models that explicitly model the correlation structure have more appropriate type 1 error rates than the methods that do not account for the lack of independence among experimental units (Table 1, tables S1-S4). All methods that treat observations as independent perform increasingly worse as the number of correlated cells increases (Table 1, tables S1-S4). One of the most heavily cited single-cell analysis tools, Model-based Analysis of Single-cell Transcriptomics (MAST), is a two-part hurdle model built to handle sparse and bi-modally distributed single-cell data^10^. Although, to our knowledge, there are no publications that employ MAST to account for pseudoreplication as discussed here, Finak et al. note that MAST “can easily be extended to accommodate random effects”^10^. When implementing MAST with a random effect for individual (i.e., MAST with RE) the type 1 error rate is well-controlled, but its type 1 error rate is just as inflated as other tools when it is not implemented with a random effect for individual. However, one approach that has been suggested to account for the within-individual correlation is the aggregation of cell type specific expression values within an individual by using either a sum or a mean^11–13^. Such analysis methods, as would be expected, do control for the type 1 error rate, but are overly conservative (Table 1, tables S1-S4).

Another method that individuals may potentially attempt to use to account for the inter-individual heterogeneity would be to apply a batch effect correction method prior to differential expression, for which the batches are individuals. Applying batch effect correction techniques prior to differential expression analysis within a cell type, shows markedly increased type 1 error rates (Table 1, tables S1-S4).

### Power Analysis

We computed an extensive simulation-based power analysis to provide researchers estimates across a wide range of experimental conditions. This was computed using a two-part hurdle model with random effects for individuals as implemented in MAST^10^. We also computed power when expression values are averaged across cells within an individual. Increasing the number of independent experimental units (e.g., individuals) in a study is the best way to increase power to detect true differences (Fig. 3A). Empirically, when sample sizes become greater than 20, there are only marginal gains in power when more than 100 cells per individual are sampled. It is important to note that methods that aggregate information across cells within an individual by averaging (i.e., “pseudo-bulk” methods) demonstrate a large loss in power relative to mixed effects models across many different sample sizes (Fig. 3, fig. S5-S7).

**Fig. 3.**
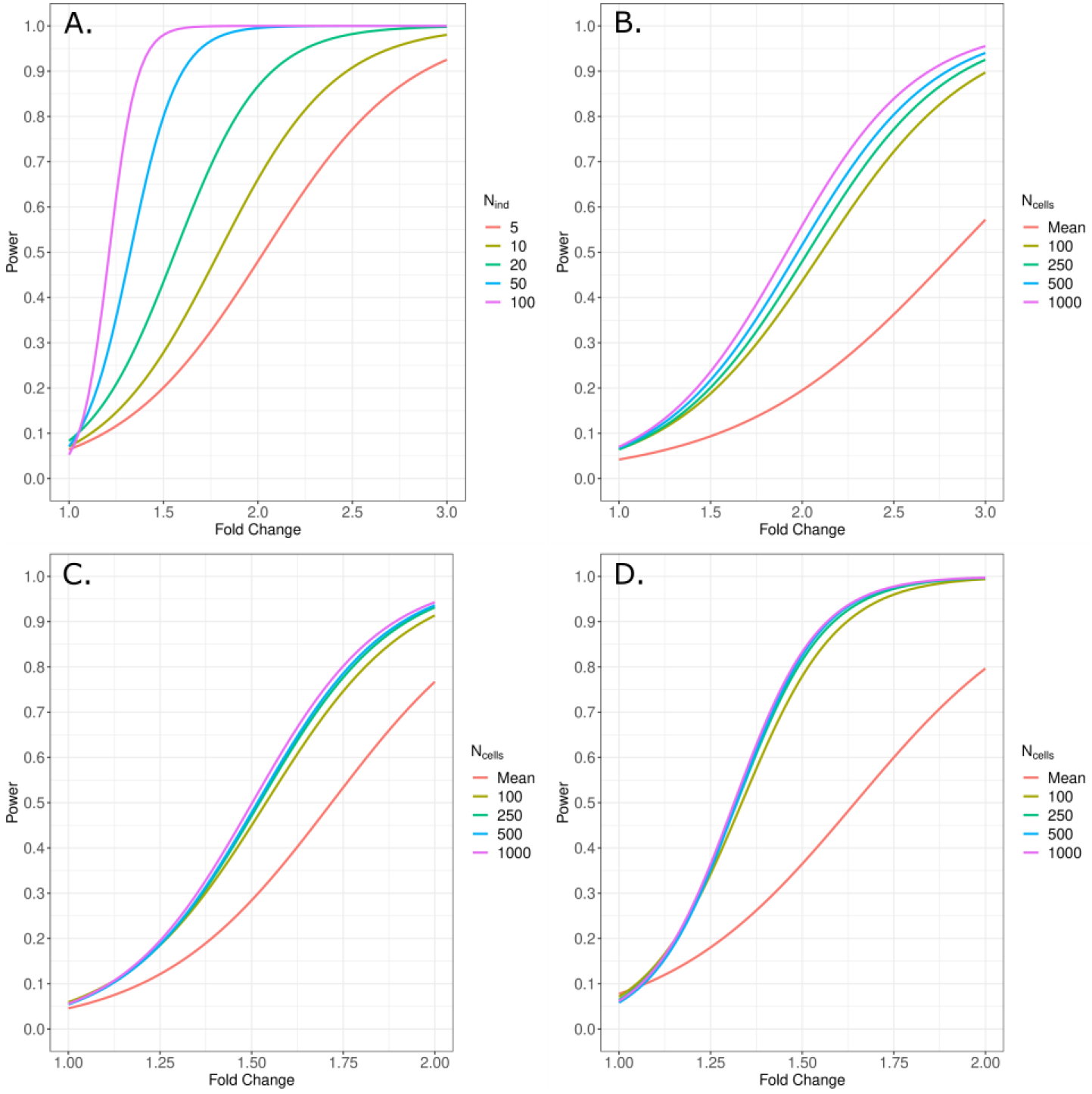
Power calculations using MAST with a random effect for individual. Power curves for various, but likely, single-cell scenarios using MAST with a random effect for individual. Fold change is simulated by multiplying the global mean gene expression values by the fold change value for one group and all power is computed at *α* = 0.05. Panel **A** demonstrates differences in power when sample sizes range between 5 individuals per group to 100 when the number of cells per individual is held constant at 250. Panels **B-D** demonstrate the differences in power when increasing the number of cells per individual (100, 250, 500, 1000) for 10, 20, and 50 individuals per group, respectively. The stark contrast in power between mixed models (100, 250, 500, 1000) and aggregation methods (Mean) is also clearly displayed. 500 cells per individual were simulated for the aggregation method. Additional power curves are supplied in the supplementary material.

## Discussion

Single-cell studies designed to identify differentially expressed genes rarely note or address the correlation among cells from the same individual or experimental unit. Excellent reviews of the field and methodological work have largely focused on challenges presented by properly classifying cell types, multimodality, dropout, and higher noise derived from biological and technical factors. However, they fail to highlight the effect of pseudoreplication and, furthermore, publications evaluating the performance single-cell specific tools all compute the simulations as if cells were independent^14–20^. The result is reduced reproducibility with real data, leading to the conclusion that tools built specifically to handle single-cell data do not appear to perform better than tools created for bulk data analysis^21–23^.

Here, we have empirically documented the correlation among measures from cells within an individual for a few independent datasets and different cell types (Figure 1). These findings imply that current practice of testing for differential expression analysis across conditions in scRNA-seq data within a cell type without considering this correlation leads to pseudoreplication. Pseudoreplication, formally defined as “the use of inferential statistics where replicates are not statistically independent”, has been addressed repeatedly in both new and well-established scientific fields^4–8^. Recently, it was acknowledged as one of the most common statistical mistakes in scientific literature^9^. Here we hope to address pseudoreplication in single-cell analysis by demonstrating what a large and long-standing body of statistical literature already confirms: applying statistical inference to replicates that are not statistically independent without properly accounting for their correlation structure will inflate type 1 error rates and lead to spurious results^4–6,9,24–27^.

In our results, the models that explicitly parameterized the correlation structure all showed improved type 1 error control over the methods that did not account for the lack of independence among experimental units (Table 1, tables S1-S4). Furthermore, all methods that treated observations independently performed increasingly worse as the number of correlated cells increased (Table 1, tables S1-S4). Both the two-part hurdle mixed model and the tweedie mixed model showed type 1 error control when adjusting for individual as a random effect (i.e., MAST with RE/Tweedie GLMM), but their type 1 error rates was highly inflated when not doing so. These specific evaluations of models with and without a random effect for individual serve as excellent example of why accounting for pseudoreplication is so important. As the denominator of most statistical tests (e.g., Wald test) is a function of the variance, not accounting for the positive correlation among sampling units underestimates the true standard error and leads to false positives^26,27^. In addition, treating each cell as independent inflates the test degrees of freedom, making it easier to falsely reject the null hypothesis (type 1 error). Too many false positives can mask true associations, especially when multiple comparison procedures such as false discovery rate are applied. In combination, this will adversely affect downstream analyses (pathway analysis), robustness, and reproducibility – increasing the cost of science.

One potential method to remove inter-individual differences prior to analysis would be to apply batch effect correction prior to differential expression analysis, for which the batches are individuals. We note that applying batch effect correction techniques to correct for intra-individual correlation should, in fact, be used more often in this way prior to cell-type clustering or prior to finding marker genes between cell-type clusters. However, when using these methods prior to differential expression analysis within a cell type, they show markedly increased type 1 error rates (Table 1, tables S1-S4). This is primarily because regressing out the person-specific effect as a batch effect and subsequently analyzing each cell as an independent observation will underestimate the overall variance by removing inter-individual differences while maintaining an inappropriately large number of degrees of freedom when treating cells as if they are independent.

Another such method that has been recommended prior to analysis is to aggregate individual gene expression values within an individual by summing or averaging them^11–13^. Such approaches, labeled “pseudo-bulk” techniques, fall under the classical definition of sacrificial pseudoreplication because they lose information. This is particularly true with respect to the within-sample variance, where heterogeneity is very likely to exist^4^. Without such information there is not a proper way to assess the significance of the difference between treatments^4^. Additionally, in imbalanced situations, “pseudo-bulk” methods cause cells from individuals with fewer cells to be more heavily weighted, where mixed models have consistent estimators and do not require balanced data^28^. The problem of imbalance is even more apparent when summing.

Overall, it has been demonstrated that mixed-effects models lead to the most accurate results when analyzing data with a hierarchical structure^6,24,25^. As we demonstrate here, aggregating values across cells from the same experimental unit will actually lead to an increased type 2 error rate and decreased power (Table 1, figure 3). This is due to an overestimation of the mean-square error relative to mixed-effects models, particularly when the intra-individual variance is larger than the inter-individual variance, as appears to be the case with scRNA-seq data^24,25^. ‘Pseudo-bulk’ techniques may control the type 1 error rates and may help remove zero-inflation, however, we strongly recommend mixed-effects models based on long-standing statistical justifications for the analysis of subsamples, including increased power.

Among the methods that explicitly model the correlation structure, generalized linear mixed models (GLMM) consistently had better type 1 error rate control than generalized estimating equations (GEE1) models, where the latter performed poorly for any number of subsampling until the number of independent experimental units approached 30. When the number of experimental units was small, the GEE1 sandwich estimator of the variance provided standard errors that were too small and therefore inflated the type 1 error rate^29,30^. Here, we emphasize the two-part hurdle mixed model, MAST, as an already well-established tool in the field and we demonstrate that MAST performs exceptionally well when adjusting for individual as a random effect^10^. We note, that MAST with RE is testing a two-part hypothesis that the other tools are not directly testing. The discrete and continuous components being tested fit together in the sense that higher mean expression will generally correlate with a higher proportion of expressing cells, but making the assumption that the two will always relate is incorrect. There will be specific instances in the simulated data when the inter-individual means are not significantly different, but the proportion of cell dropout is significantly different (even though the probability of dropout for any one gene across cells is held constant) and is driving a significant result. This will be particularly true with smaller sample sizes, and may contribute to the slightly elevated type 1 error rates with smaller sample sizes and cell counts. While we do recommend computing differential expression analysis using MAST with RE, alternative methods include the tweedie GLMM or permutation testing. In order not to violate the exchangeability assumption, permutation methods must randomize at the independent experimental unit (e.g., individual) and properly account for covariates (i.e., conditional permutation). The tweedie GLMM method could be implemented using the ‘glmmTMB’ R-package^31^, but neither of these alternative approaches explicitly incorporate some of the single-cell specific concepts implemented in MAST (e.g., cellular detection rate).

We computed power analyses for a variety of sample sizes and cell amounts. Empirically, we demonstrated that increasing the amount of cells captured per sample returns very little gain in power after 100 cells per individual in most scenarios, particularly after sample sizes increase (Figure 3, fig. S5-S7). Instead, we suggest that increasing sample size is the most efficient way to improve power (Figure 3A). Increasing the number of cells per individual does provide more precision in the estimate for an individual. However, it has limited effects on the power for detecting differences across individuals, such as differences among treatments applied to individuals (i.e., cases/control studies). We note that estimating power with more than 1,000 cells per individual is computationally expensive. Because 1,000s of cells per individual is not atypical for single-cell experiments, tools that account for the correlation structure when analyzing these data need to be further developed to increase computational efficiency.

Most papers compare cells across very few individuals, sometimes even a single case and control (simple pseudoreplication). In the former case the estimate of the inter-individual variance is possible but has wide bounds on parameter confidence intervals, and in the latter case the variance is not estimable from the data. Simulations indicate that the majority of published studies are underpowered (Fig. 3, fig. S5-S7). The majority of single-cell papers show a deep understanding of the underlying biology and conduct otherwise very informative experiments, appropriately landing in very high visibility journals. However, our type 1 error and power simulations document that many published studies are missing important true effects while reporting too many false positives generated via pseudoreplication.

As single-cell technology continues to evolve and costs decrease, scientists need to be aware of this issue to improve study design and avoid proliferation of irreproducible results. Our results encourage the use of mixed models, such as the two-part hurdle model with a random effect (e.g., as implemented in MAST with RE), as ways of accounting for the repeated observations from an individual while being able to adjust for covariates at the individual level and, if appropriate, at the individual cell level. Additional random effects, such as sampling time, may also be included^32^. Our extensive simulation study provides valuable information for understanding the power of specific designs and can be used in grant reviews as one justification of the design and analyses employed. Finally, we note that although our focus here is on hypothesis testing for finding differentially expressed genes within a cell type across conditions, the concept is applicable when comparing expression patterns between cell-types and is broadly appropriate for all single-cell sequencing technologies such as proteomics, metabolomics, and epigenetics.

## Methods

### Literature Review

A PubMed search in January of 2019 for the keywords “single-cell differential expression” returned 251 articles published in the last 3 years which were subsequently sorted and filtered by each of their abstracts. Many of the returned articles were associated with bulk RNA sequencing or completely irrelevant to differential expression analysis in single-cell and were therefore eliminated. Of the 251 original hits, 76 of them were deemed appropriate for further consideration. Of those, 10 of them were reviews, 36 of them were methods papers, and 30 of them were implementation papers. This method is not meant to be a perfect capture of all of the literature, but provides a clear snapshot of the current state of single-cell differential expression analyses. Each of the methods and implementation articles was thoroughly reviewed and tabled along with its number of citations, date of publication, and any other pertinent information such as number of independent samples, tools used, or number of cells captured.

### Intra and Inter-correlation analyses

Pairwise comparisons between all cells of interest were made to compute intra- and inter-individual correlations. Genes were filtered if the average transcript-per-million (TPM) value was less than five. To control for the correlation structure between genes, genes were sampled one at a time and any genes with a Spearman’s correlation coefficient > 0.25 relative to the gene that was drawn were subsequently trimmed from the dataset. This was repeated until either no more uncorrelated genes remained or a total of 500 uncorrelated genes were obtained, whichever happened first. For intra-individual correlation, Spearman’s correlation was computed for all possible pairs of cells within an individual. For inter-individual correlation, Spearman’s correlation was computed for all possible pairs of cells from a random draw of one cell from each individual. 1,000 draws were computed. To compute the correlation structure across multiple cell types, intra-individual correlation was assessed by repeatedly drawing one cell per cell type within an individual and computing all pairwise correlations. Inter-individual correlation was assessed by sub-setting the data to a balanced set of observations with exactly ten cells of each of the three main cell types being retained for each individual. Correlations and their means were examined for differences (Fig. 1). The measures were compared in six different cell types across three different single-cell studies. These studies are publically available under the accession numbers GSE81861, GSE72056, and E-MTAB-5061. The GSE81861 dataset contains 161 normal mucosal cells from 6 individuals and were sequenced on Fluidigm’s C1.

The GSE72056 data were also sequenced on Fluidigm’s C1 and the data used here contains 337 B-cells and 1,186 T-cells from tumor tissue of 11 individuals. The E-MTAB-5061 data were sequenced using the Smart-Seq2 protocol and contains cells pancreatic cells taken from 10 individuals. Here, only data from 886 alpha cells, 270 beta cells, and 336 ductal cells were used. The cell type designations that were used were given by the authors of these studies and more detailed information is provided in their respective manuscripts.

### Simulation

To control for the correlation structure between genes, genes were sampled one at a time and any genes with a Spearman’s correlation coefficient > 0.25 relative to the gene that was drawn were subsequently trimmed from the dataset. This was repeated until either no more uncorrelated genes remained or a total of 500 uncorrelated genes were obtained, whichever happened first. Means and variances were computed empirically from the normalized read count values previously reported in six different cell types across three different single-cell studies. Once consistent patterns were identified across cell types, alpha cells from the pancreas dataset, were used as the model data for our simulation. A gamma distribution was fit to the global mean of the normalized read count values of each gene using maximum-likelihood estimation to obtain a grand mean, *μ*_*i*_. The variance of the individual-specific means (inter-individual variance) was modeled as a linear function of the grand mean, *f*_1_(*μ*_*i*_) and the within-sample dispersion (intra-individual variance) was estimated as a logarithmic function of the inter-individual means, *f*_2_(*μ*_*i*_), and the probability of dropout was estimated independently as a gamma distribution (Fig. 2). The objective for obtaining the inter-individual variance as function of the grand mean was to simply capture the average associations between the variance and the grand mean - the uncertainty in this relationship was not of particular interest for this simulation. Using a normal distribution with an expected value of zero and a variance computed by the first linear relationship, *f*_1_(*μ*_*i*_), a difference in means was drawn for each individual in the simulation. This difference was summed with the grand mean to obtain an individual mean, *μ*_*ij*_. A Poisson distribution with a λ equal to the expected number of cells desired for each individual was then used to obtain the count of cells per individual. For each cell assigned to an individual, a read count value, *Y*_*ijk*_, was drawn from a negative binomial distribution with an expected value equal to the individual’s assigned read count value, *μ*_*ij*_, and a dispersion parameter, *α*_*ij*_, computed by the logarithmic function of the grand mean *f*_2_(*μ*_*i*_). Along with the distributions of the primary parameters of interest, tSNE plots were made of the simulated data to assess how realistic the simulated data appeared (Fig S1-4).

### Type I error

In total, 250,000 iterations of our simulation were computed for varying numbers of cells and varying numbers of individuals. The number of individuals per group was fixed at either 5, 10, 20, 30 or 40. The number of cells per individual was drawn from a Poisson distribution with either a λ of 50, 100, 250, or 500. For each of the 250,000 iterations, the number of results that met our significance threshold were counted and the type I error was computed as the percentage of significant results. After analysis of the type I error using a tweedie mixed-effects model^31^, type I error was computed with the following tools: MAST^10^, MAST with random effects^10^, MAST with a batch effect correction^10,33^, DESeq2 with aggregation methods (‘Pseudo-bulk’ summing and averaging)^11–13,34^, Monocle^35^, ROTS^36^, Tweedie GLM^31^, and a GEE1 with a Gaussian link and exchangeable correlation^37^. MAST was implemented with and without the use of a random effect for individual and the remaining single-cell tools were implemented exactly as their vignettes instruct. GEE1 with exchangeable correlation was implemented to compare its performance to the mixed-effects model, particularly where the numbers of donors are low. Type I errors were computed using significance thresholds of 0.05, 0.01, 0.001, and 0.0001 (Table 1, tables S1-S4).

MAST models a log(x + 1) transformed gene expression matrix as a two-part generalized regression model^10^. As in this cited manuscript, the addition of random effects for differences among persons is:

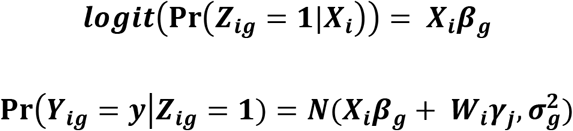

where *Y*_*ig*_ is the expression level for gene *g* and cell *i*, *Z*_*ig*_ is an indicator for whether gene *g* is expressed in cell *i*, *X*_*i*_ contains the predictor variables for each cell *i*, and *W*_*i*_ is the design matrix for the random effects of each cell *i* belonging to each individual *j* (i.e., the random complement to the fixed *X_i_*). β_*g*_ represents the vector of fixed-effects regression coefficients and γ_*j*_ represents the vector of random effects (i.e., the random complement to the fixed β_*g*_). γ_*j*_ is distributed normally with a mean of zero and variance 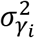. To obtain a single result for each gene, likelihood ratio or Wald test results from each of the two components are summed and the corresponding degrees of freedom for each component are added^10^. These tests have asymptotic χ^2^ null distributions and they can be summed and remain asymptotically χ^2^ because *Z*_*g*_ and *Y*_*g*_ are defined conditionally independent for each gene^10^.

### Power calculations

Using MAST with a random effect for individual, we computed power curves to estimate how well this tool functions with varying numbers and ratios of cells and individuals.

Computations were identical to the type I error analyses with exception of multiplying a constant, hereafter labeled fold change, with the global mean gene expression value of a gene to spike the expression values in one group. Power was computed at small increments between a fold change of 1 and 5 and each value was used fit a sigmoidal curve to approximate power (Fig. S4-S7).

#### Code availability

Data were simulated in R-3.5.1. All of the code for the simulations and the evaluation of intra- and inter-individual correlation structure is available on GitHub at kdzimmer/PseudoreplicationPaper. This base code was modified to run type I error and power analyses in parallel on Wake Forest’s High Performance Computing Cluster, DEMON.

## Acknowledgments

We thank T. D. Howard, L. D. Miller, and M. Olivier (All WFU School of Medicine) for critical review of this content. This work was supported by The Center for Public Health Genomics and grant U01 NS036695 (Co-PI Langefeld) from NIH.

## Author contributions

C.D.L. and K.D.Z conceived the study together. K.D.Z implemented simulations and analyses with guidance from C.D.L. K.D.Z wrote the original draft and reviewed and edited it with M.A.E. and C.D.L.

## Data and materials availability

All data are publically available under the accession numbers GSE81861, GSE72056, and E-MTAB-5061. All base code is available on GitHub at kdzimmer/PseudoreplicationPaper

## Supplementary Materials

Figures S1-S7 (tSNE and power curves)

Tables S1-S4

