## Supplemental Materials for "Pseudoreplication bias in single-cell studies; a practical solution"

**This PDF file includes:**

Figs. S1 to S7
Tables S1 to S4

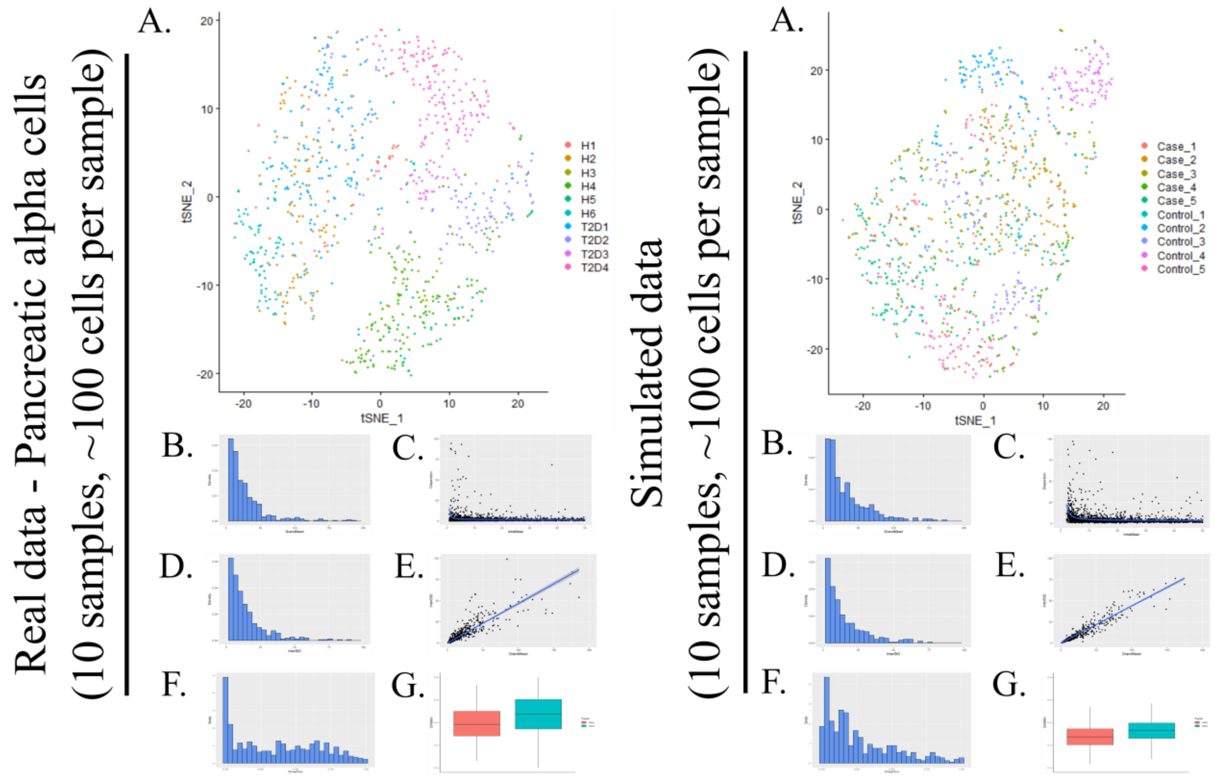


**Fig. S1. | Similarity between simulated data and real data**. tSNE plots and plots of various parameters demonstrate the similarity between simulated data and real data. All axes, with the exception of the tSNE plot axes, are held constant across the two sets of panels. Estimation of the parameters was completed on pancreatic alpha cells with 500 genes correlated with a Spearman's coefficient < 0.25. The simulated data consists of 500 independently simulated genes. Panels **A**. represent the tSNE plots of the data. Panels **B**. are histograms of the grand mean and panels **C**. demonstrate the relationship between intra-individual means and intra-individual dispersion. Panels **D**. are histograms of the inter-individual variance and panels **E**. show the relationship between the grand mean and the inter-individual variance. **F**. shows the distribution of dropout and **G**. demonstrates the distributions of the intra- and inter-individual correlations where intra-individual correlation is blue and inter-individual correlation is red. The difference between inter-individual correlation and intra-individual correlation is slightly less exaggerated in our simulated data.

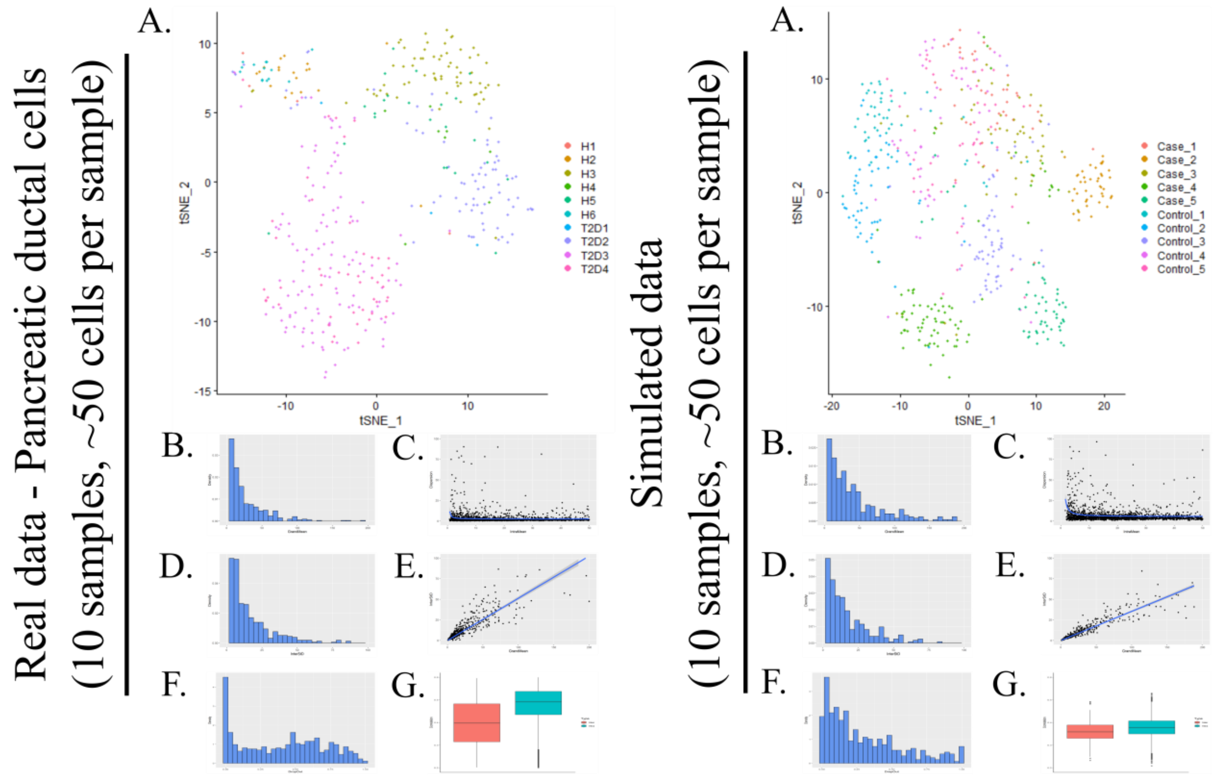


**Fig. S2. | Similarity between simulated data and real data**. tSNE plots and plots of various parameters demonstrate the similarity between simulated data and real data. All axes, with the exception of the tSNE plot axes, are held constant across the two sets of panels. Estimation of the parameters was completed on pancreatic ductal cells with 500 genes correlated with a Spearman's coefficient < 0.25. The simulated data consists of 500 independently simulated genes. Panels **A**. represent the tSNE plots of the data. Panels **B**. are histograms of the grand mean and panels **C**. demonstrate the relationship between intra-individual means and intra-individual dispersion. Panels **D**. are histograms of the inter-individual variance and panels **E**. show the relationship between the grand mean and the inter-individual variance. **F**. shows the distribution of dropout and **G**. demonstrates the distributions of the intra- and inter-individual correlations where intra-individual correlation is blue and inter-individual correlation is red. The difference between inter-individual correlation and intra-individual correlation is slightly less exaggerated in our simulated data.

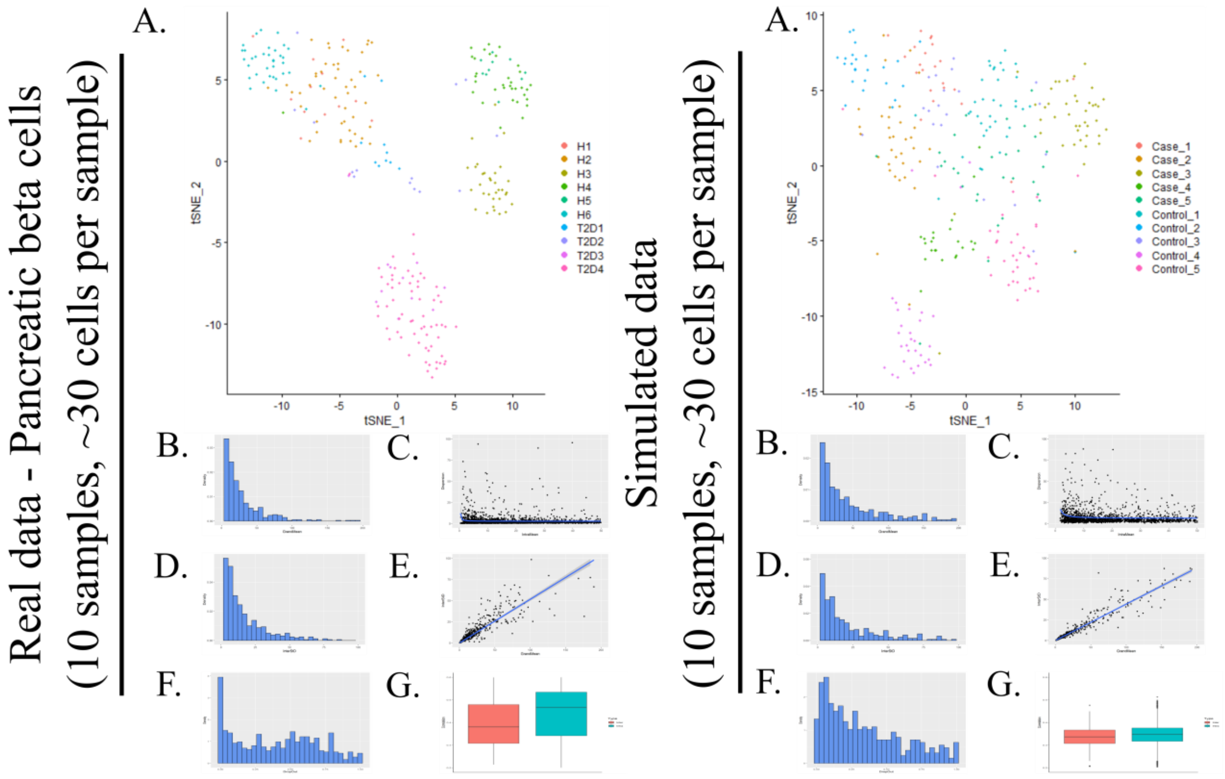


**Fig. S3. | Similarity between simulated data and real data**. tSNE plots and plots of various parameters demonstrate the similarity between simulated data and real data. All axes, with the exception of the tSNE plot axes, are held constant across the two sets of panels. Estimation of the parameters was completed on pancreatic beta cells with 500 genes correlated with a Spearman's coefficient < 0.25. The simulated data consists of 500 independently simulated genes. Panels **A**. represent the tSNE plots of the data. Panels **B**. are histograms of the grand mean and panels **C**. demonstrate the relationship between intra-individual means and intra-individual dispersion. Panels **D**. are histograms of the inter-individual variance and panels **E**. show the relationship between the grand mean and the inter-individual variance. **F**. shows the distribution of dropout and **G**. demonstrates the distributions of the intra- and inter-individual correlations where intra-individual correlation is blue and inter-individual correlation is red. The difference between inter-individual correlation and intra-individual correlation is slightly less exaggerated in our simulated data.

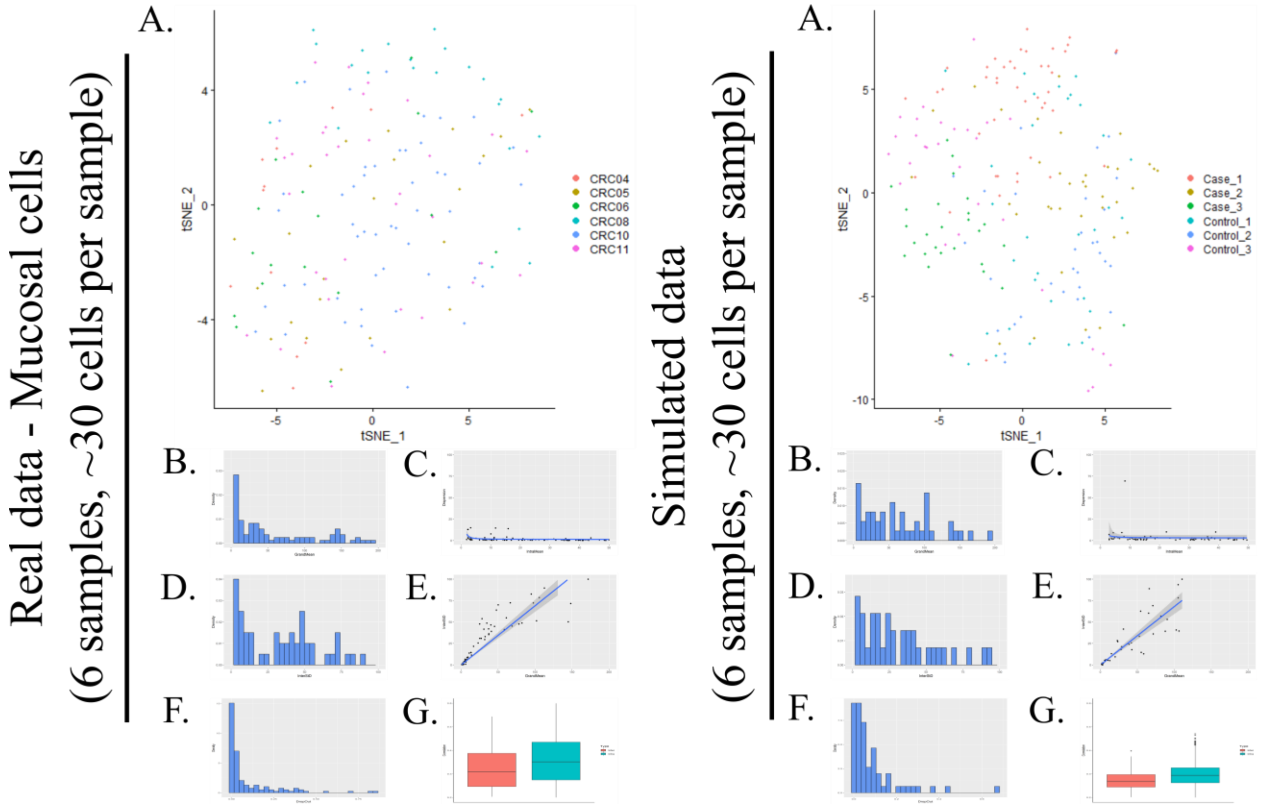


**Fig. S4. | Similarity between simulated data and real data**. tSNE plots and plots of various parameters demonstrate the similarity between simulated data and real data. All axes, with the exception of the tSNE plot axes, are held constant across the two sets of panels. Estimation of the parameters was completed on normal mucosal cells with 215 genes correlated with a Spearman's coefficient $< 0.25$. The simulated data consists of 215 independently simulated genes. Panels **A**. represent the tSNE plots of the data. Panels **B**. are histograms of the grand mean and panels **C**. demonstrate the relationship between intra-individual means and intra-individual dispersion. Panels **D**. are histograms of the inter-individual variance and panels **E**. show the relationship between the grand mean and the inter-individual variance. **F**. shows the distribution of dropout and **G**. demonstrates the distributions of the intra- and inter-individual correlations where intra-individual correlation is blue and inter-individual correlation is red. The difference between inter-individual correlation and intra-individual correlation is slightly less exaggerated in our simulated data.

**Fig. S5. | Power calculations using MAST with a random effect for individual.** Power curves for MAST using a random effect to account for intra-individual correlation. Curves are computed for 100, 250, 500, and 1,000 cells per individual using an $\alpha = 0.05$. The number of individuals per group ranges from 3 to 100 and is

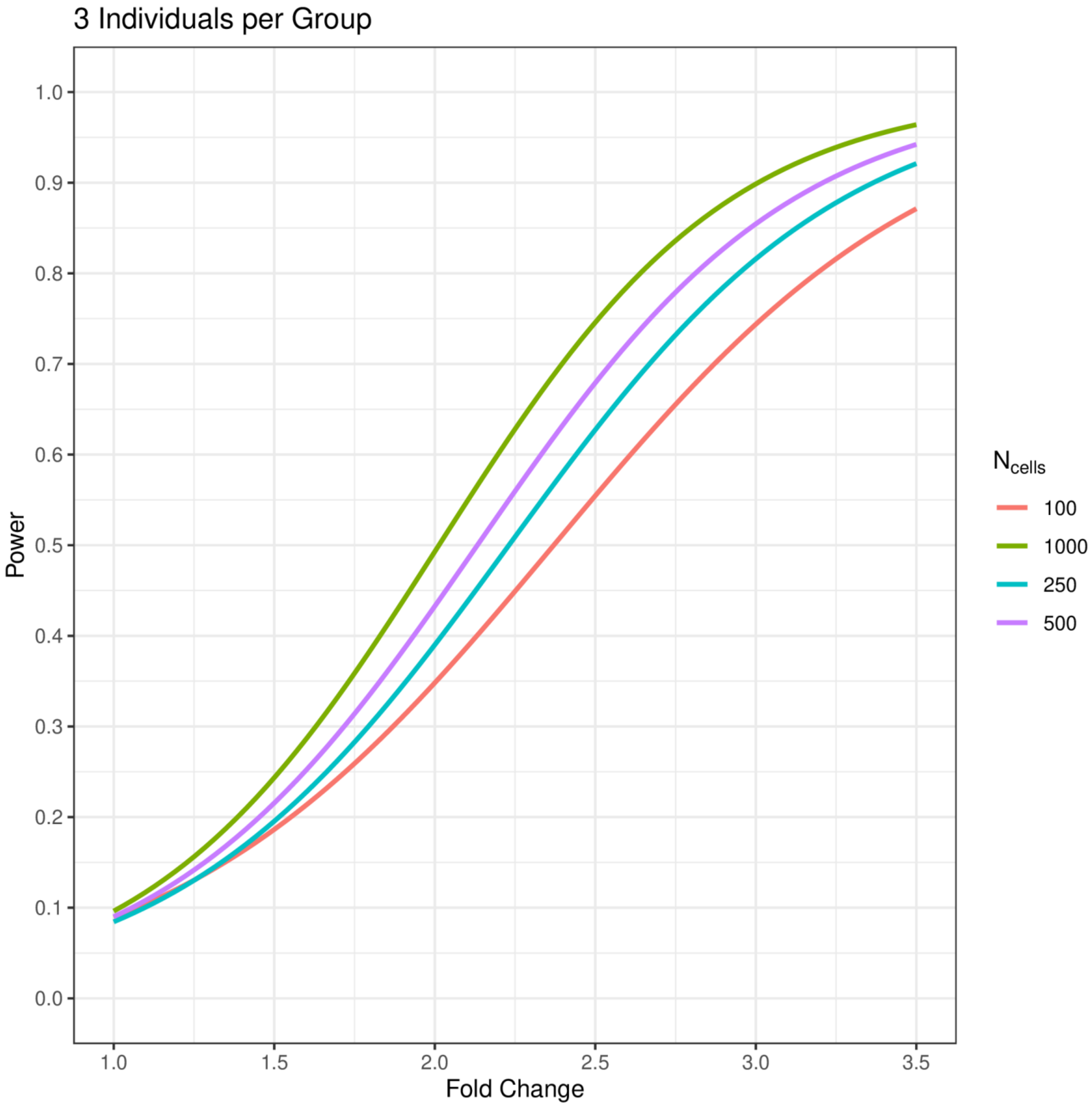

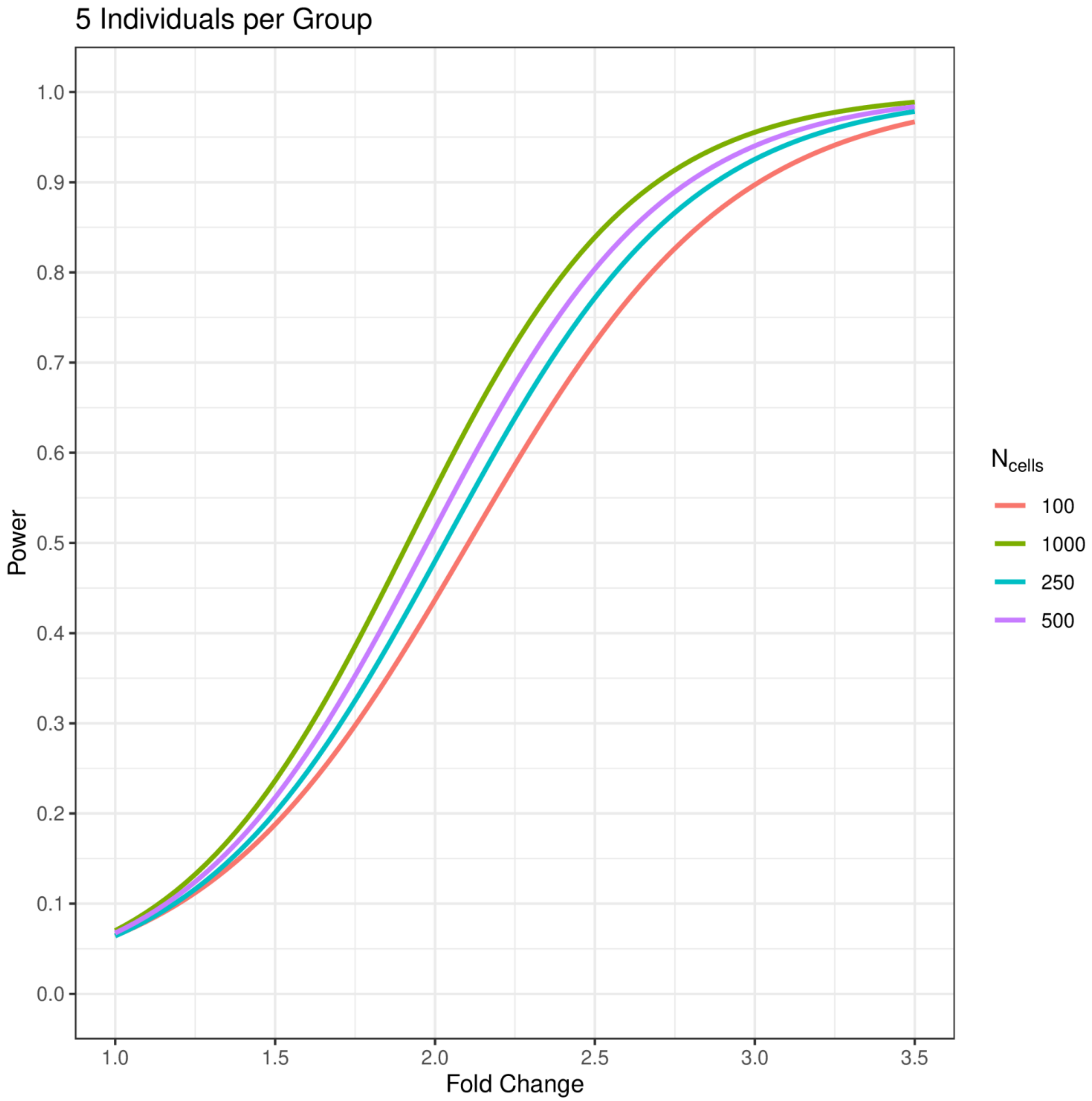

5 Individuals per Group
1.0
0.9
0.8
0.7
0.6
0.5
0.4
0.3
0.2
0.1
0.0
Power
1.0
1.5
2.0
2.5
3.0
3.5
Fold Change
$N_{cells}$
100
1000
250
500

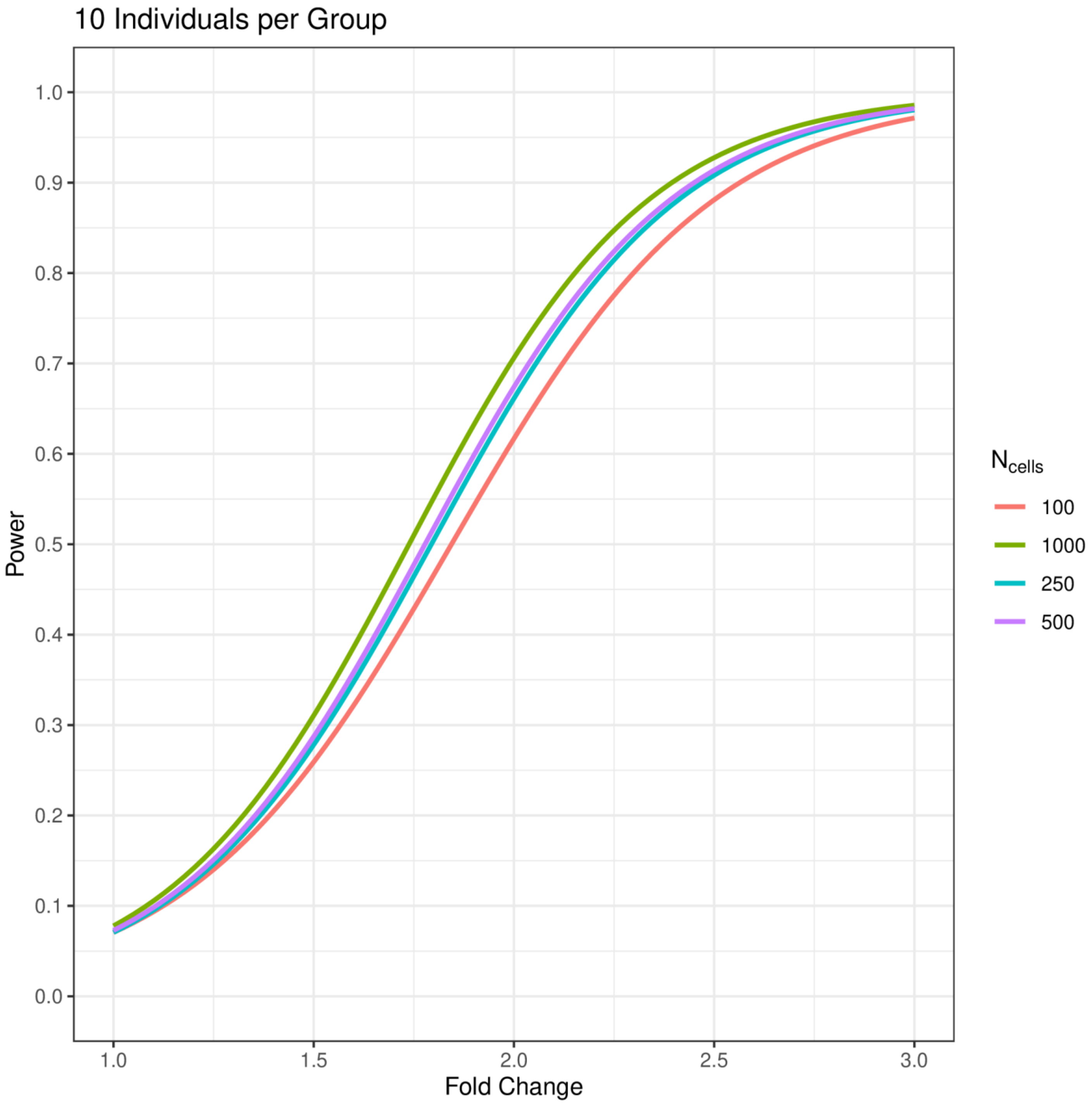
10 Individuals per Group
Power
Fold Change
$N_{cells}$
100
1000
250
500
1.0
0.9
0.8
0.7
0.6
0.5
0.4
0.3
0.2
0.1
0.0
1.0
1.5
2.0
2.5
3.0

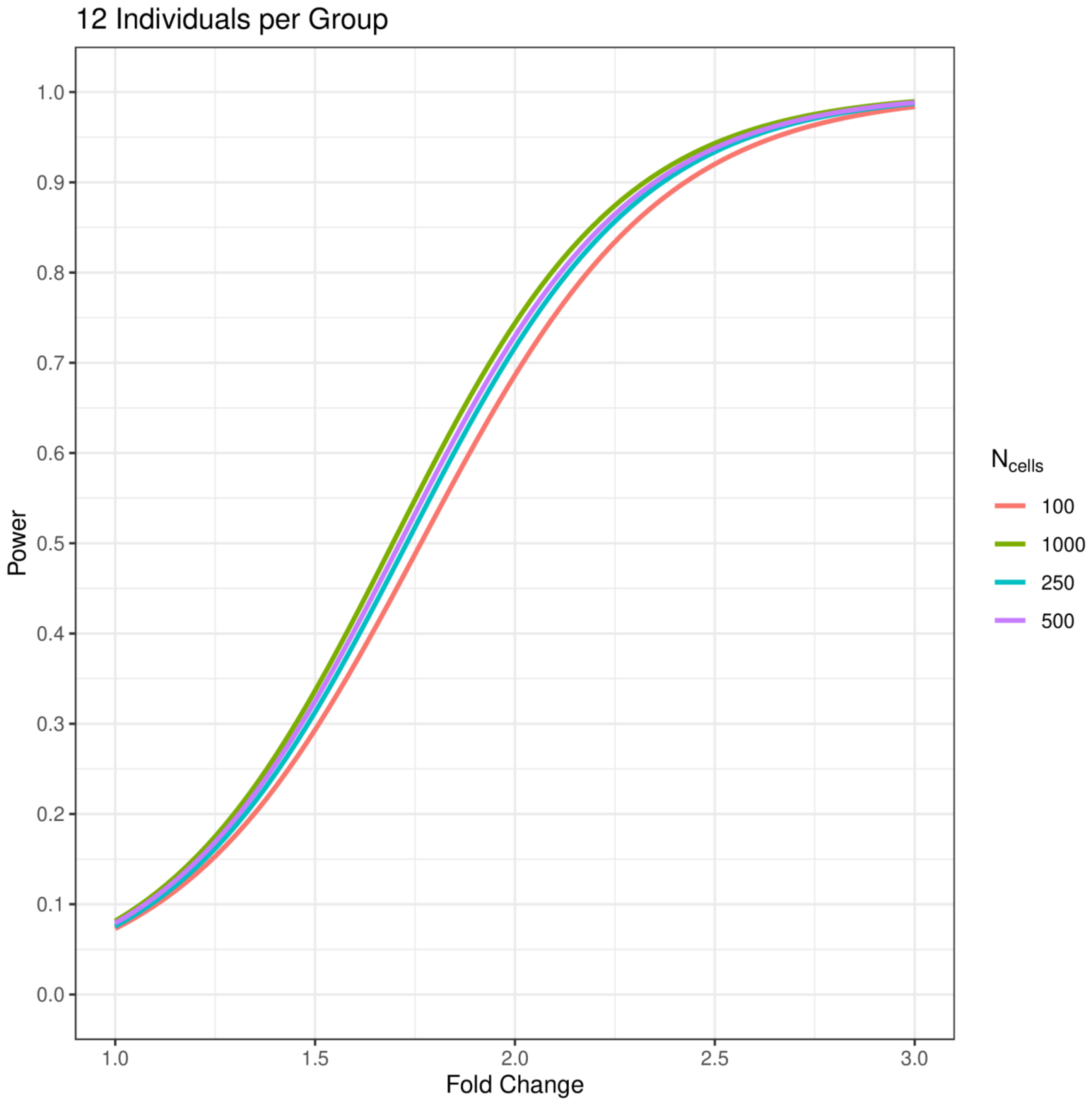

12 Individuals per Group
Power
Fold Change
1.0
0.9
0.8
0.7
0.6
0.5
0.4
0.3
0.2
0.1
0.0
1.0
1.5
2.0
2.5
3.0
$N_{cells}$
100
1000
250
500

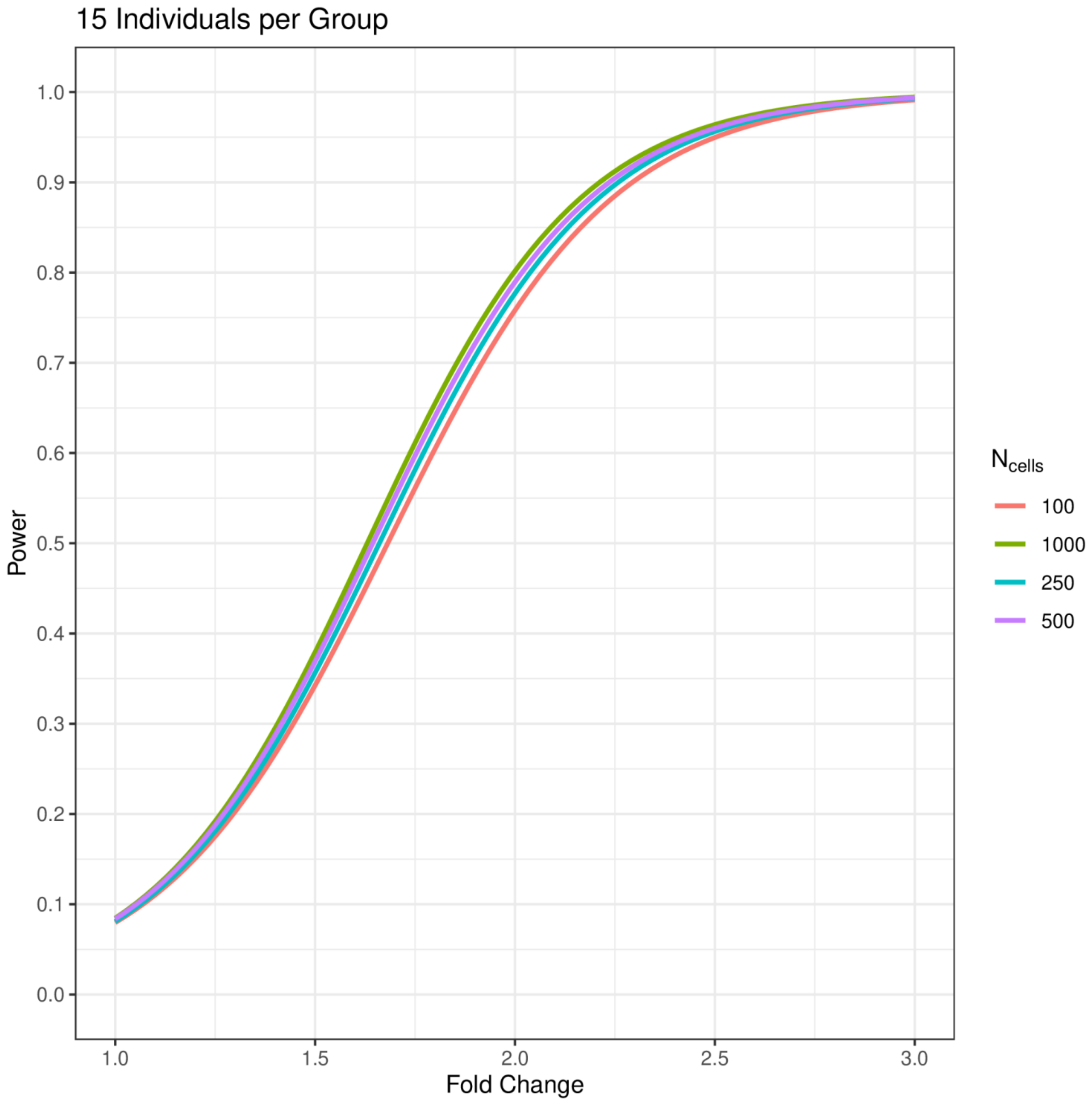
15 Individuals per Group
Power
1.0
0.9
0.8
0.7
0.6
0.5
0.4
0.3
0.2
0.1
0.0
1.0
1.5
2.0
2.5
3.0
Fold Change
N cells
100
1000
250
500

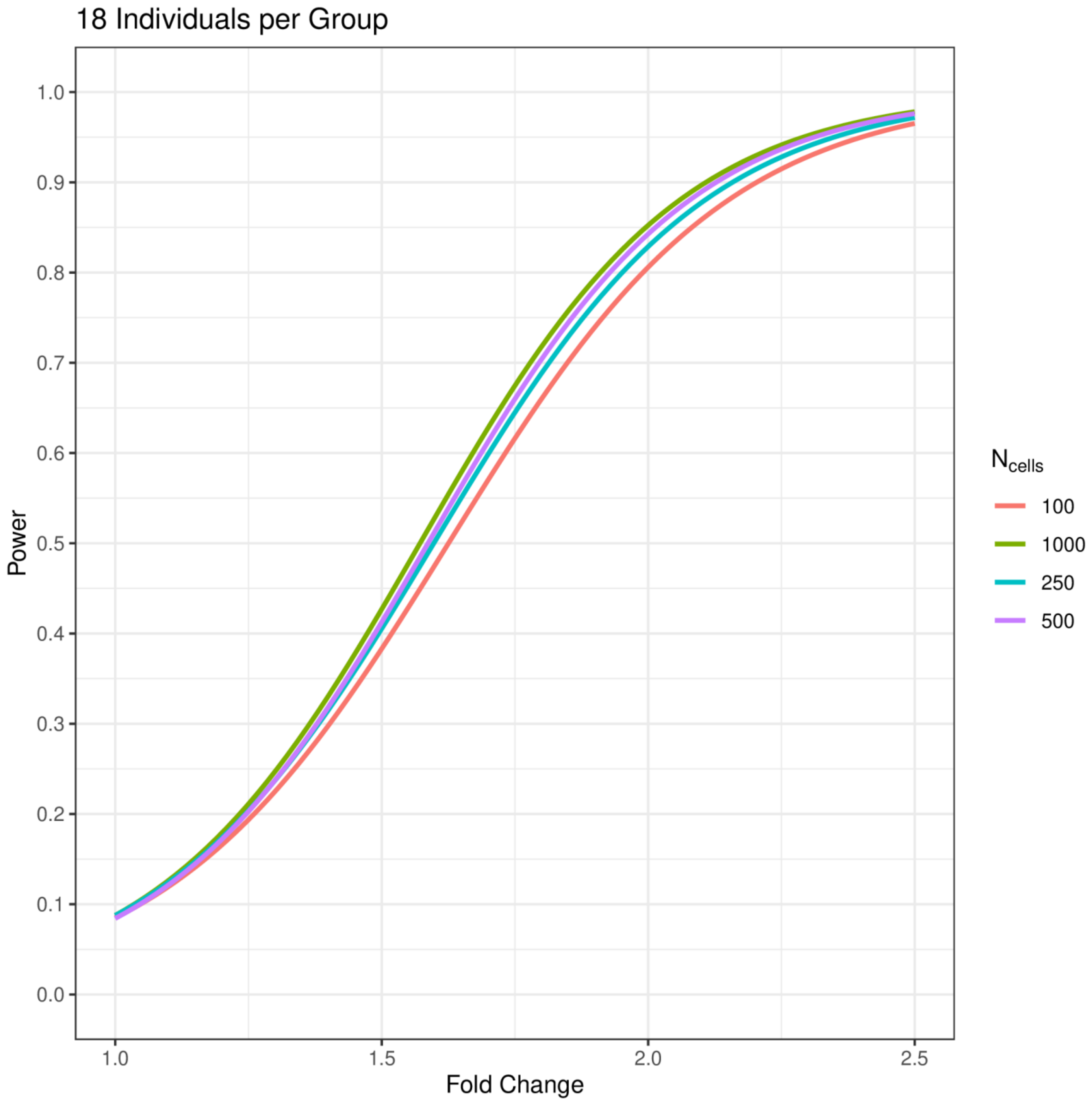

18 Individuals per Group
Power
1.0
0.9
0.8
0.7
0.6
0.5
0.4
0.3
0.2
0.1
0.0
1.0
1.5
2.0
2.5
Fold Change
N cells
100
1000
250
500

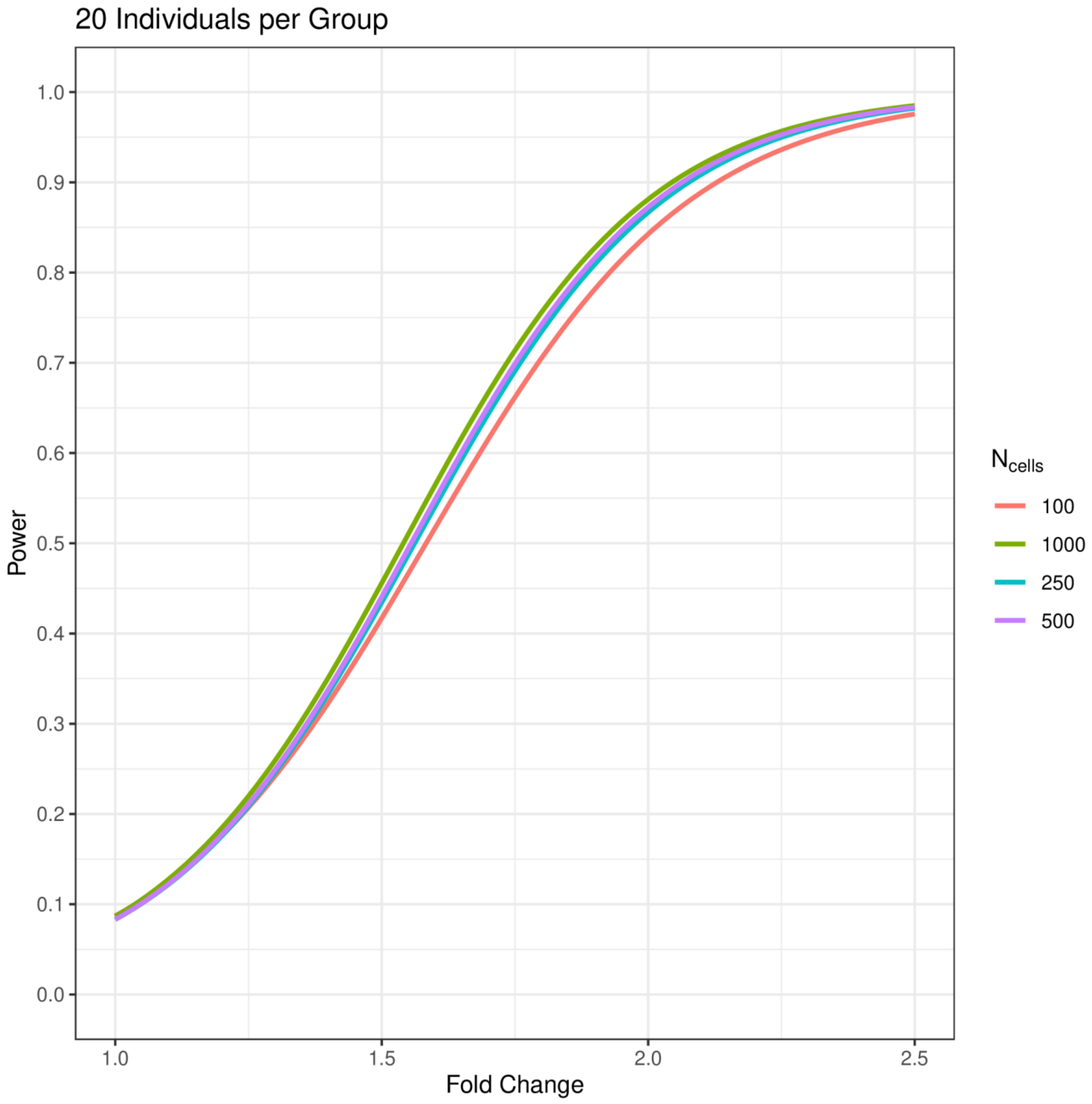

20 Individuals per Group
Power
1.0
0.9
0.8
0.7
0.6
0.5
0.4
0.3
0.2
0.1
0.0
1.0
1.5
2.0
2.5
Fold Change
N cells
100
1000
250
500

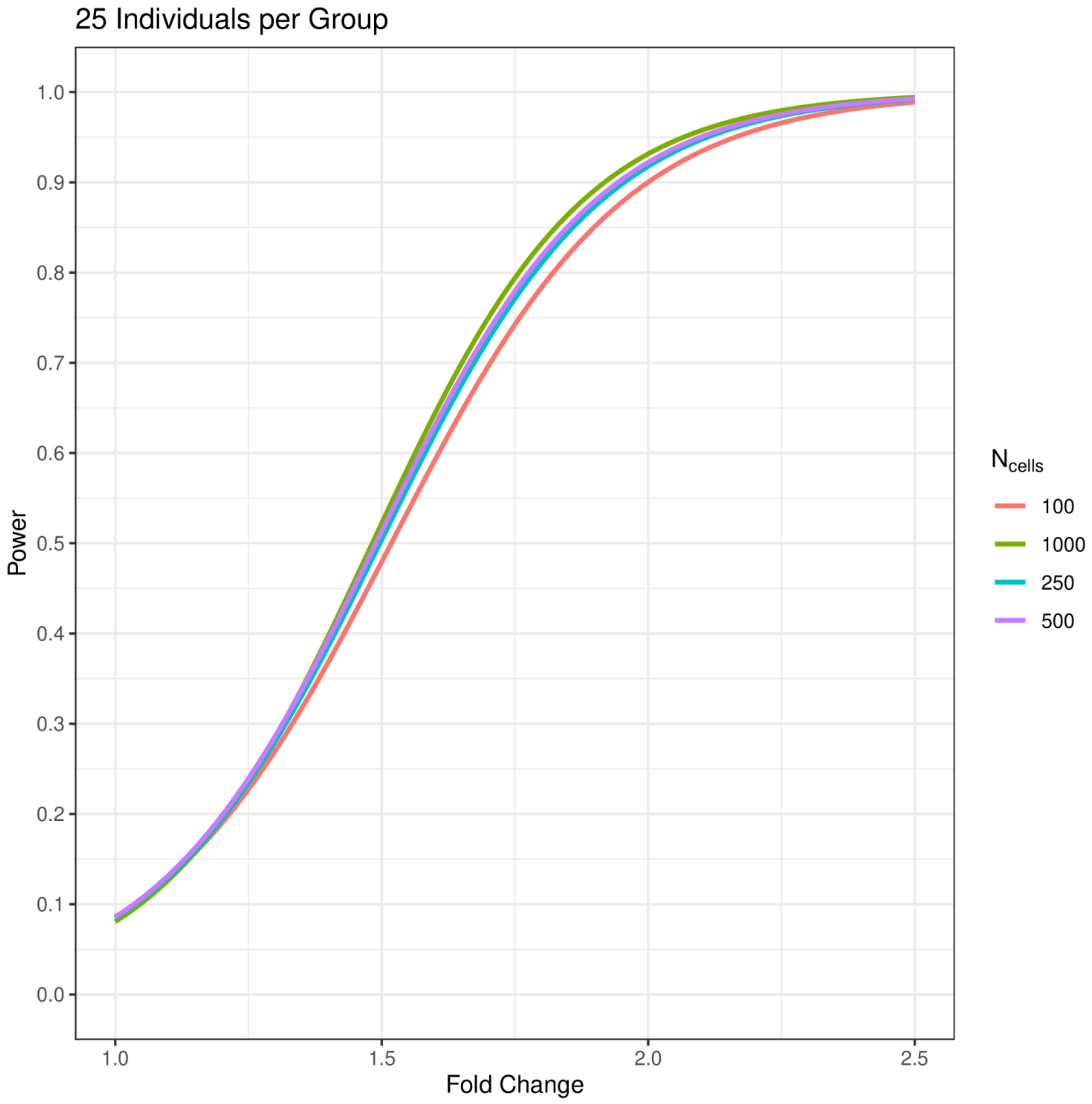
25 Individuals per Group
Power
Fold Change
1.0
0.9
0.8
0.7
0.6
0.5
0.4
0.3
0.2
0.1
0.0
1.0
1.5
2.0
2.5
$N_{cells}$
100
1000
250
500

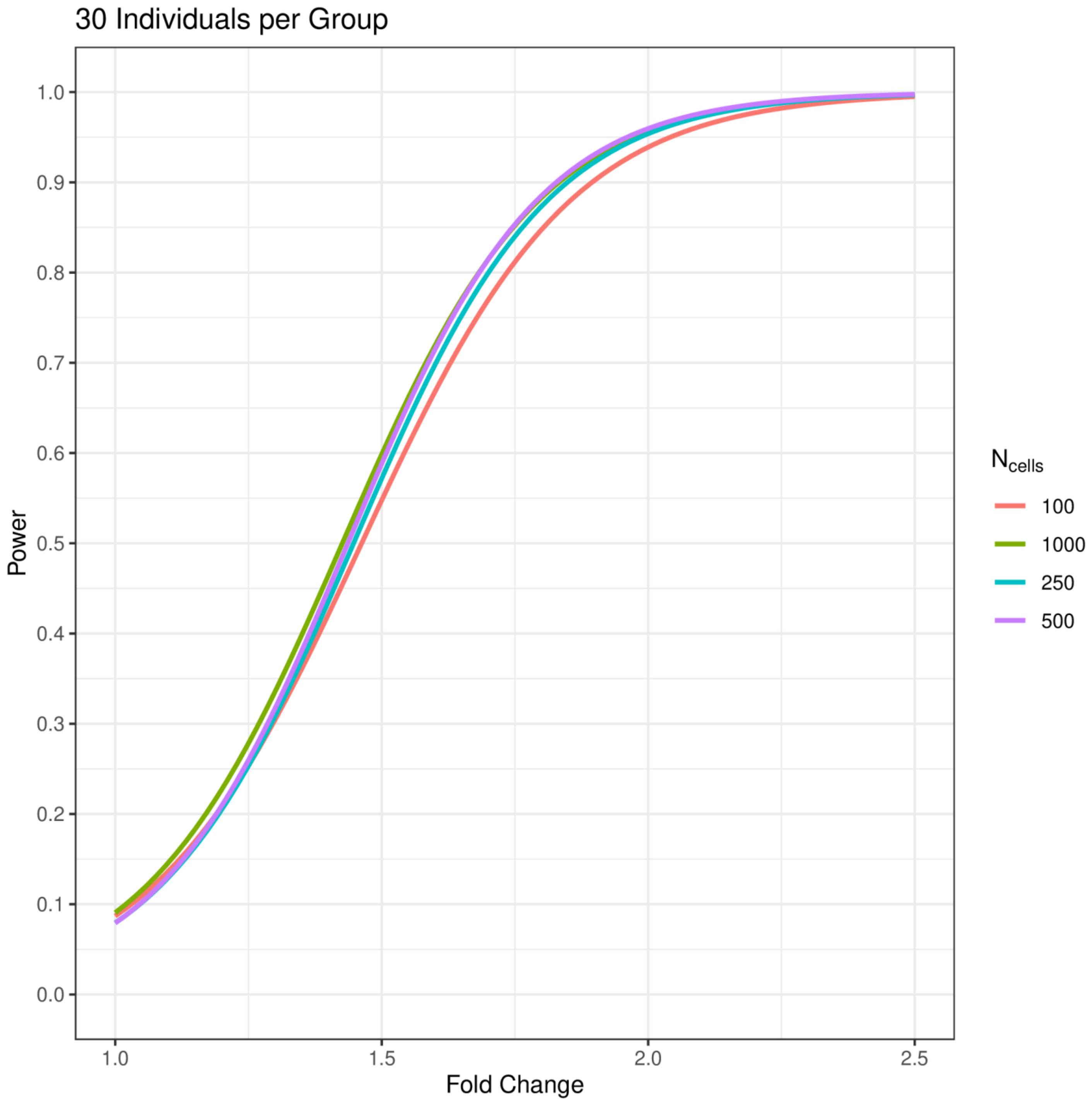
30 Individuals per Group
1.0
0.9
0.8
0.7
0.6
0.5
0.4
0.3
0.2
0.1
0.0
Power
1.0
1.5
2.0
2.5
Fold Change
$N_{cells}$
100
1000
250
500

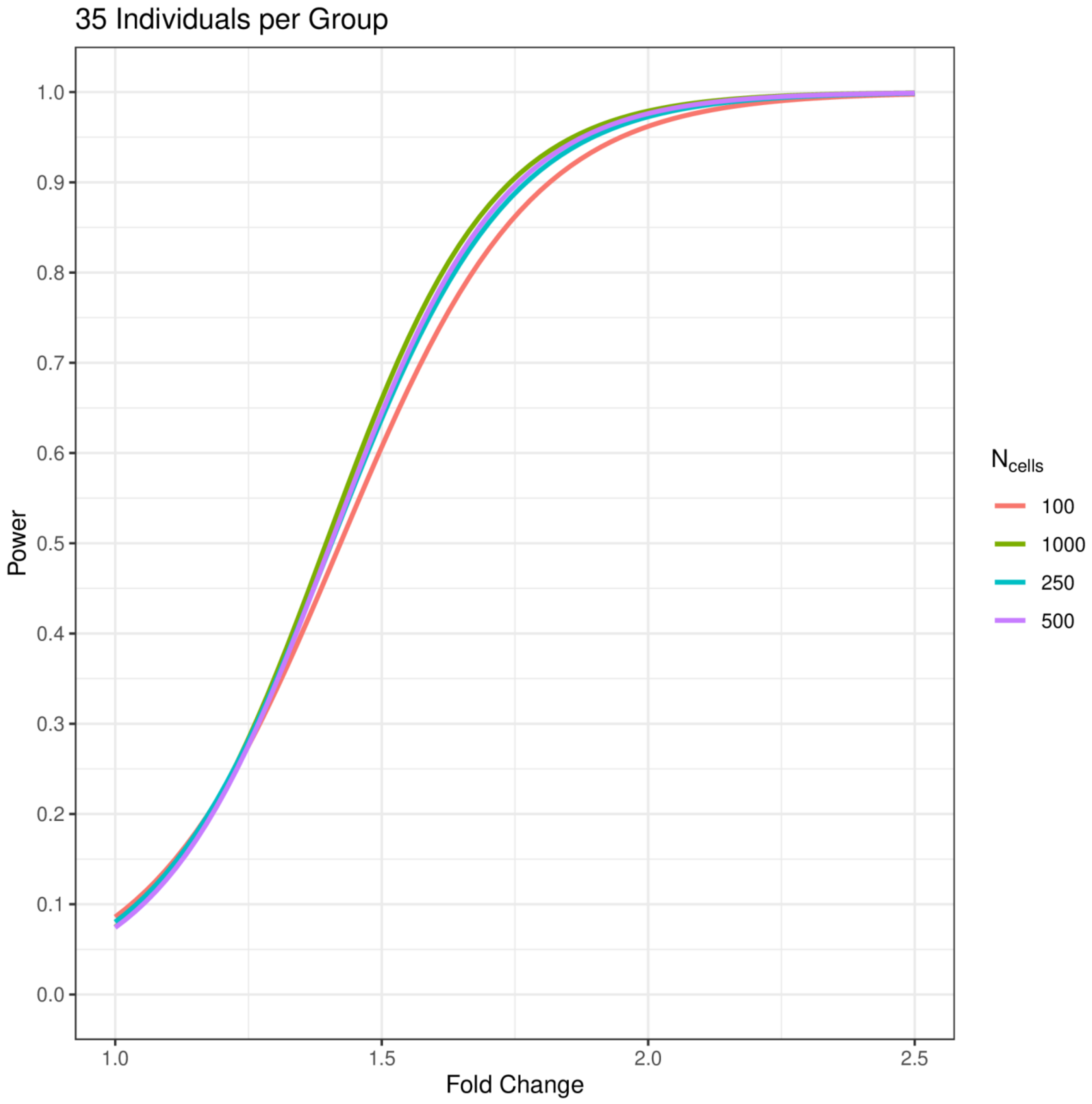
35 Individuals per Group
Power
1.0
0.9
0.8
0.7
0.6
0.5
0.4
0.3
0.2
0.1
0.0
1.0
1.5
2.0
2.5
Fold Change
N cells
100
1000
250
500

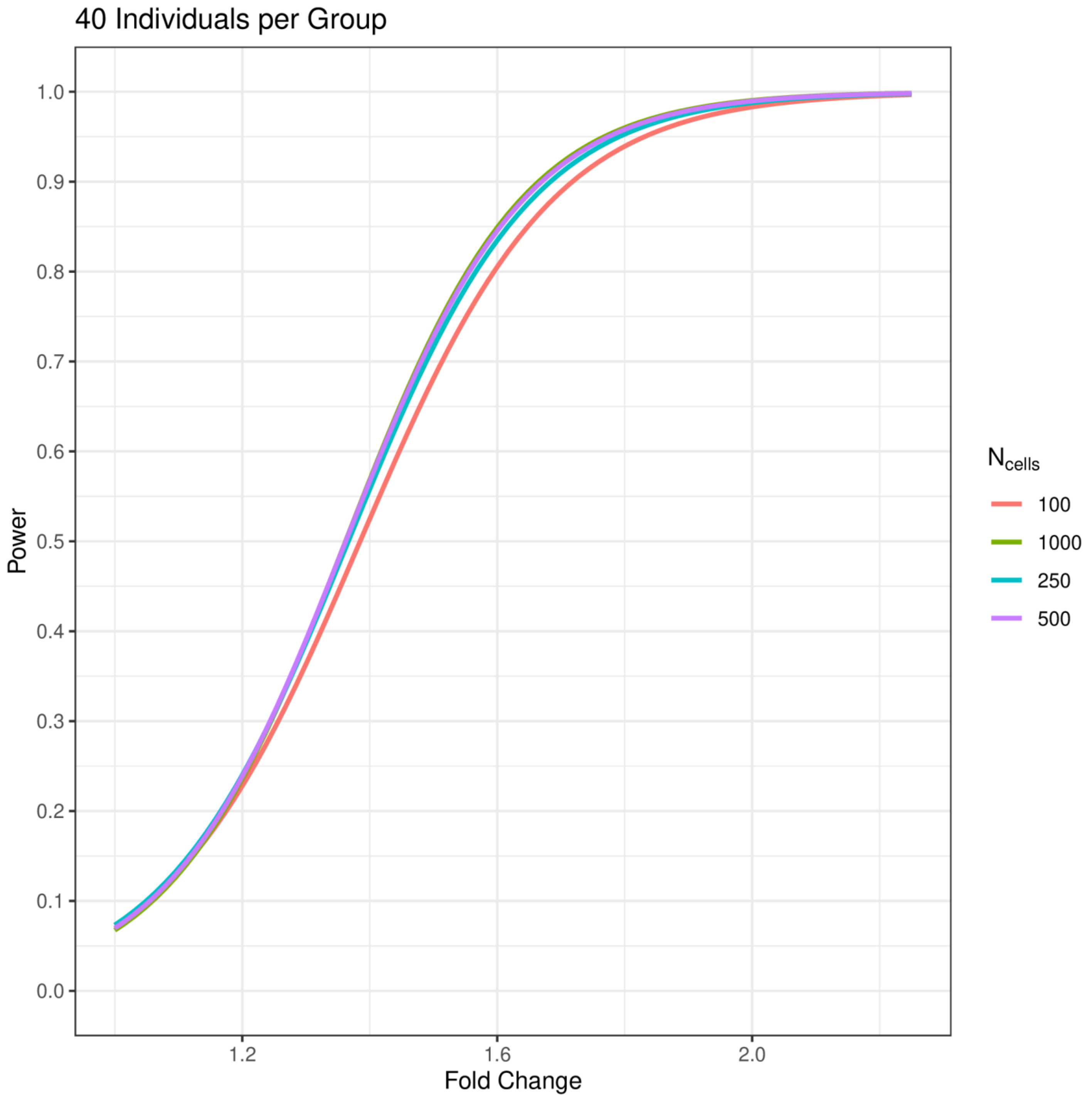
40 Individuals per Group
Power
1.0
0.9
0.8
0.7
0.6
0.5
0.4
0.3
0.2
0.1
0.0
1.2
1.6
2.0
Fold Change
$N_{cells}$
100
1000
250
500

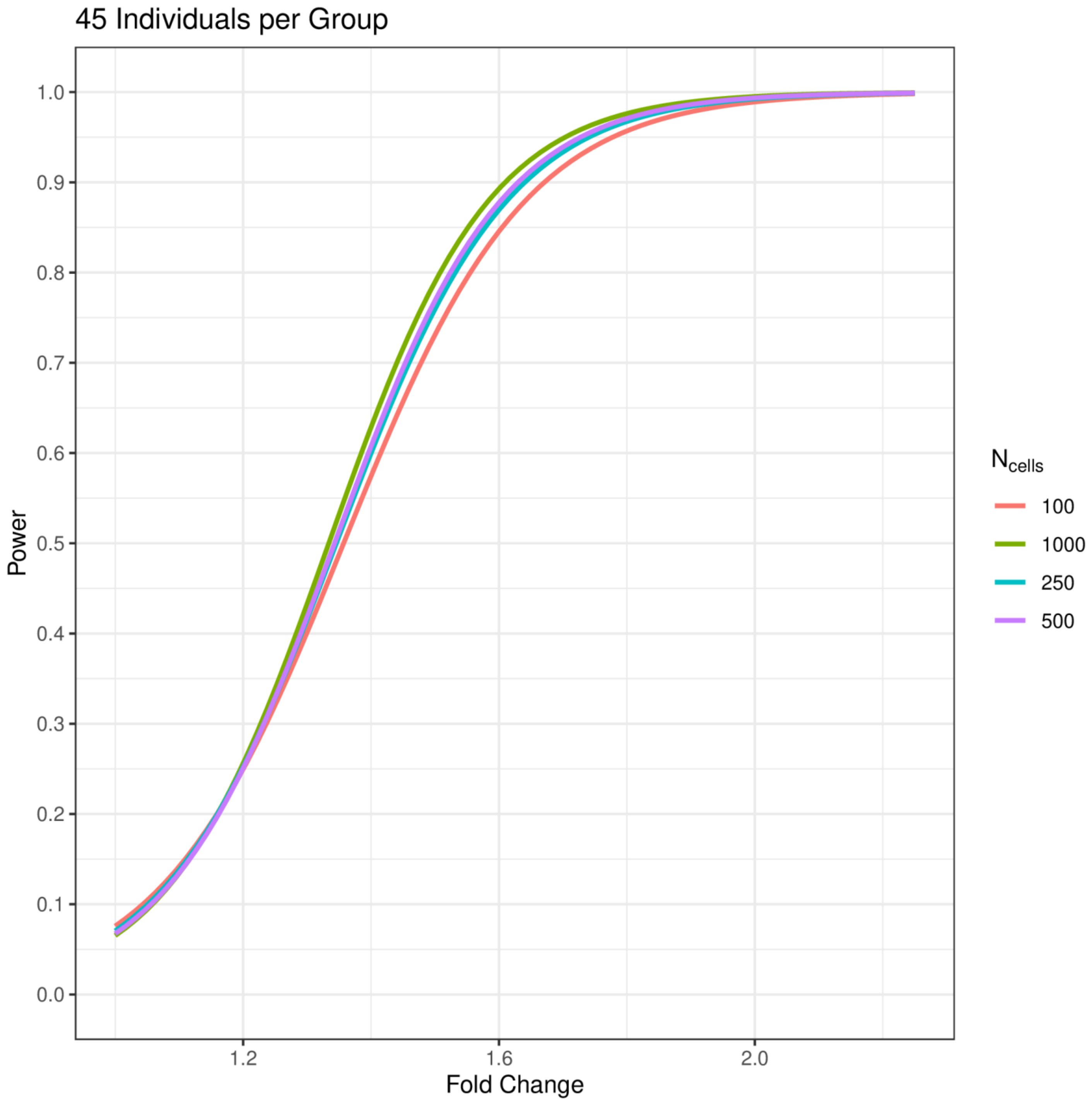
45 Individuals per Group
Power
1.0
0.9
0.8
0.7
0.6
0.5
0.4
0.3
0.2
0.1
0.0
1.2
1.6
2.0
Fold Change
$N_{cells}$
100
1000
250
500

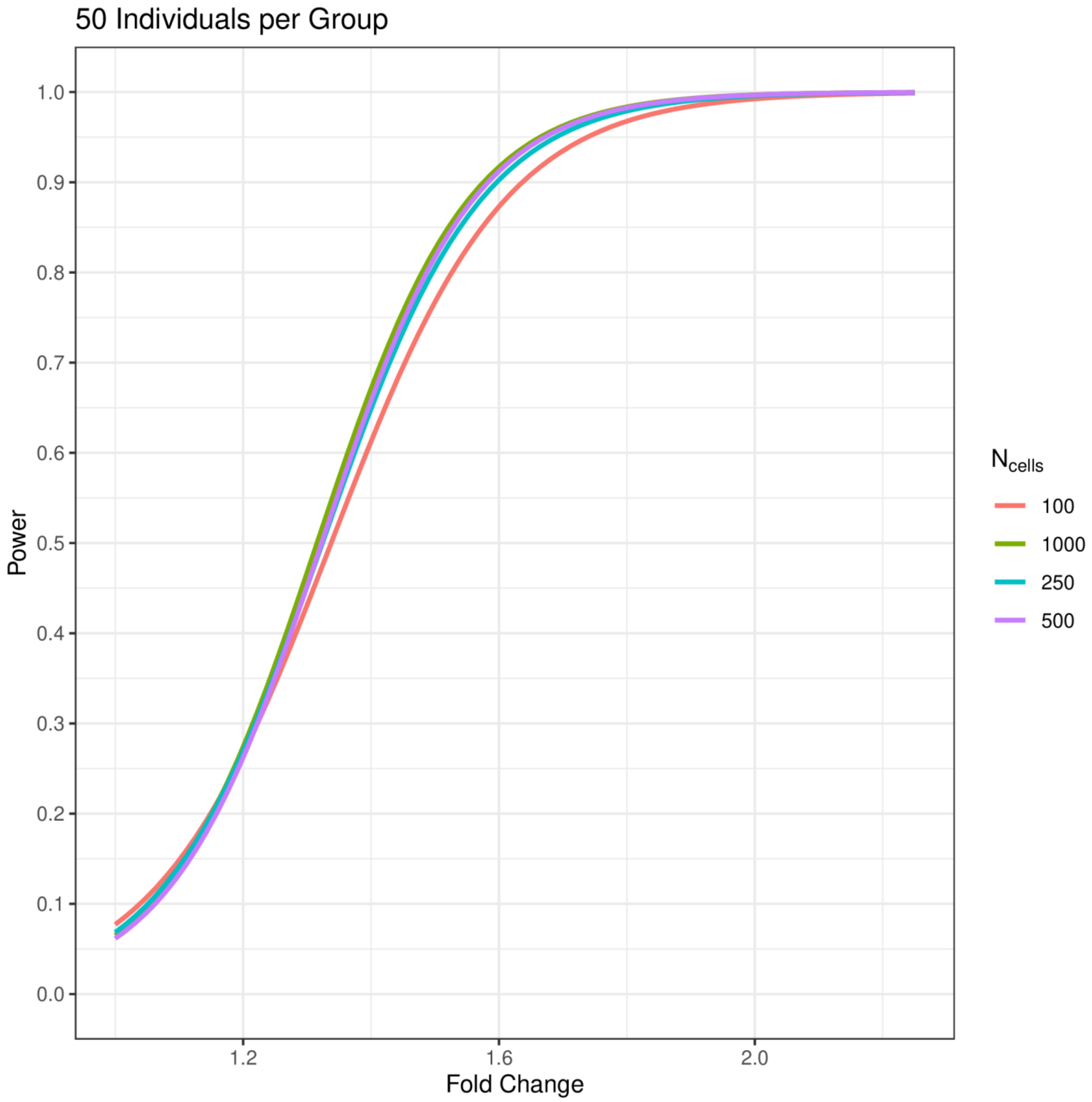
50 Individuals per Group
Power
Fold Change
1.0
0.9
0.8
0.7
0.6
0.5
0.4
0.3
0.2
0.1
0.0
1.2
1.6
2.0
$N_{cells}$
100
1000
250
500

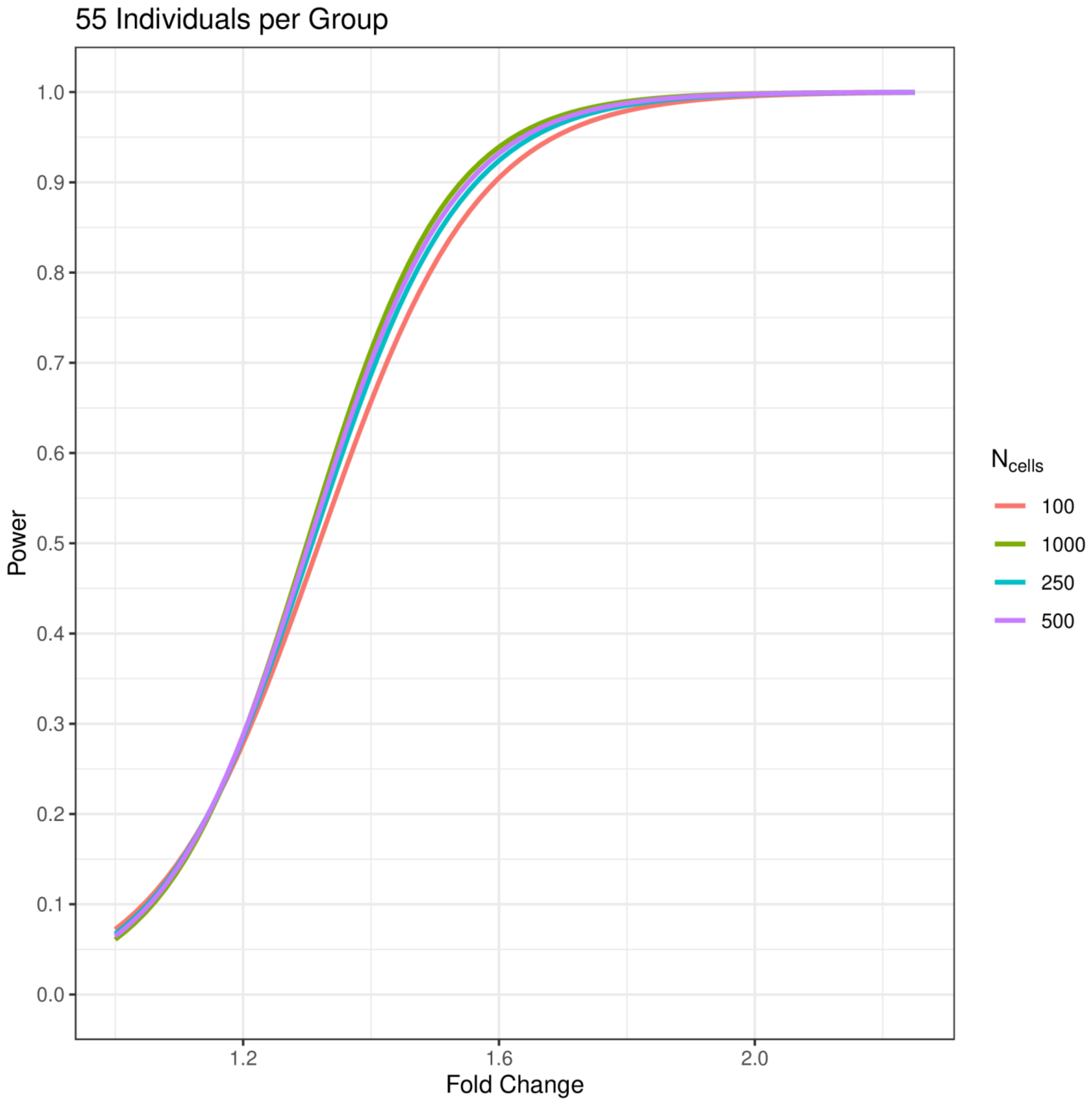
55 Individuals per Group
Power
1.0
0.9
0.8
0.7
0.6
0.5
0.4
0.3
0.2
0.1
0.0
1.2
1.6
2.0
Fold Change
$N_{cells}$
100
1000
250
500

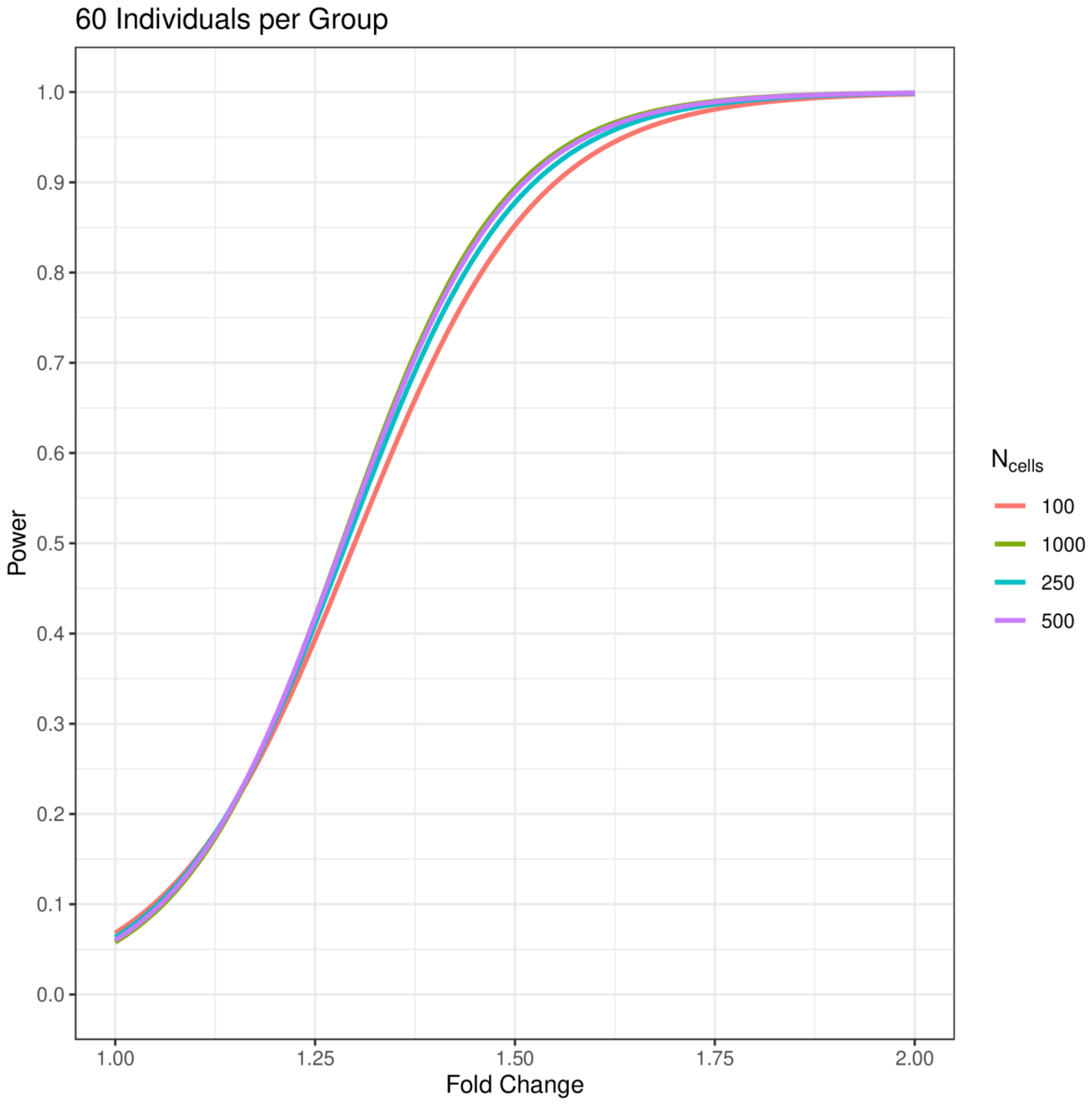
60 Individuals per Group
Power
1.0
0.9
0.8
0.7
0.6
0.5
0.4
0.3
0.2
0.1
0.0
1.00
1.25
1.50
1.75
2.00
Fold Change
$N_{cells}$
100
1000
250
500

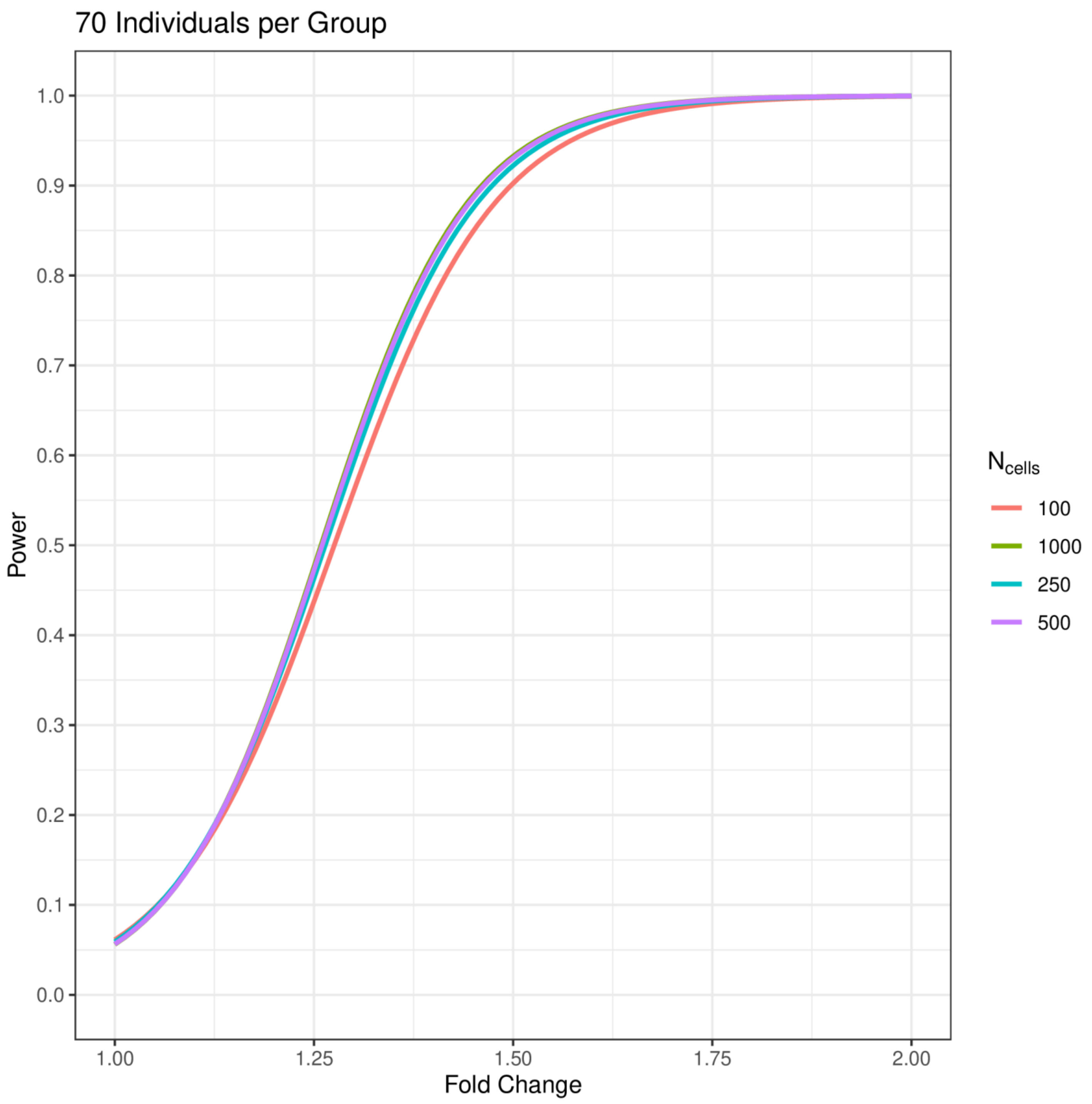
70 Individuals per Group
Power
1.0
0.9
0.8
0.7
0.6
0.5
0.4
0.3
0.2
0.1
0.0
1.00
1.25
1.50
1.75
2.00
Fold Change
$N_{cells}$
100
1000
250
500

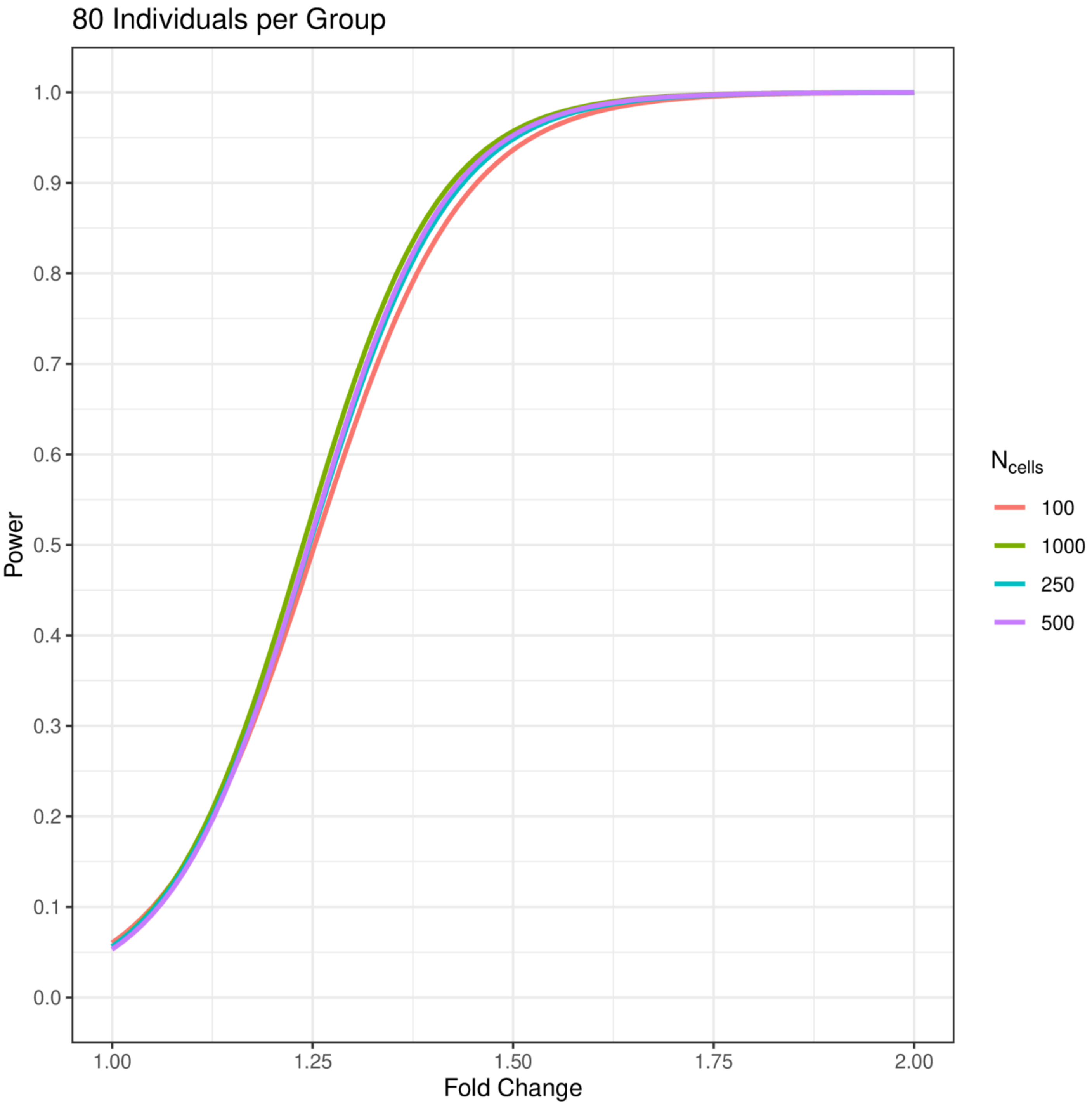

80 Individuals per Group
Power
1.0
0.9
0.8
0.7
0.6
0.5
0.4
0.3
0.2
0.1
0.0
1.00
1.25
1.50
1.75
2.00
Fold Change
Ncells
100
1000
250
500

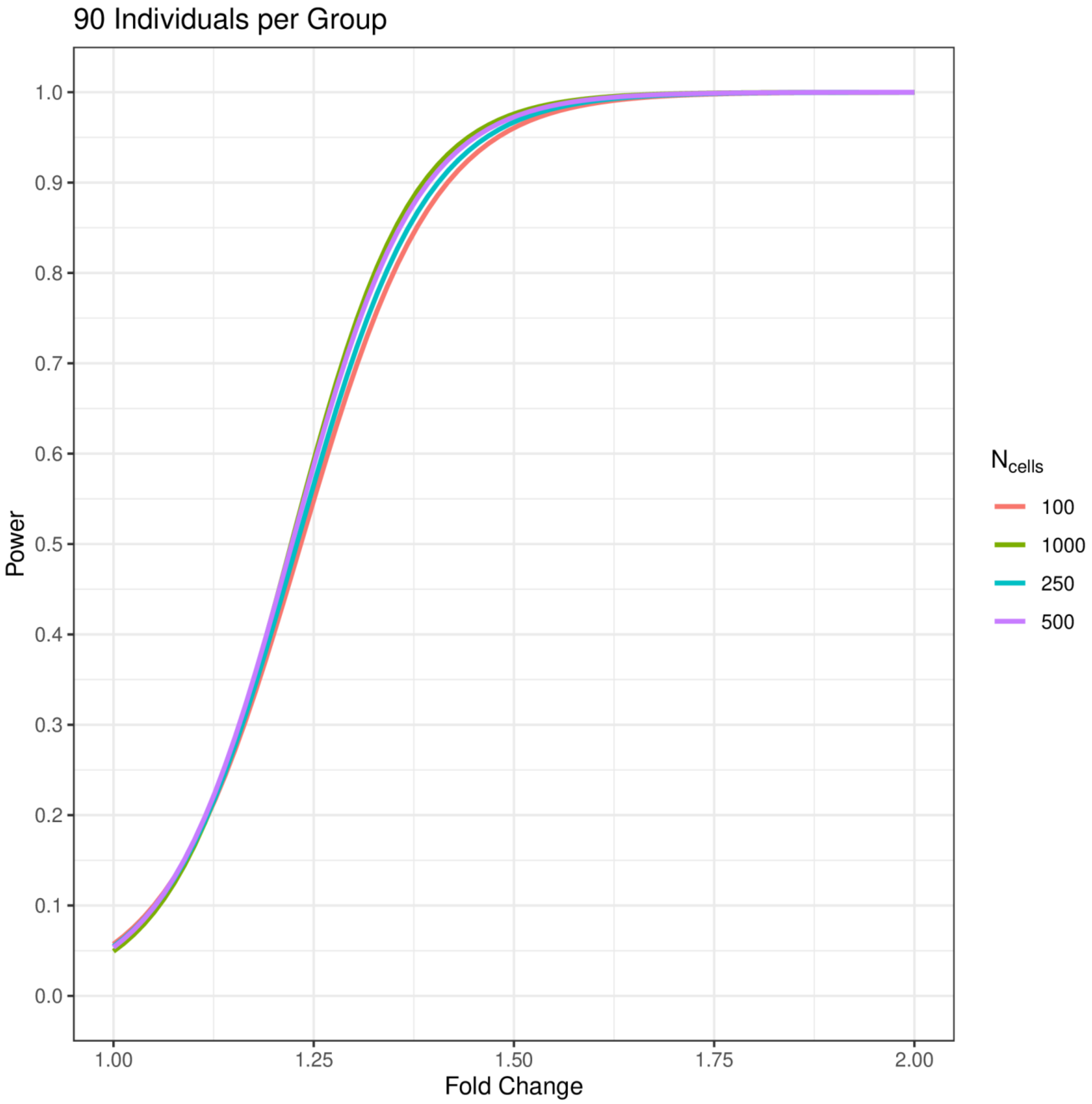
90 Individuals per Group
Power
Fold Change
1.0
0.9
0.8
0.7
0.6
0.5
0.4
0.3
0.2
0.1
0.0
1.00
1.25
1.50
1.75
2.00
N cells
100
1000
250
500

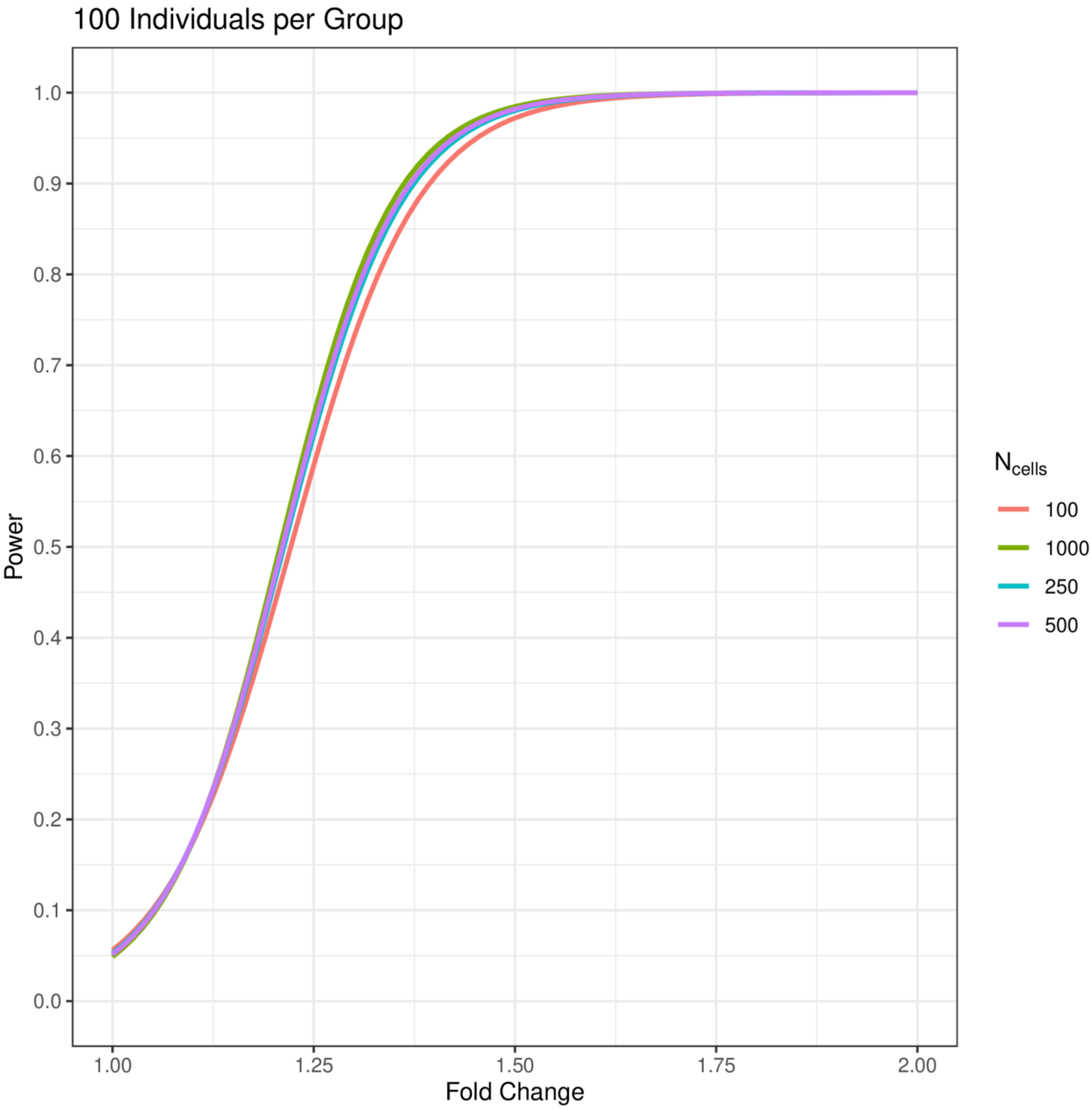

100 Individuals per Group
1.0
0.9
0.8
0.7
0.6
0.5
0.4
0.3
0.2
0.1
0.0
Power
1.00
1.25
1.50
1.75
2.00
Fold Change
$N_{cells}$
100
1000
250
500

**Fig. S6. | Power calculations using MAST with a random effect for individual.** Power curves for MAST using a random effect to account for intra-individual correlation. Curves are computed for 100, 250, 500, and 1,000 cells per individual using an $\alpha = 0.01$. The number of individuals per group ranges from 3 to 100 and is listed above each plot.

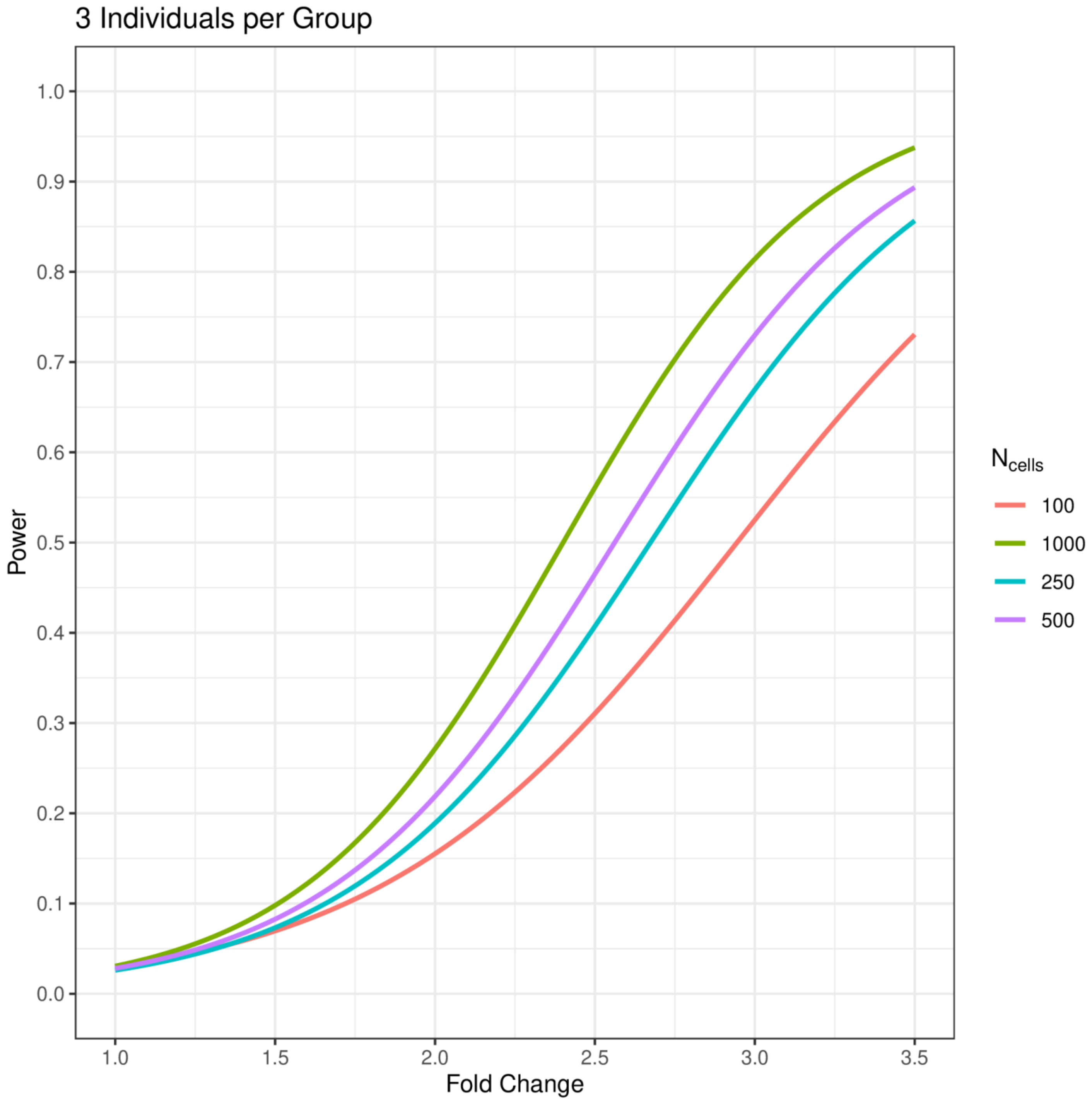

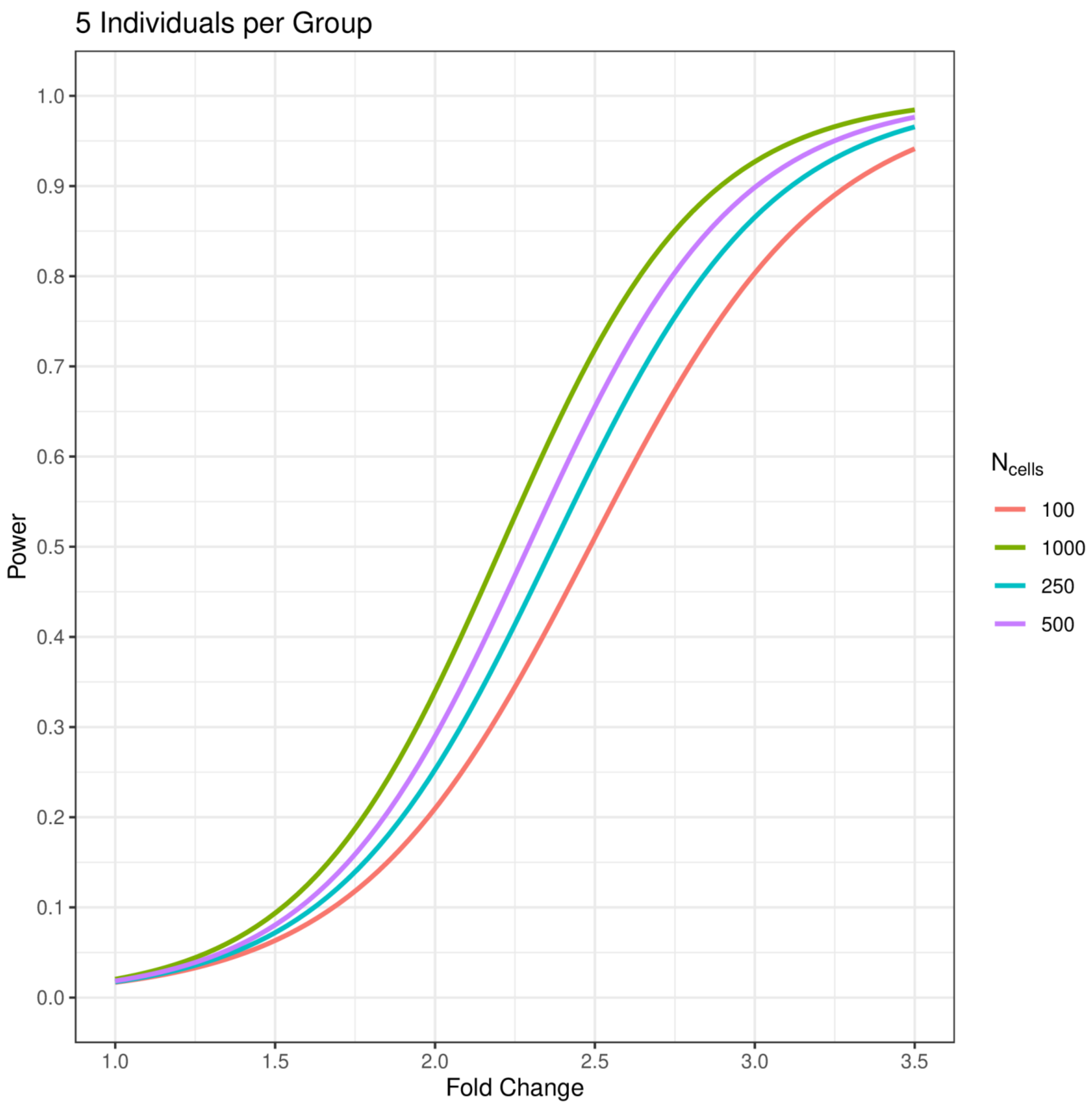

5 Individuals per Group
Power
Fold Change
1.0
0.9
0.8
0.7
0.6
0.5
0.4
0.3
0.2
0.1
0.0
1.0
1.5
2.0
2.5
3.0
3.5
N cells
100
1000
250
500

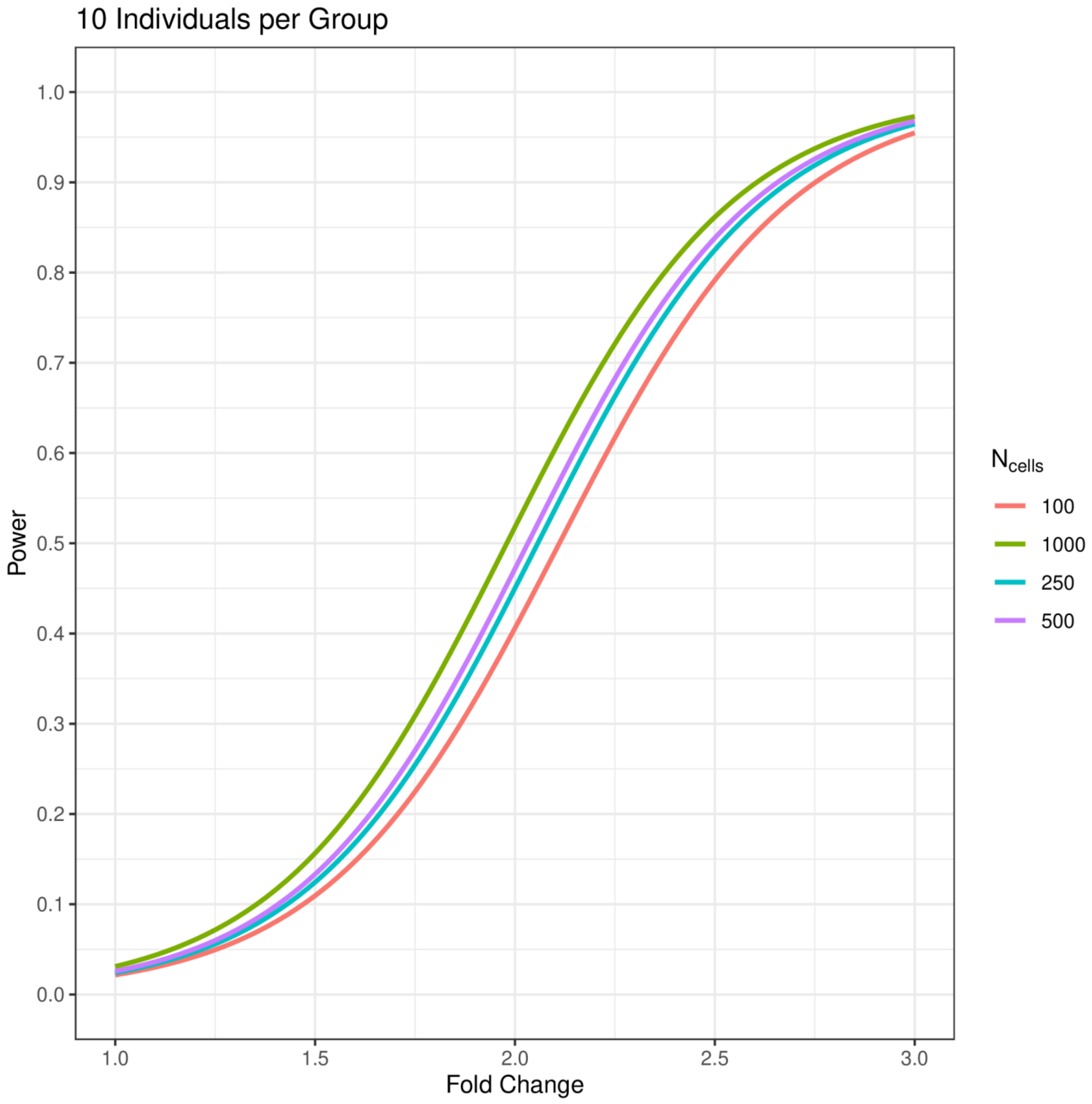

10 Individuals per Group
Power
1.0
0.9
0.8
0.7
0.6
0.5
0.4
0.3
0.2
0.1
0.0
1.0
1.5
2.0
2.5
3.0
Fold Change
$N_{cells}$
100
1000
250
500

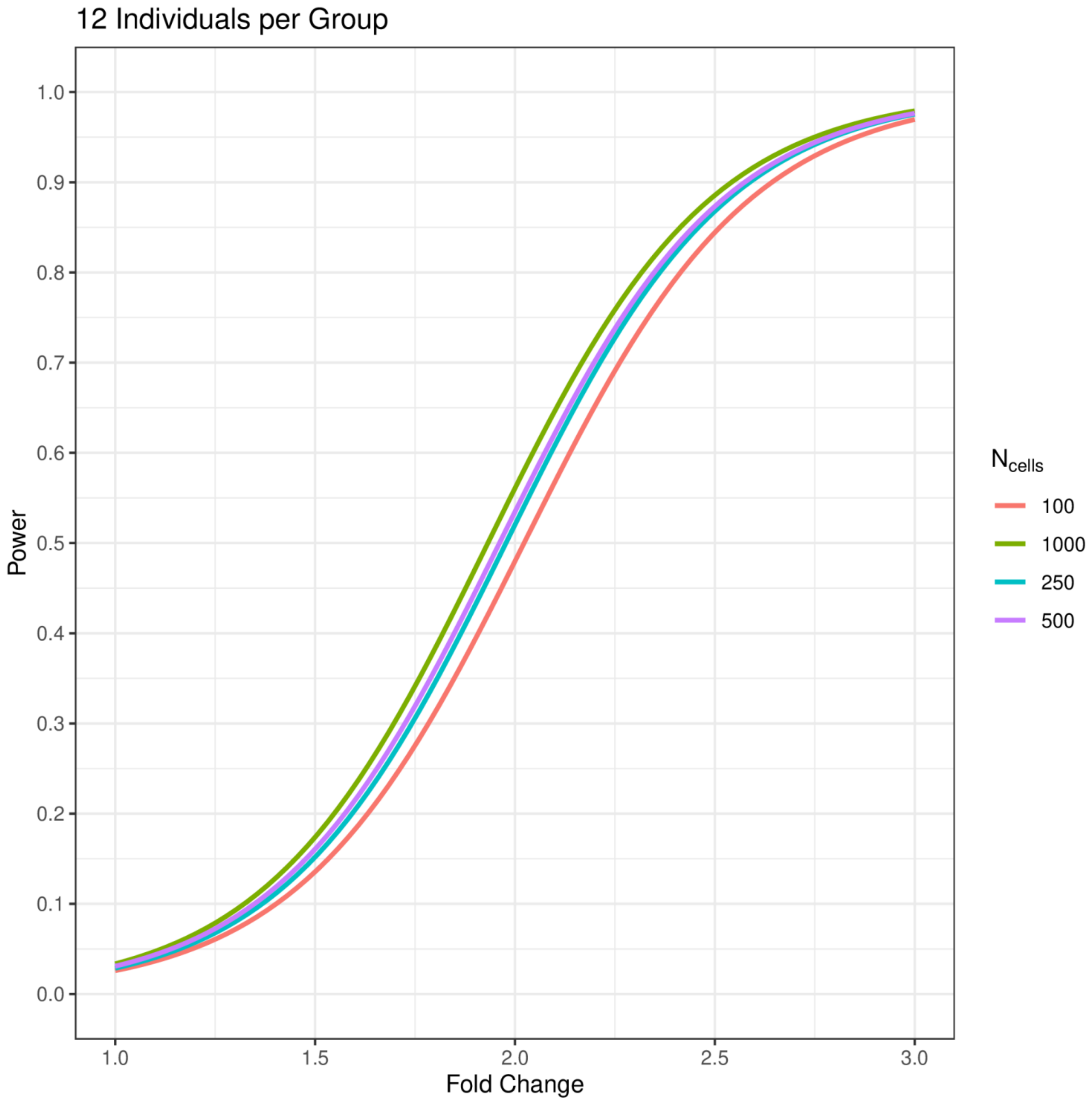
12 Individuals per Group
Power
Fold Change
1.0
0.9
0.8
0.7
0.6
0.5
0.4
0.3
0.2
0.1
0.0
1.0
1.5
2.0
2.5
3.0
$N_{cells}$
100
1000
250
500

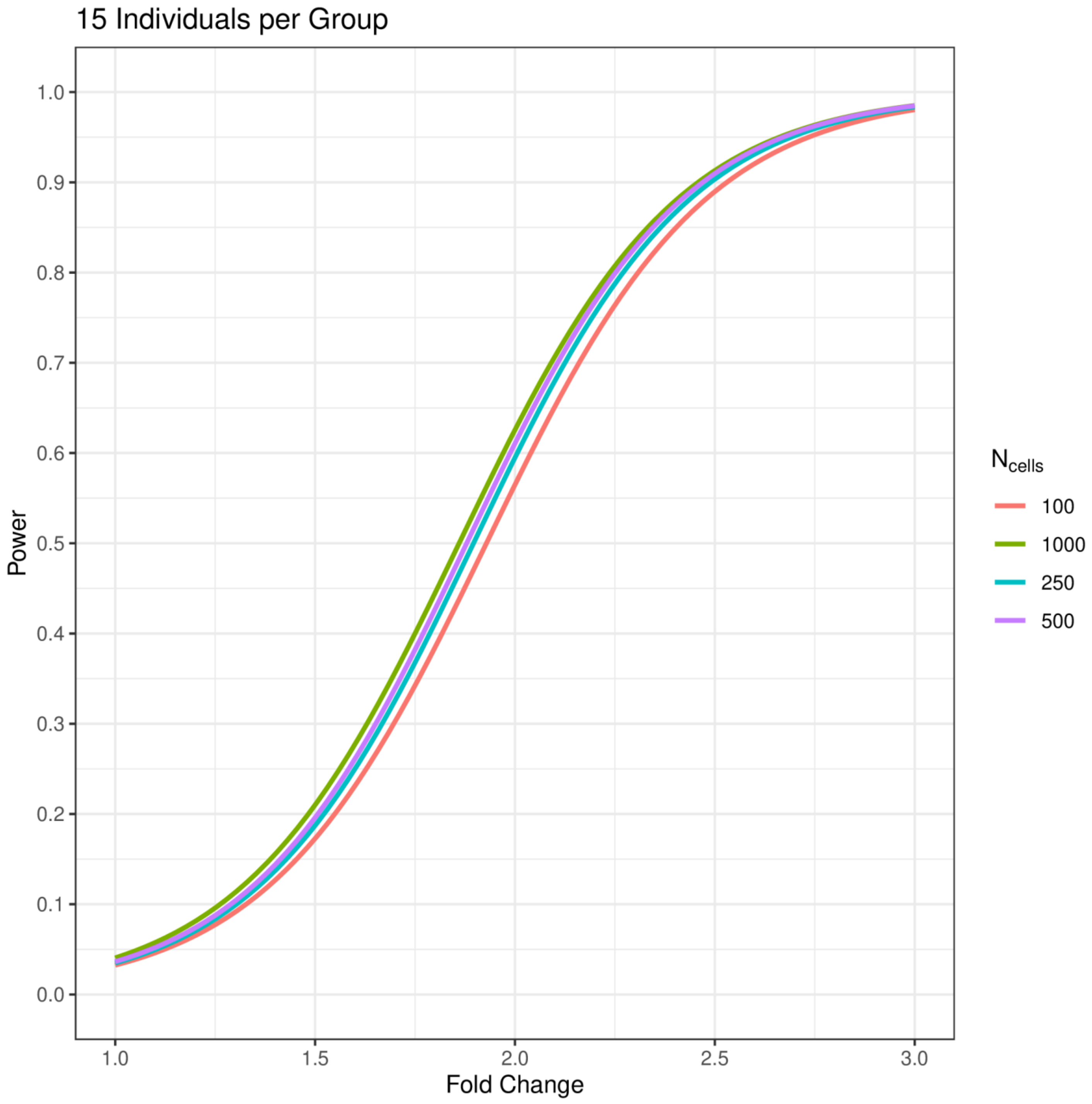
15 Individuals per Group
Power
Fold Change
1.0
0.9
0.8
0.7
0.6
0.5
0.4
0.3
0.2
0.1
0.0
1.0
1.5
2.0
2.5
3.0
$N_{cells}$
100
1000
250
500

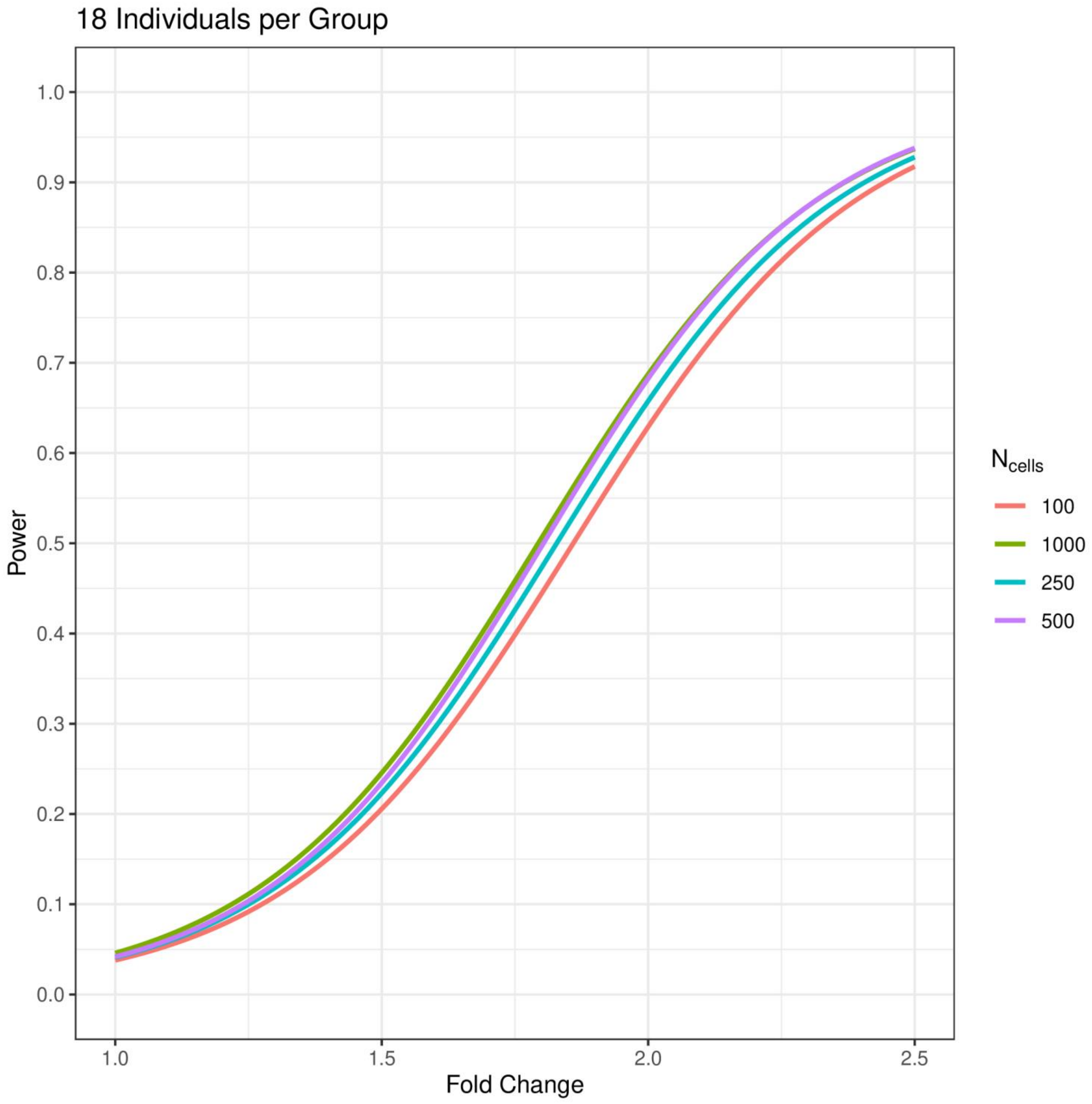

18 Individuals per Group
1.0
0.9
0.8
0.7
0.6
0.5
0.4
0.3
0.2
0.1
0.0
Power
1.0
1.5
2.0
2.5
Fold Change
$N_{cells}$
100
1000
250
500

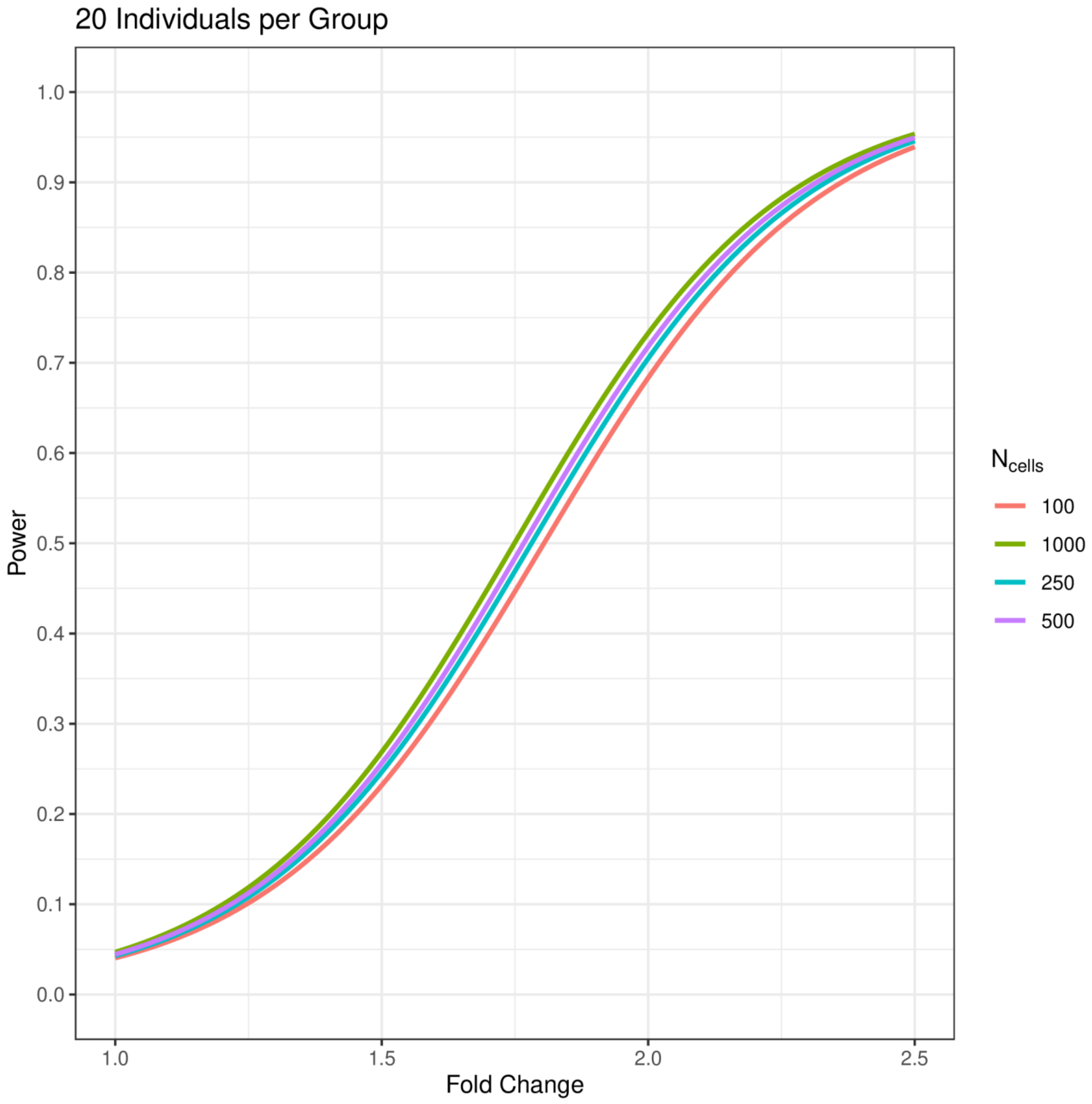
20 Individuals per Group
Power
1.0
0.9
0.8
0.7
0.6
0.5
0.4
0.3
0.2
0.1
0.0
1.0
1.5
2.0
2.5
Fold Change
$N_{cells}$
100
1000
250
500

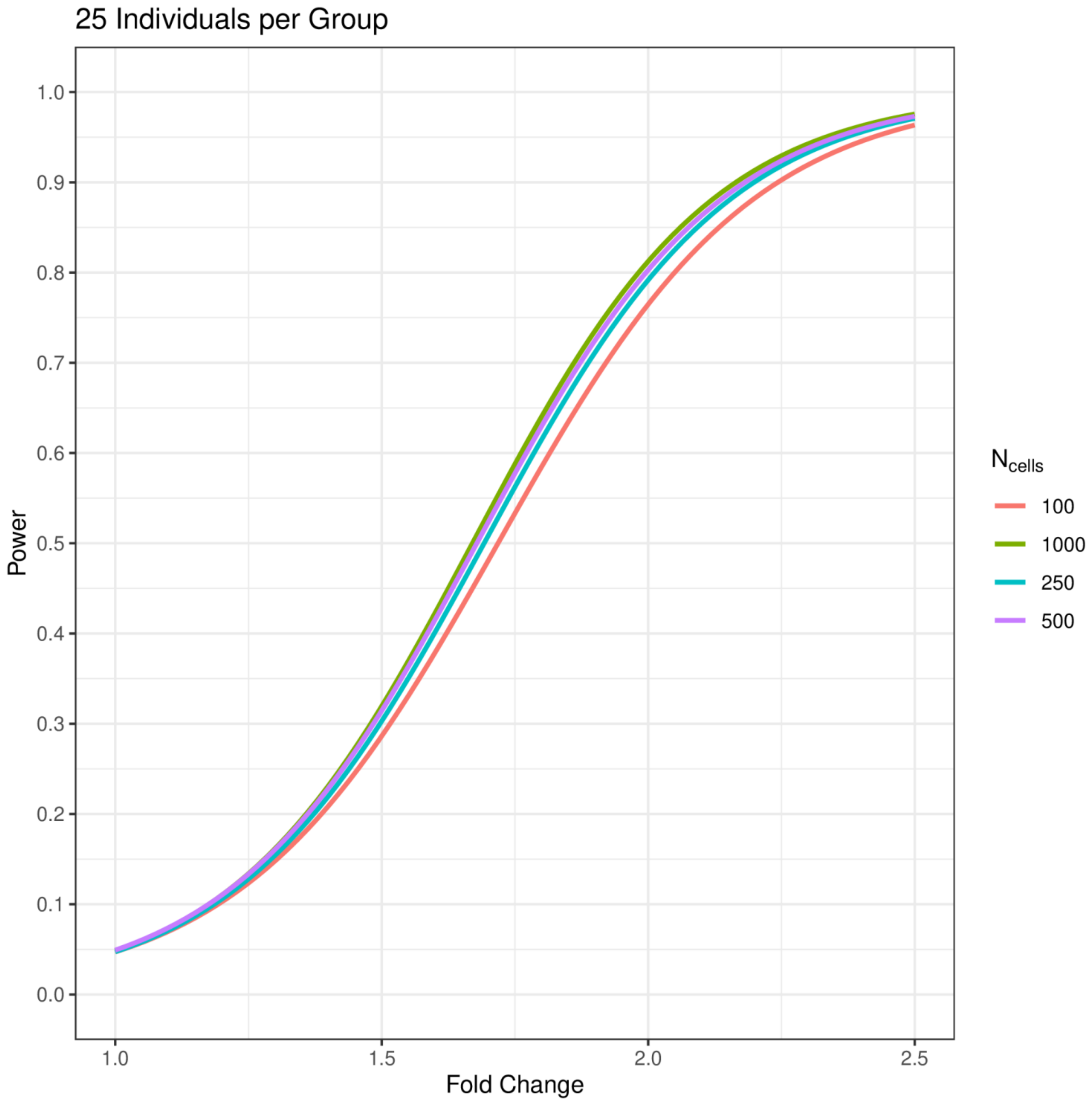

25 Individuals per Group
Power
Fold Change
1.0
0.9
0.8
0.7
0.6
0.5
0.4
0.3
0.2
0.1
0.0
1.0
1.5
2.0
2.5
N cells
100
1000
250
500

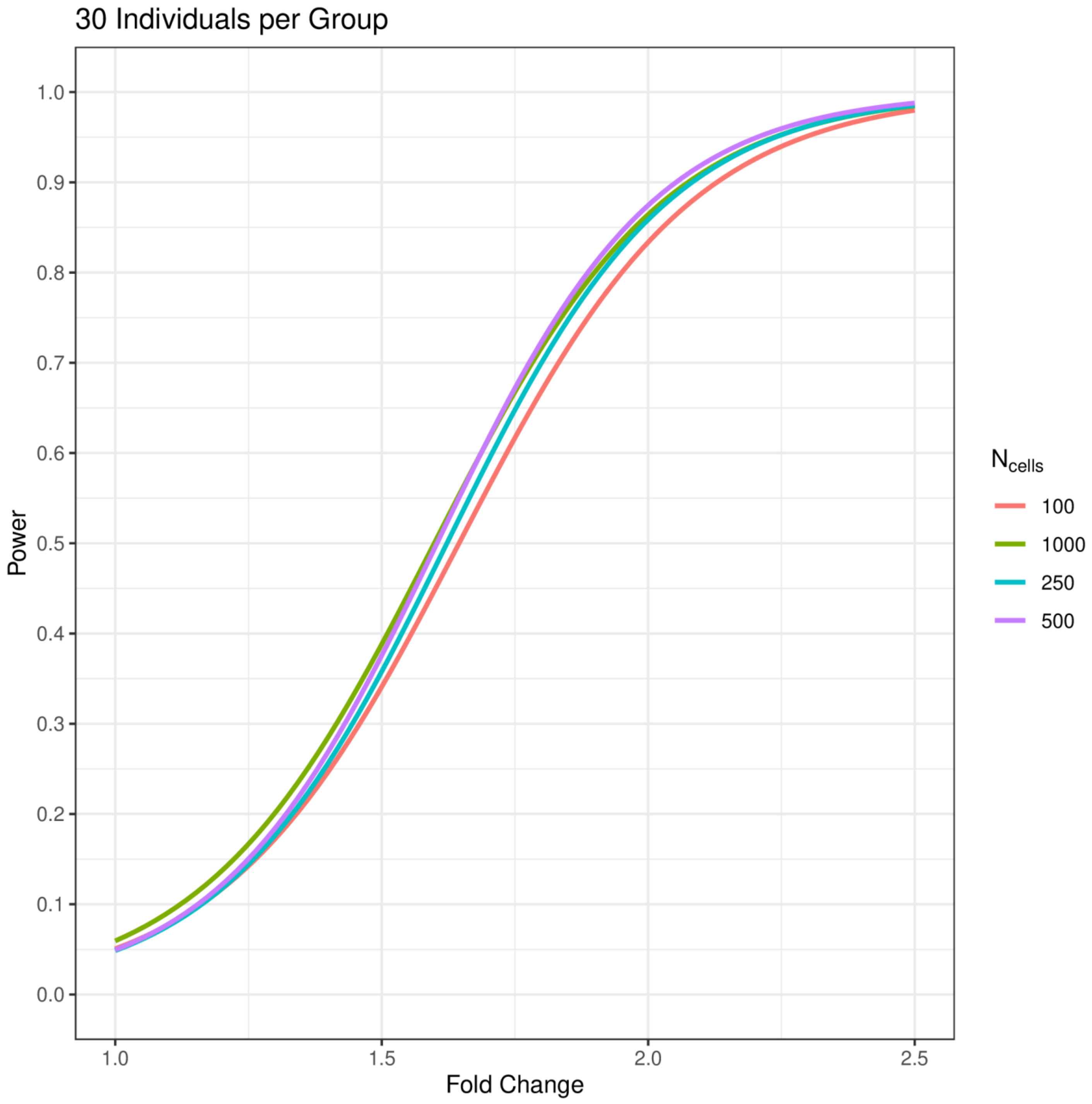
30 Individuals per Group
Power
1.0
0.9
0.8
0.7
0.6
0.5
0.4
0.3
0.2
0.1
0.0
1.0
1.5
2.0
2.5
Fold Change
$N_{cells}$
100
1000
250
500

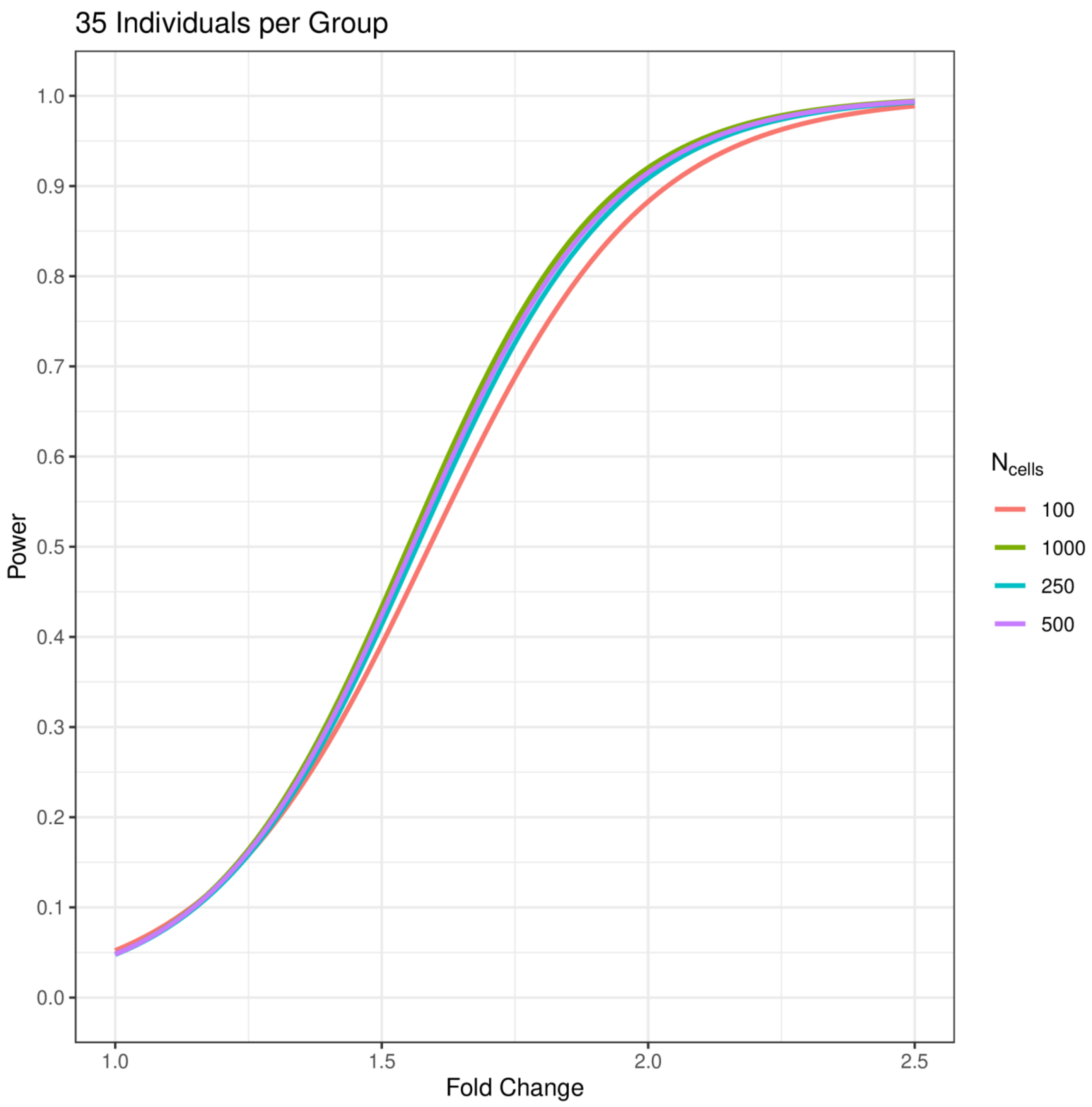
35 Individuals per Group
Power
Fold Change
1.0
0.9
0.8
0.7
0.6
0.5
0.4
0.3
0.2
0.1
0.0
1.0
1.5
2.0
2.5
$N_{cells}$
100
1000
250
500

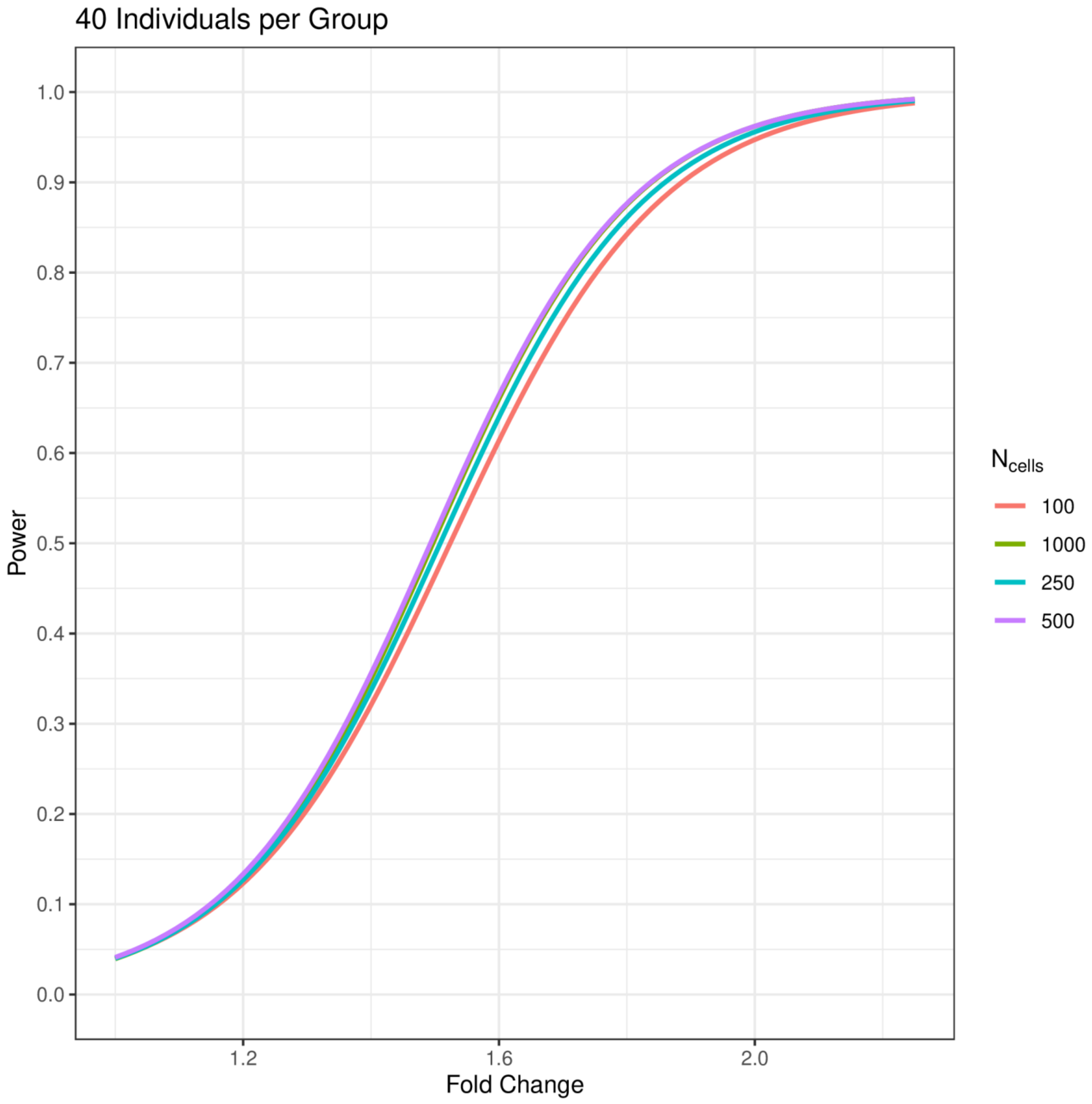
40 Individuals per Group
Power
1.0
0.9
0.8
0.7
0.6
0.5
0.4
0.3
0.2
0.1
0.0
1.2
1.6
2.0
Fold Change
$N_{cells}$
100
1000
250
500

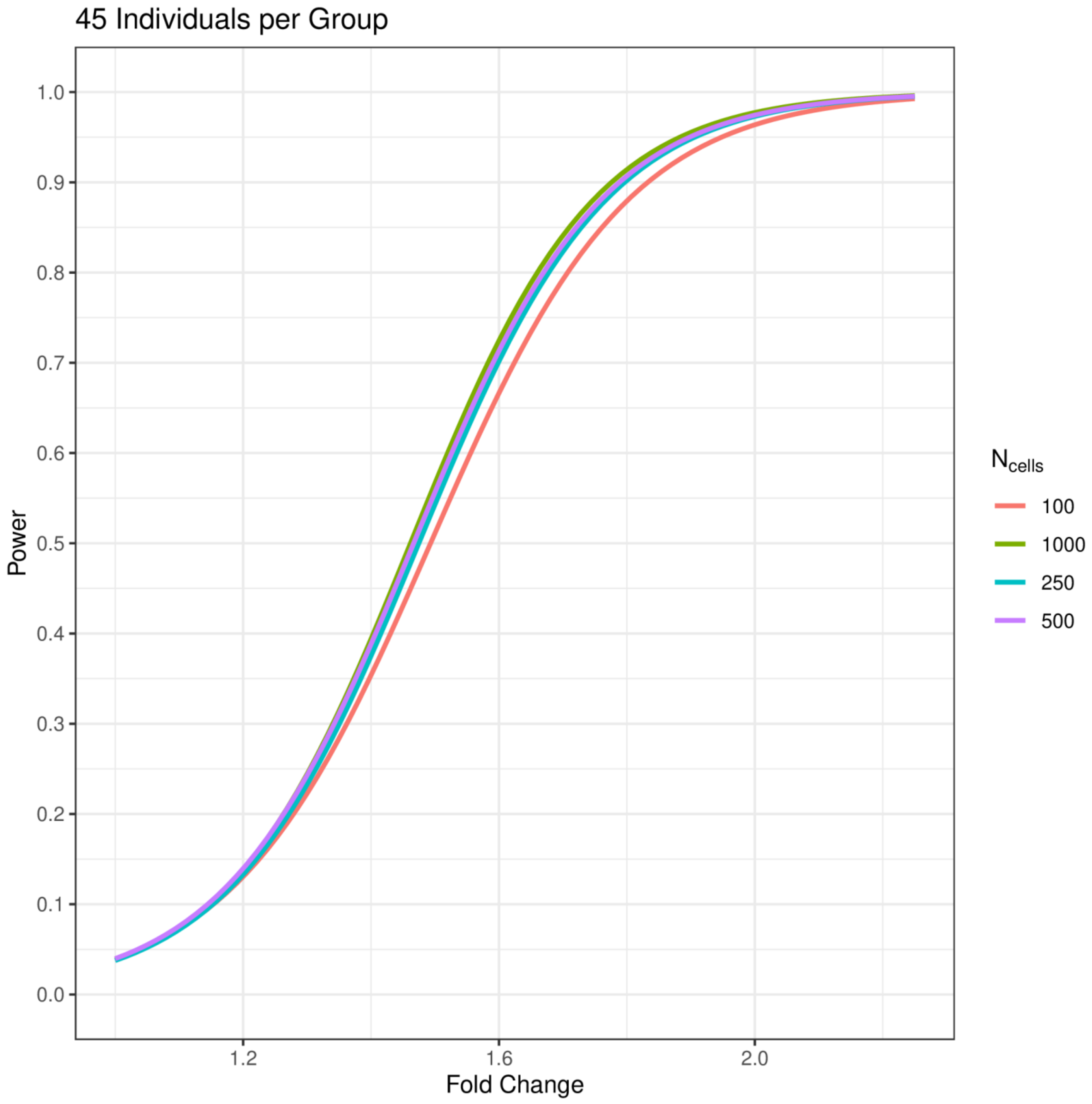
45 Individuals per Group
Power
Fold Change
1.0
0.9
0.8
0.7
0.6
0.5
0.4
0.3
0.2
0.1
0.0
1.2
1.6
2.0
$N_{cells}$
100
1000
250
500

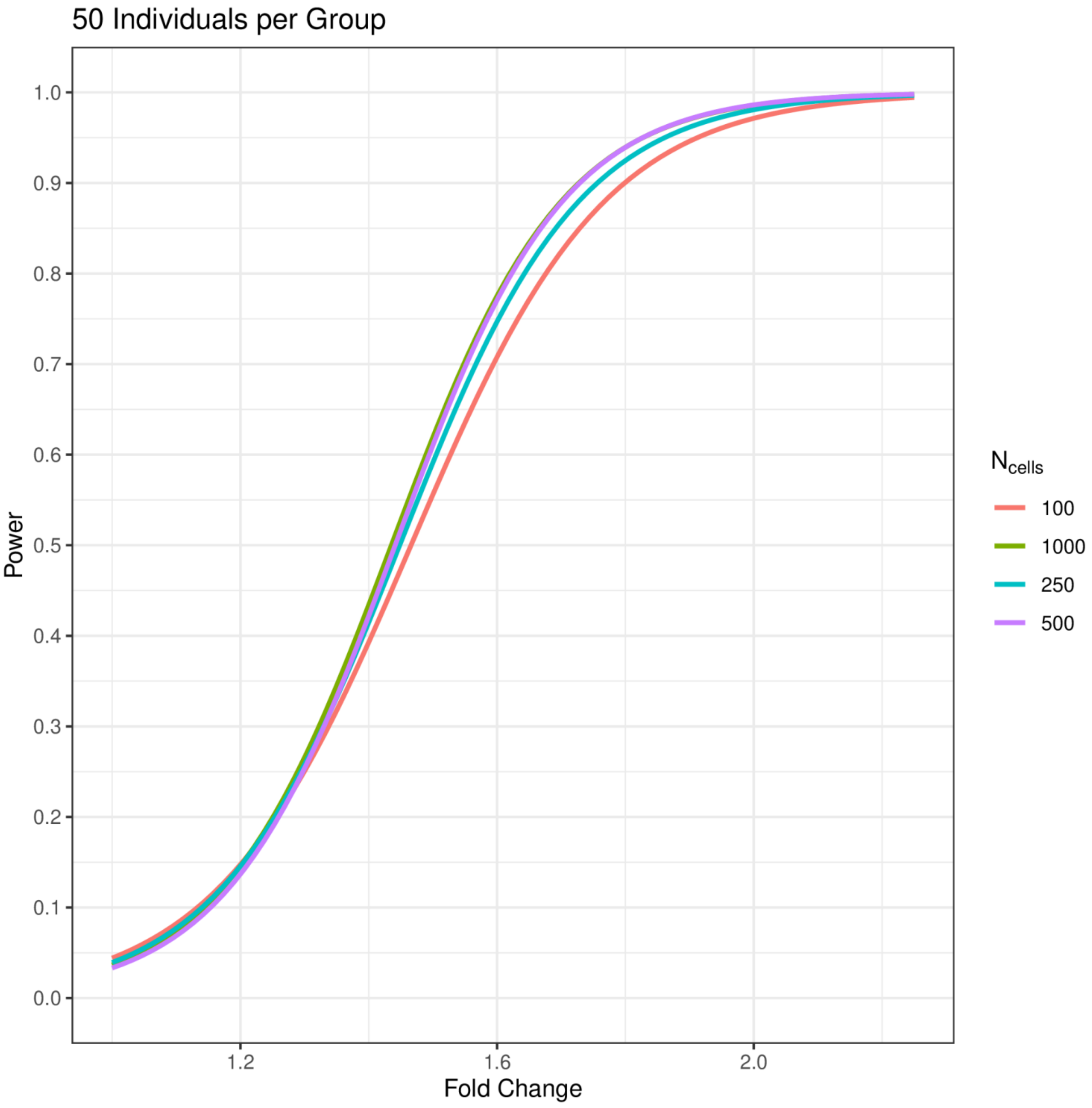

50 Individuals per Group
1.0
0.9
0.8
0.7
0.6
0.5
0.4
0.3
0.2
0.1
0.0
Power
1.2
1.6
2.0
Fold Change
$N_{cells}$
100
1000
250
500

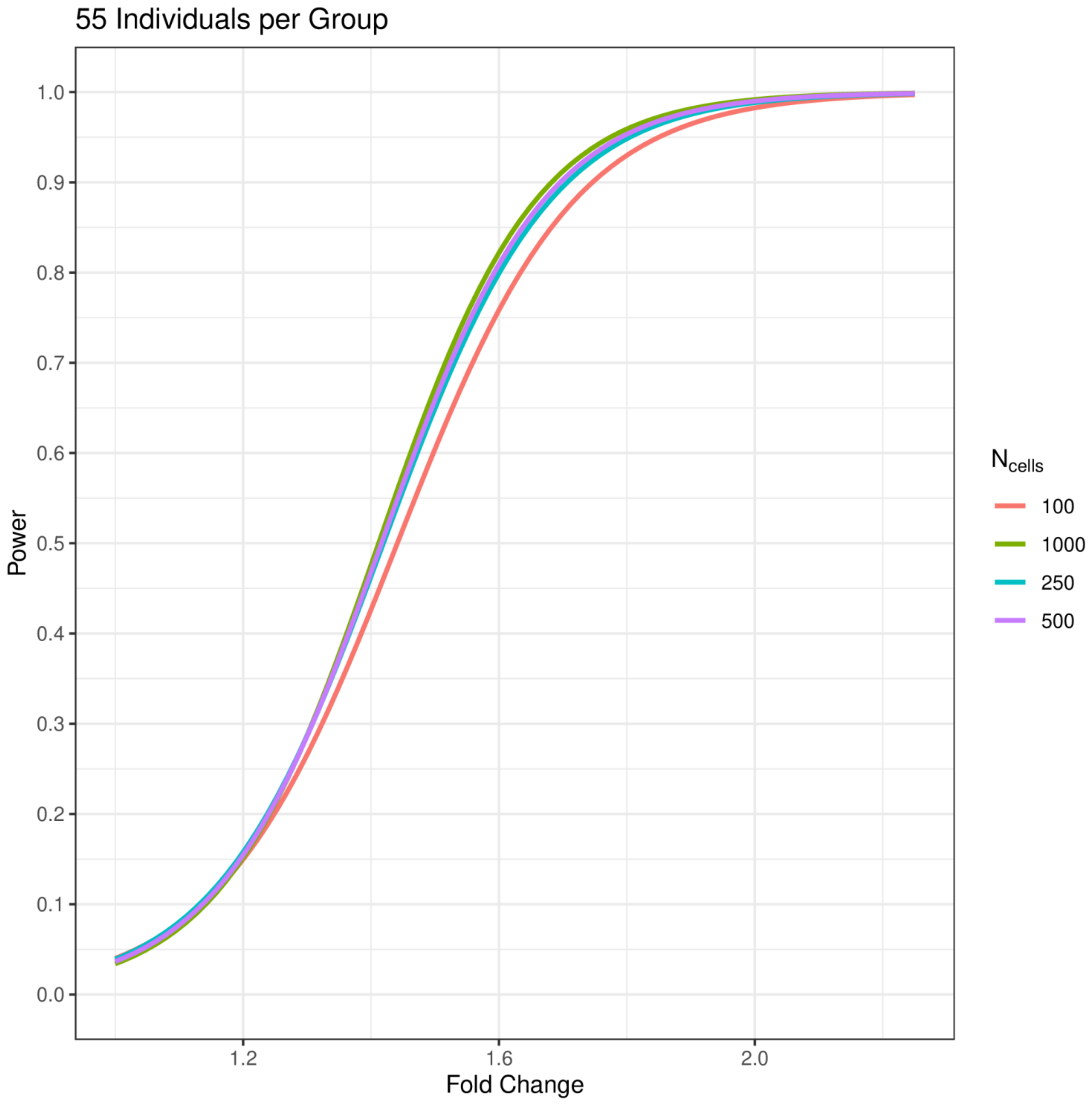

55 Individuals per Group
1.0
0.9
0.8
0.7
0.6
0.5
0.4
0.3
0.2
0.1
0.0
Power
1.2
1.6
2.0
Fold Change
$N_{cells}$
100
1000
250
500


115

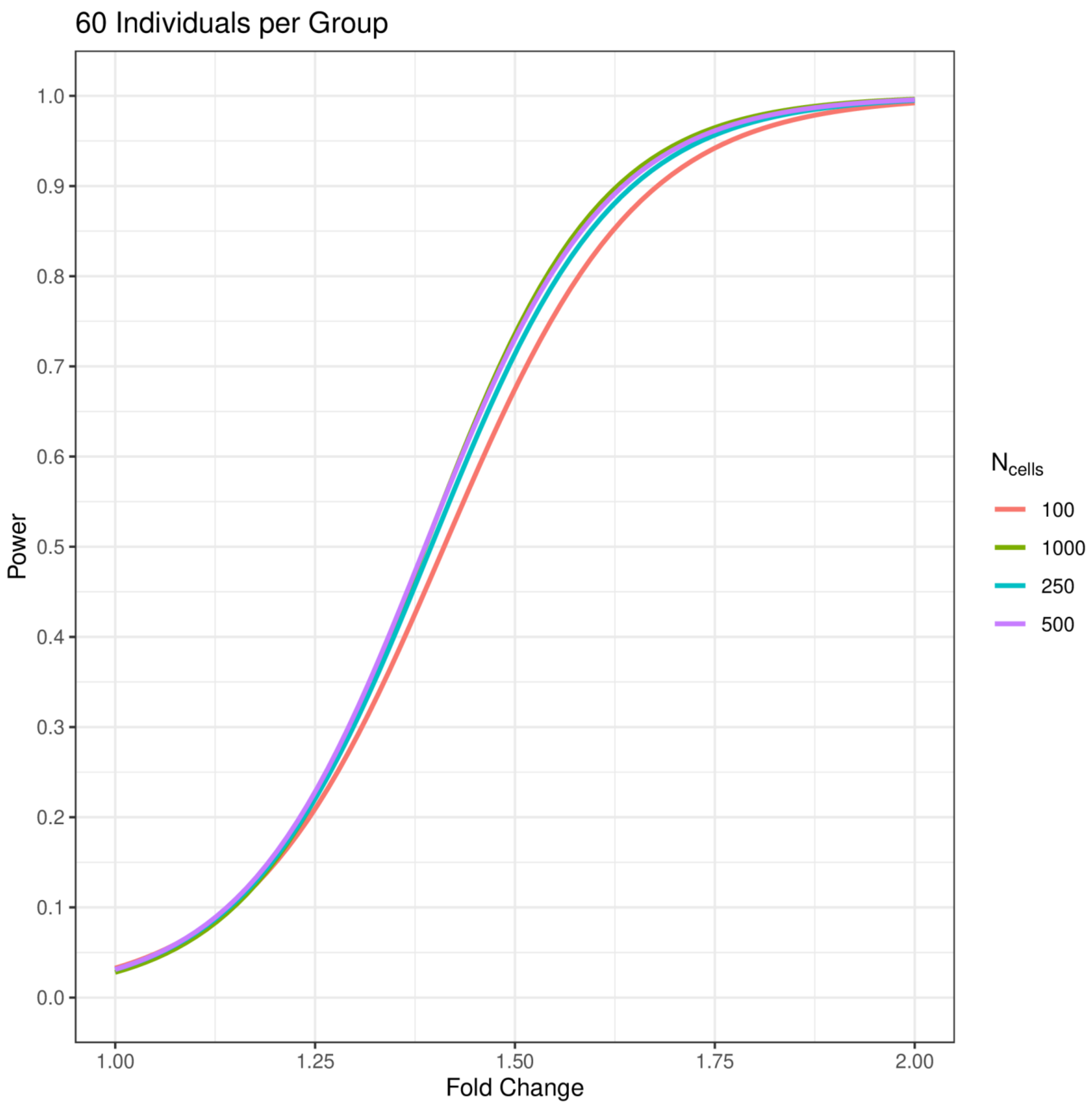
60 Individuals per Group
Power
1.0
0.9
0.8
0.7
0.6
0.5
0.4
0.3
0.2
0.1
0.0
1.00
1.25
1.50
1.75
2.00
Fold Change
$N_{cells}$
100
1000
250
500

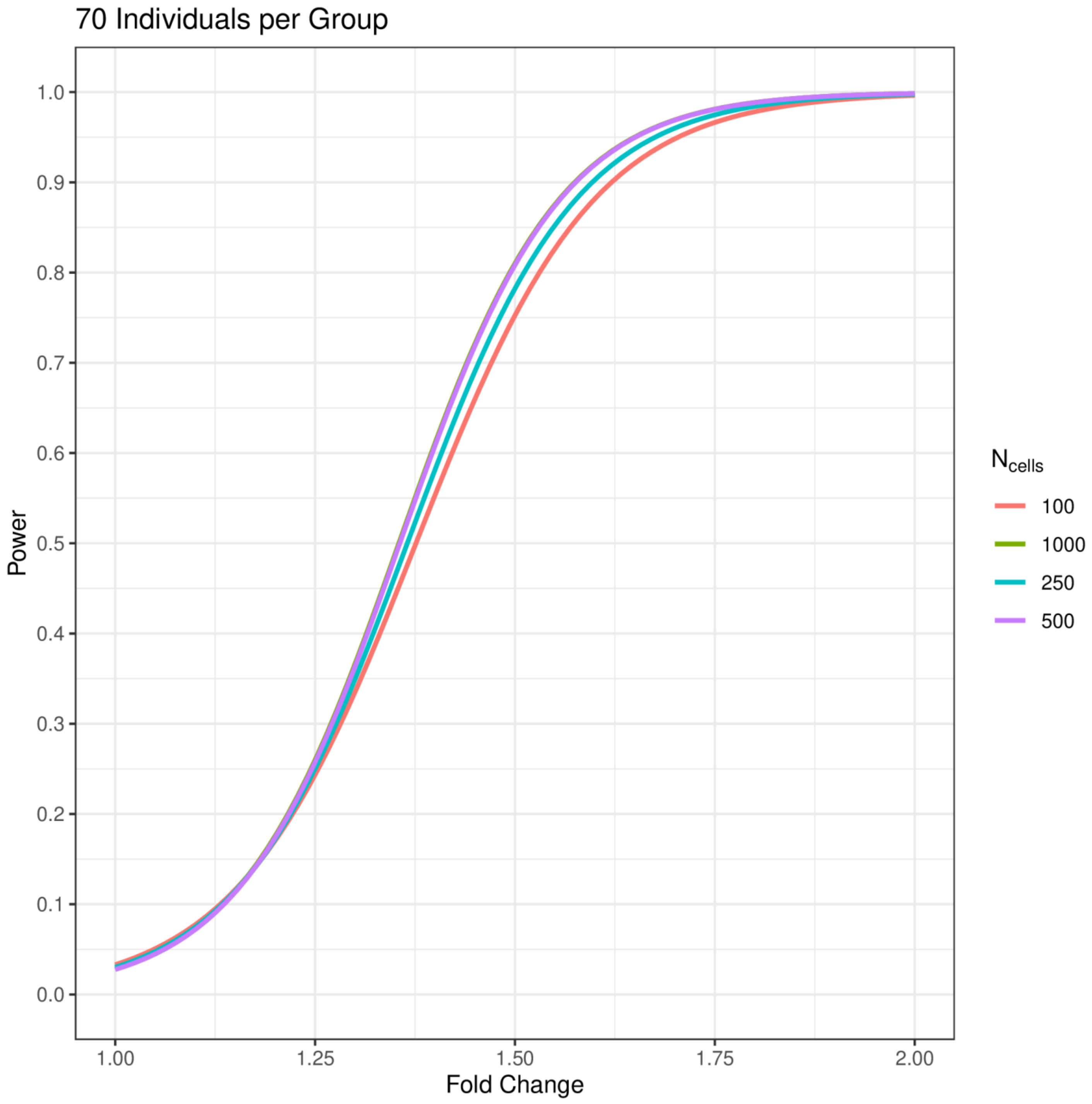
70 Individuals per Group
Power
1.0
0.9
0.8
0.7
0.6
0.5
0.4
0.3
0.2
0.1
0.0
1.00
1.25
1.50
1.75
2.00
Fold Change
$N_{cells}$
100
1000
250
500

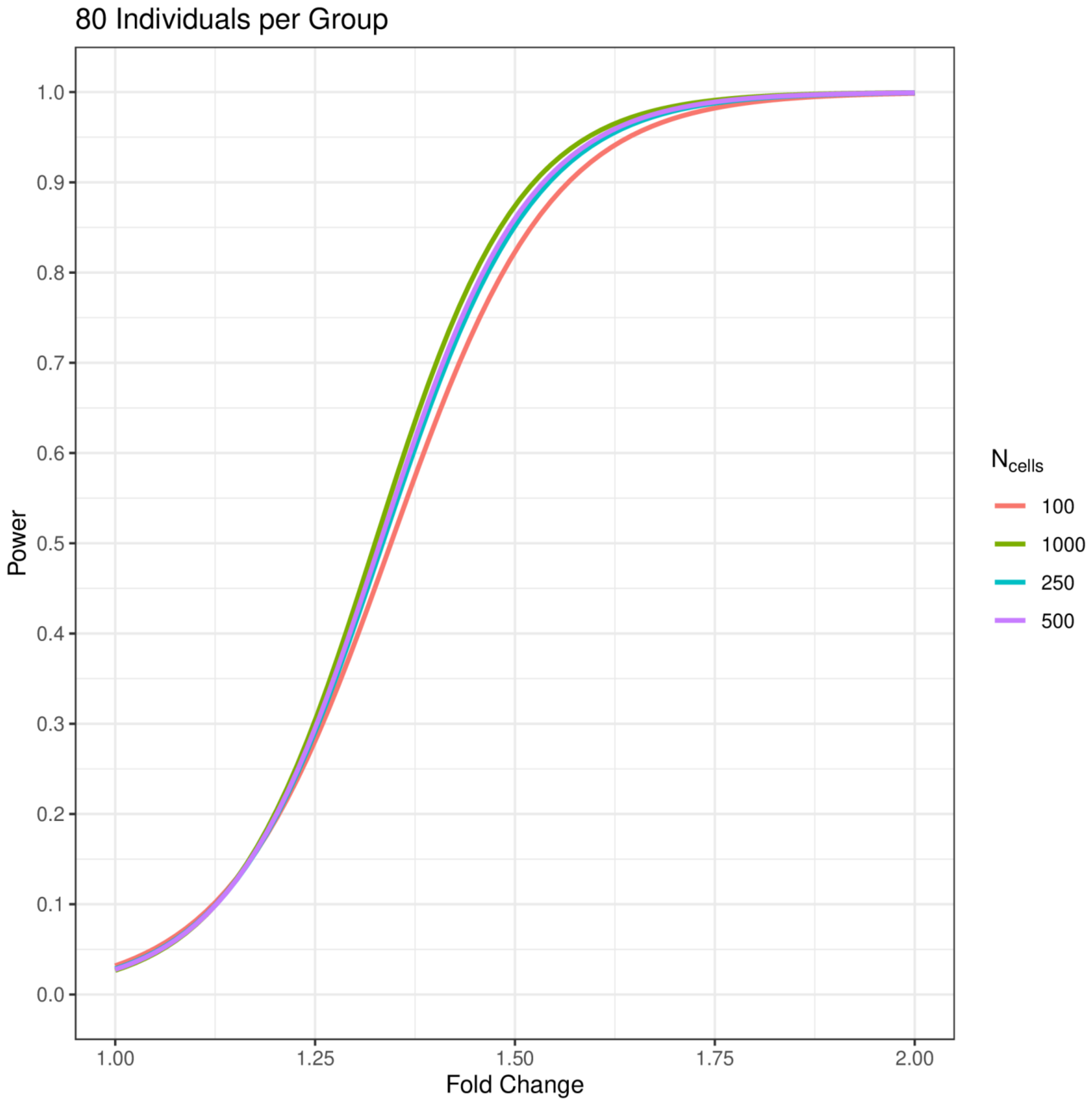
80 Individuals per Group
Power
1.0
0.9
0.8
0.7
0.6
0.5
0.4
0.3
0.2
0.1
0.0
1.00
1.25
1.50
1.75
2.00
Fold Change
$N_{cells}$
100
1000
250
500

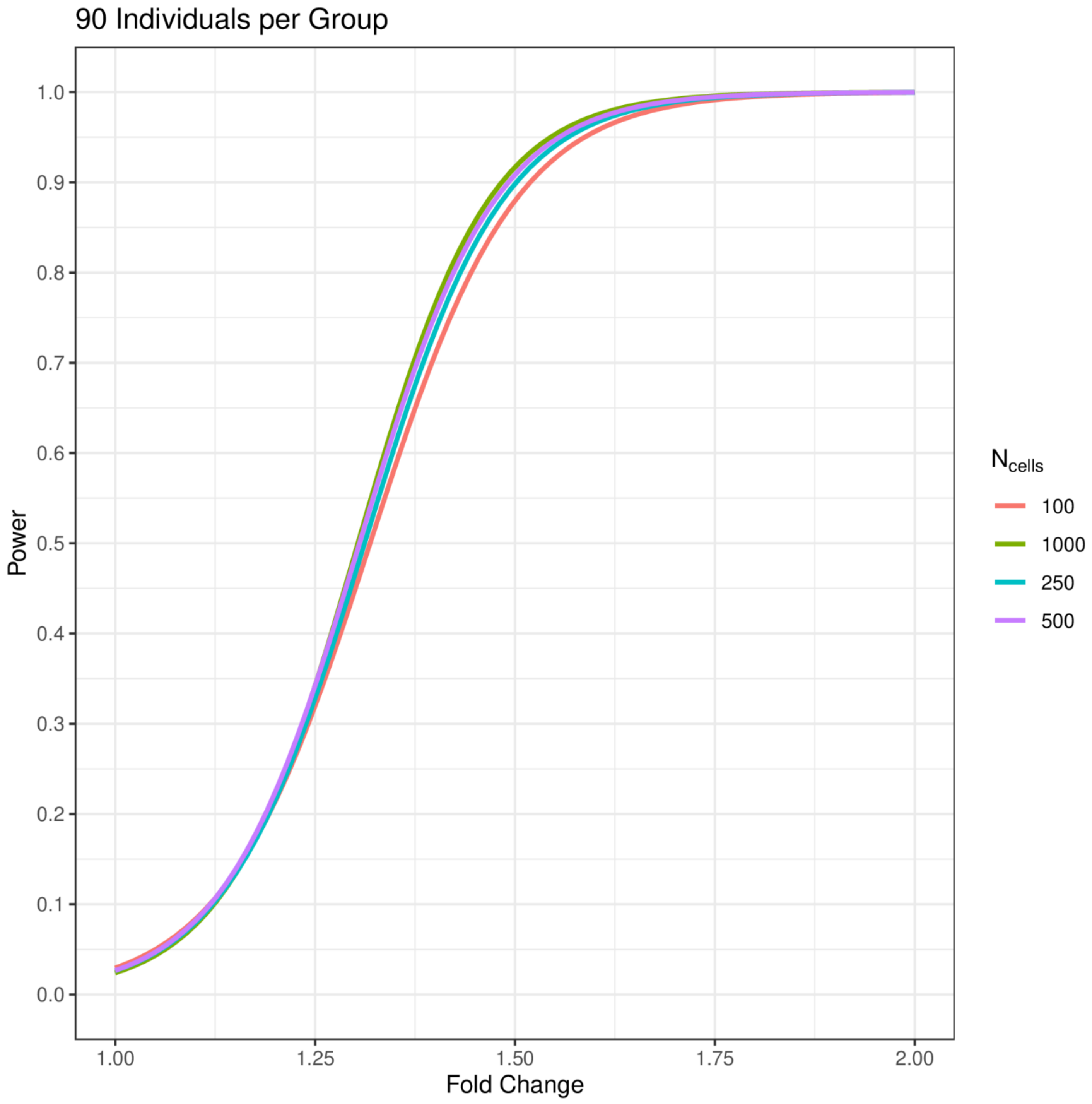
90 Individuals per Group
1.0
0.9
0.8
0.7
0.6
0.5
0.4
0.3
0.2
0.1
0.0
Power
1.00
1.25
1.50
1.75
2.00
Fold Change
$N_{cells}$
100
1000
250
500

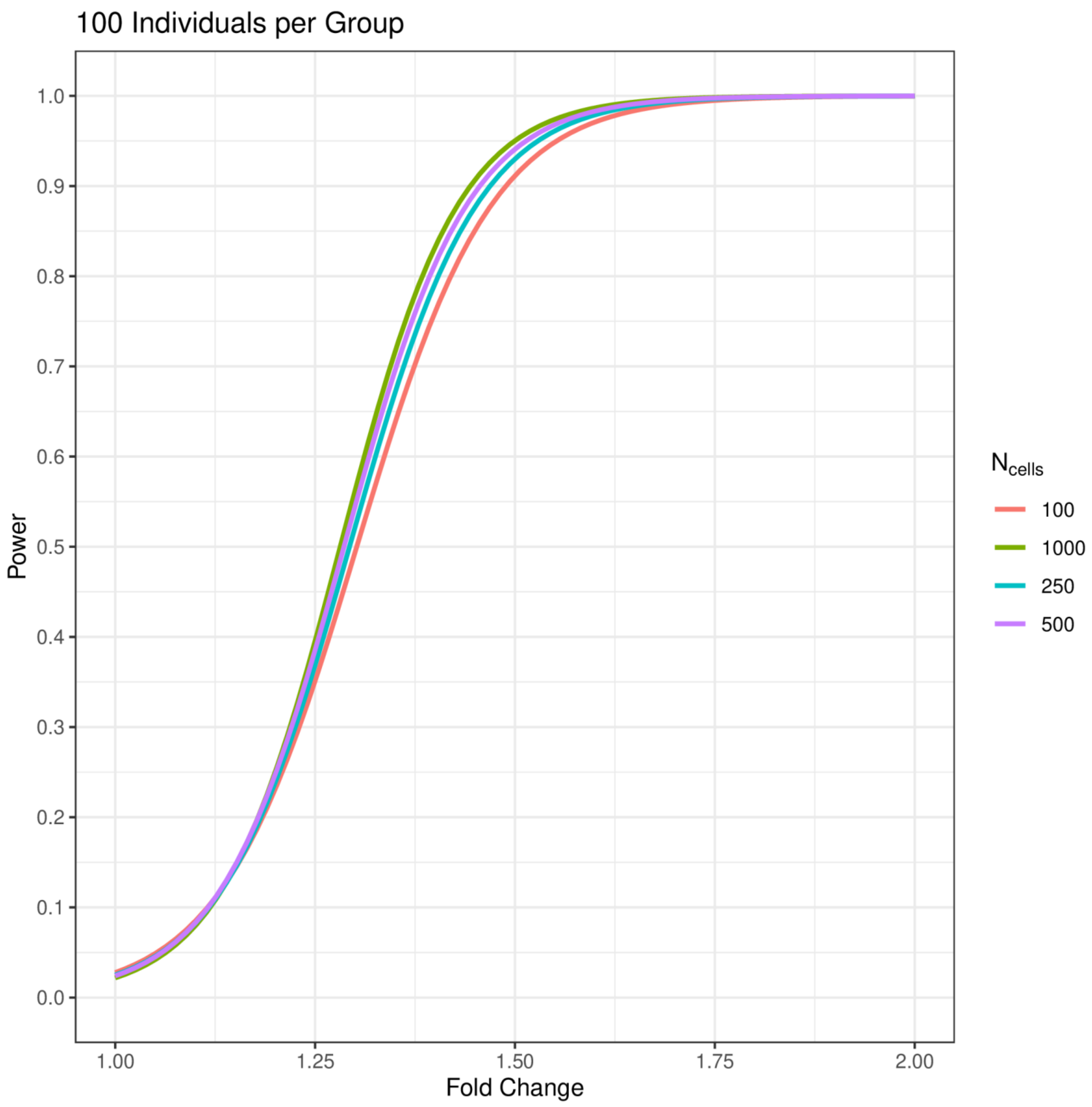
100 Individuals per Group
Power
1.0
0.9
0.8
0.7
0.6
0.5
0.4
0.3
0.2
0.1
0.0
1.00
1.25
1.50
1.75
2.00
Fold Change
N cells
100
1000
250
500

**Fig. S7. | Power calculations using MAST with a random effect for individual.** Power curves for MAST using a random effect to account for intra-individual correlation. Curves are computed for 100, 250, 500, and 1,000 cells per individual using an α = 0.001. The number of individuals per group ranges from 3 to 100 and is listed above each plot.

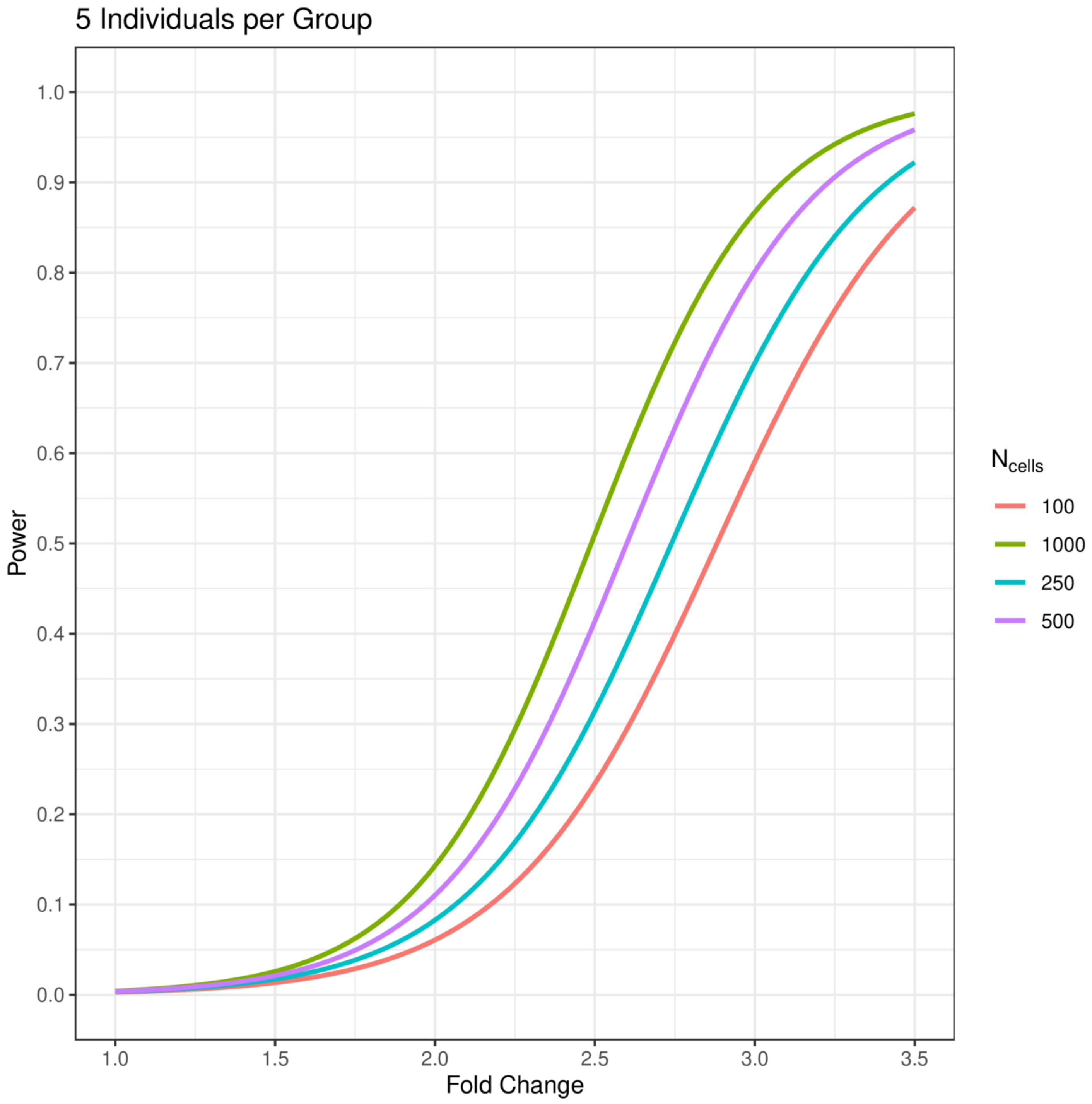

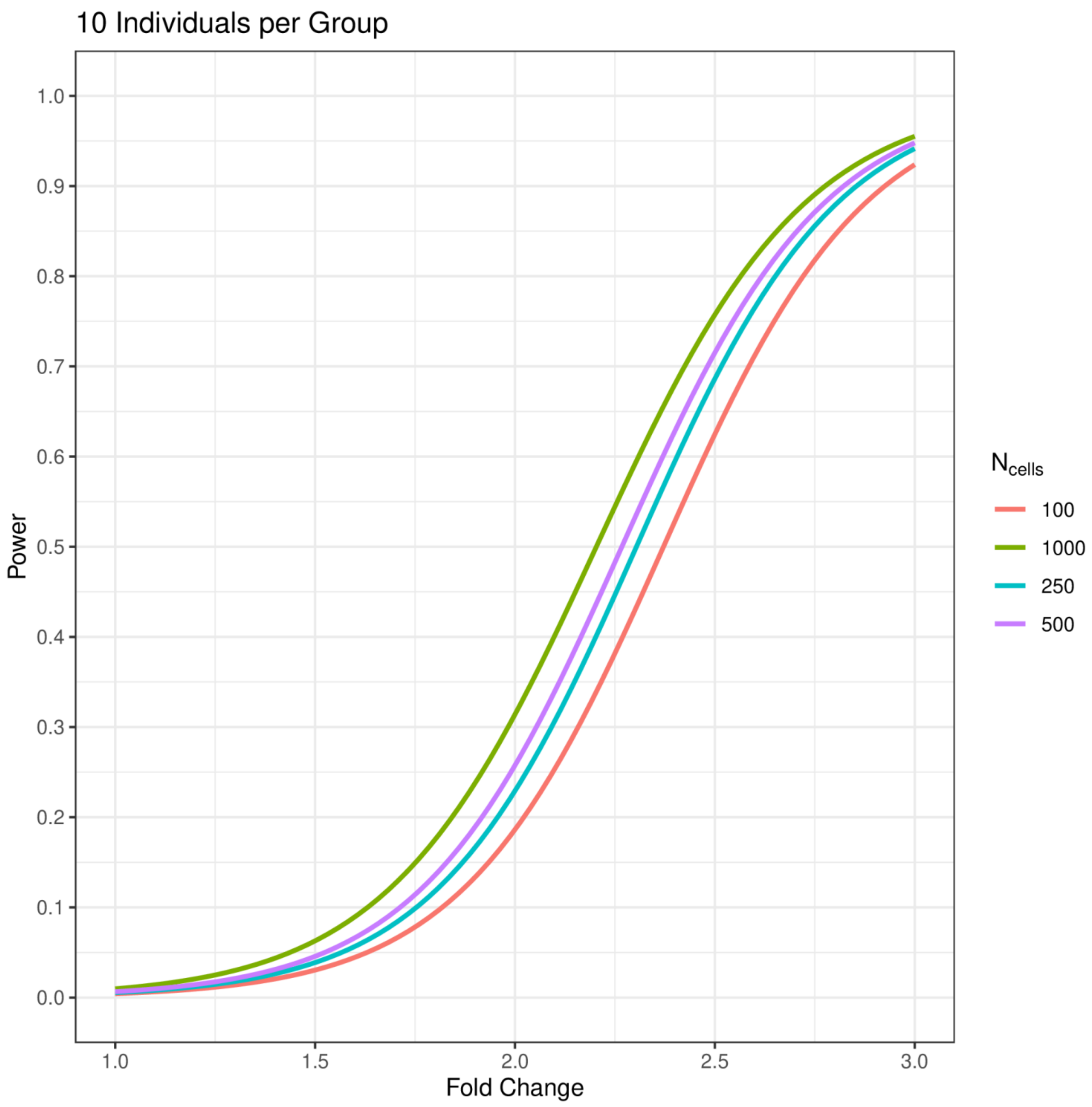
10 Individuals per Group
Power
1.0
0.9
0.8
0.7
0.6
0.5
0.4
0.3
0.2
0.1
0.0
1.0
1.5
2.0
2.5
3.0
Fold Change
$N_{cells}$
100
1000
250
500

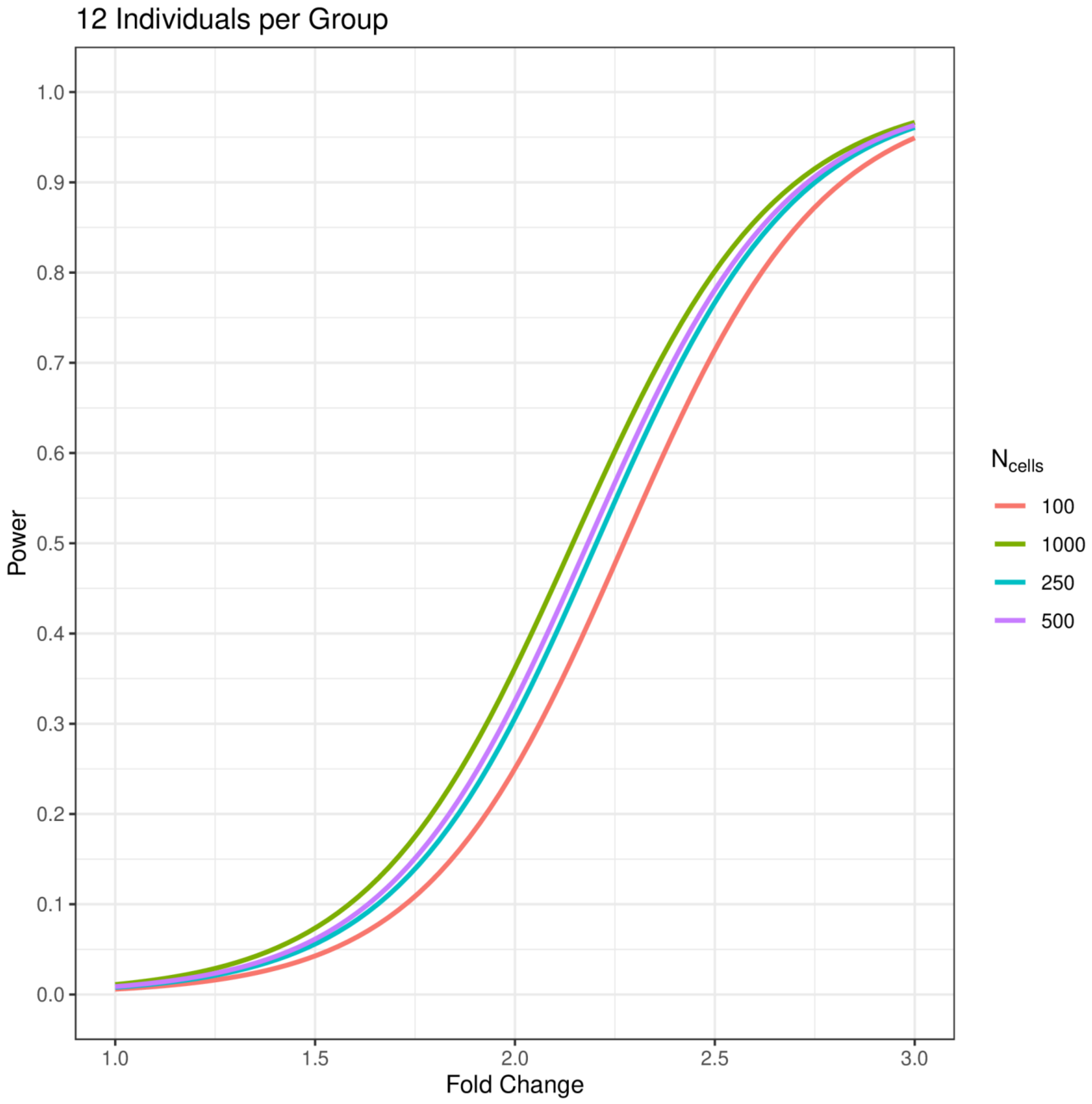

12 Individuals per Group
Power
1.0
0.9
0.8
0.7
0.6
0.5
0.4
0.3
0.2
0.1
0.0
1.0
1.5
2.0
2.5
3.0
Fold Change
$N_{cells}$
100
1000
250
500

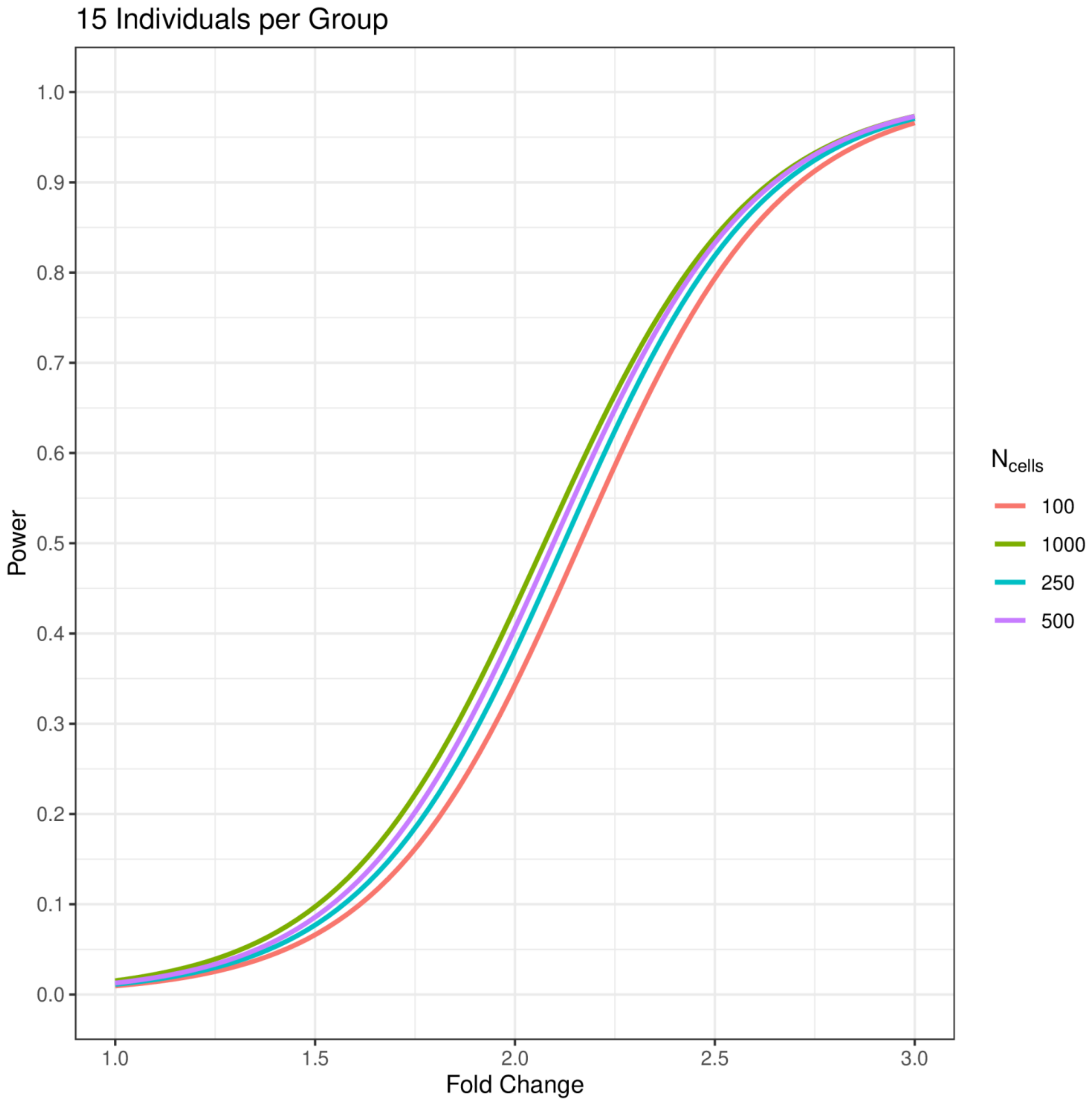
15 Individuals per Group
1.0
0.9
0.8
0.7
0.6
0.5
0.4
0.3
0.2
0.1
0.0
Power
1.0
1.5
2.0
2.5
3.0
Fold Change
N cells
100
1000
250
500

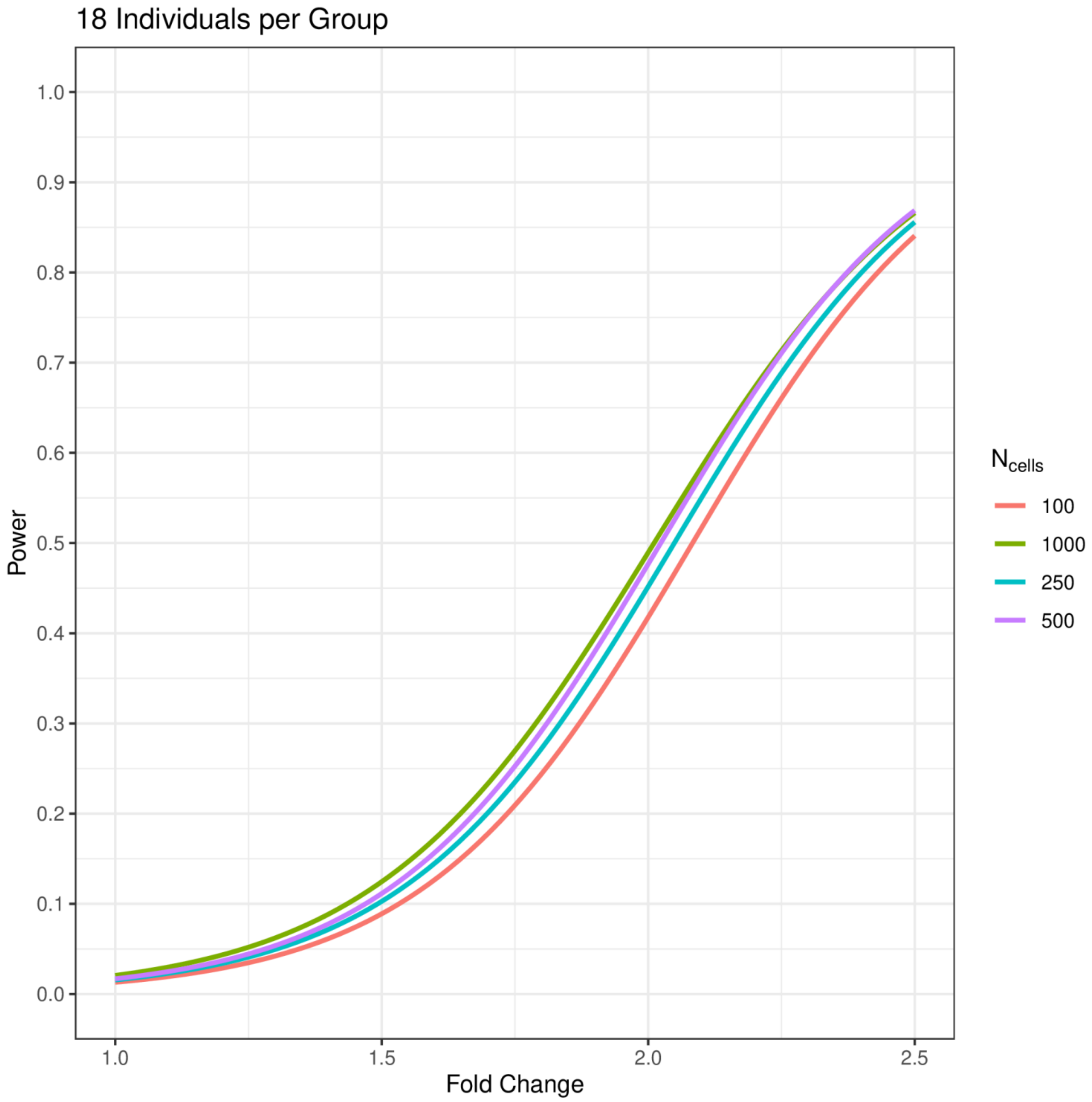
18 Individuals per Group
Power
1.0
0.9
0.8
0.7
0.6
0.5
0.4
0.3
0.2
0.1
0.0
1.0
1.5
2.0
2.5
Fold Change
N cells
100
1000
250
500

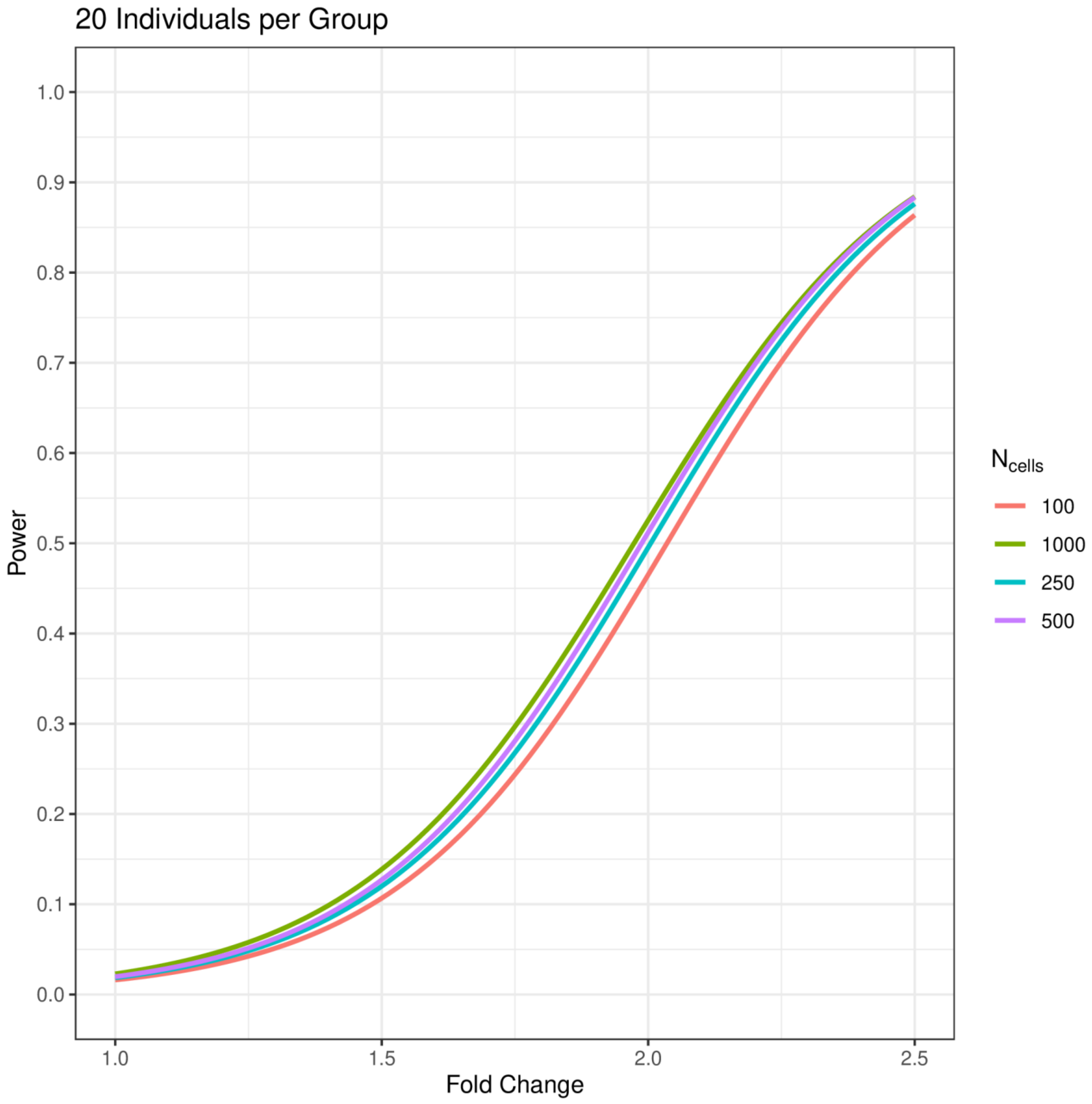

20 Individuals per Group
Power
1.0
0.9
0.8
0.7
0.6
0.5
0.4
0.3
0.2
0.1
0.0
1.0
1.5
2.0
2.5
Fold Change
$N_{cells}$
100
1000
250
500

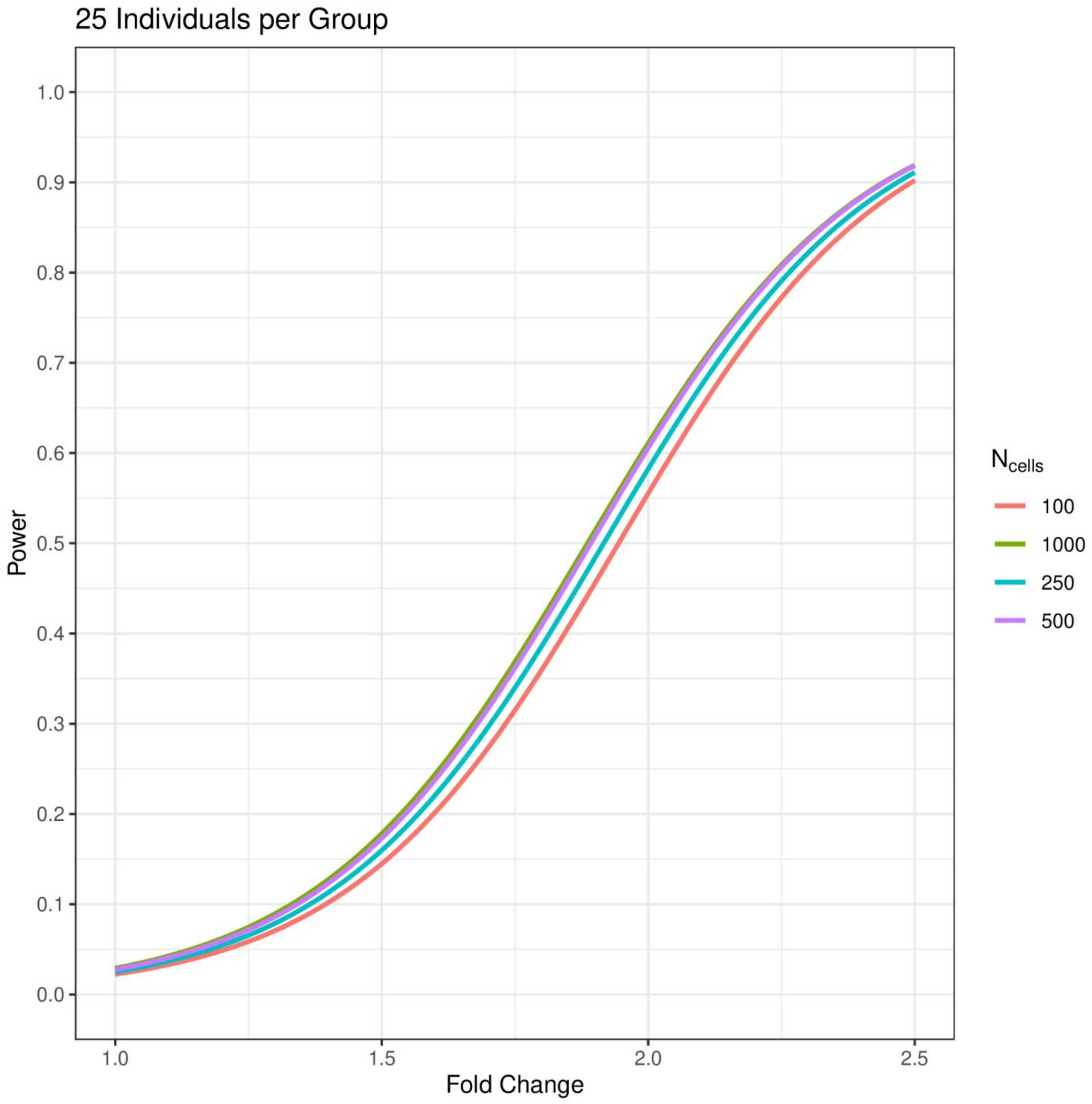
25 Individuals per Group
Power
Fold Change
1.0
0.9
0.8
0.7
0.6
0.5
0.4
0.3
0.2
0.1
0.0
1.0
1.5
2.0
2.5
N cells
100
1000
250
500

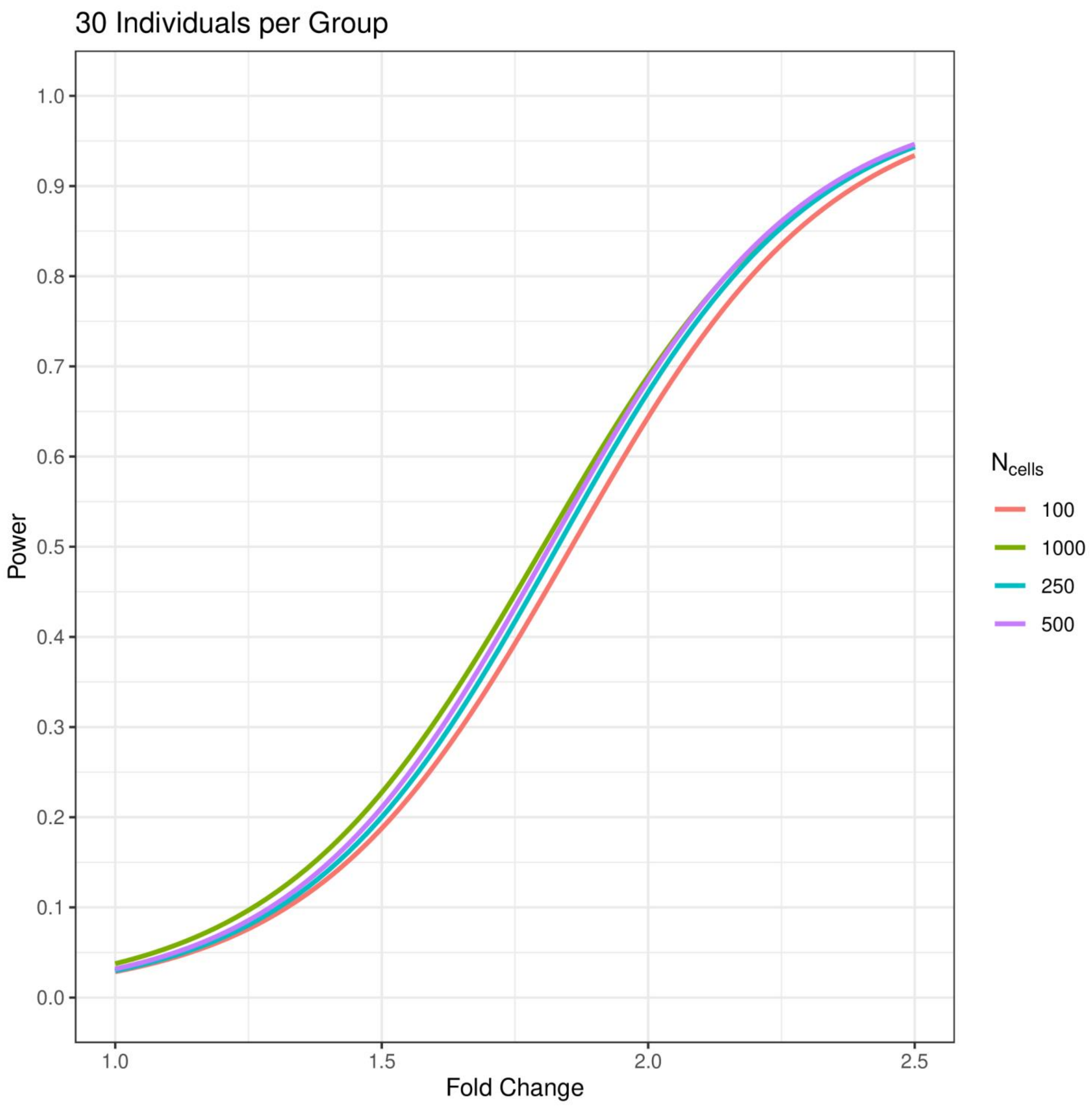
30 Individuals per Group
Power
Fold Change
1.0
0.9
0.8
0.7
0.6
0.5
0.4
0.3
0.2
0.1
0.0
1.0
1.5
2.0
2.5
$N_{cells}$
100
1000
250
500

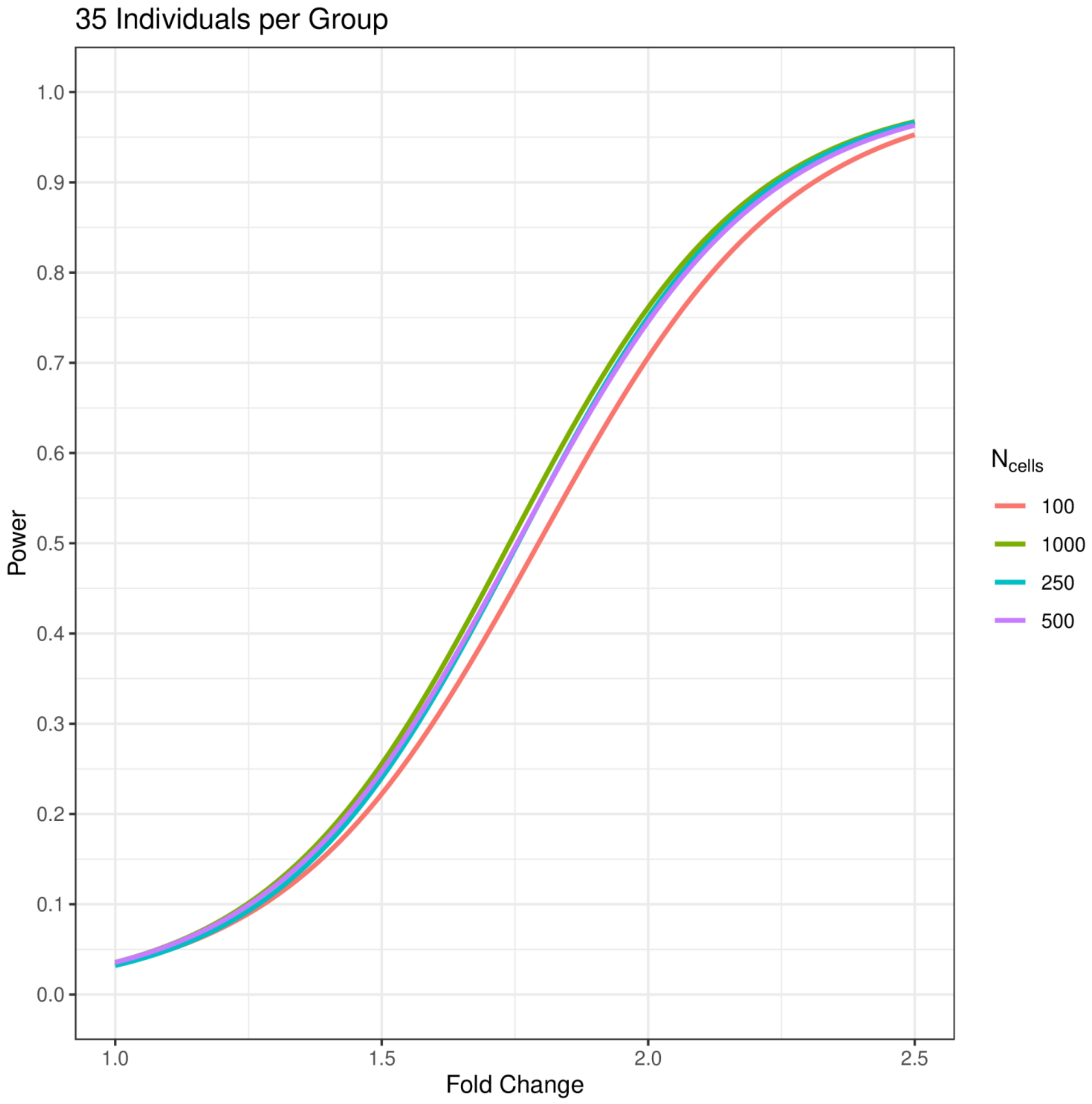

35 Individuals per Group
1.0
0.9
0.8
0.7
0.6
0.5
0.4
0.3
0.2
0.1
0.0
Power
1.0
1.5
2.0
2.5
Fold Change
$N_{cells}$
100
1000
250
500

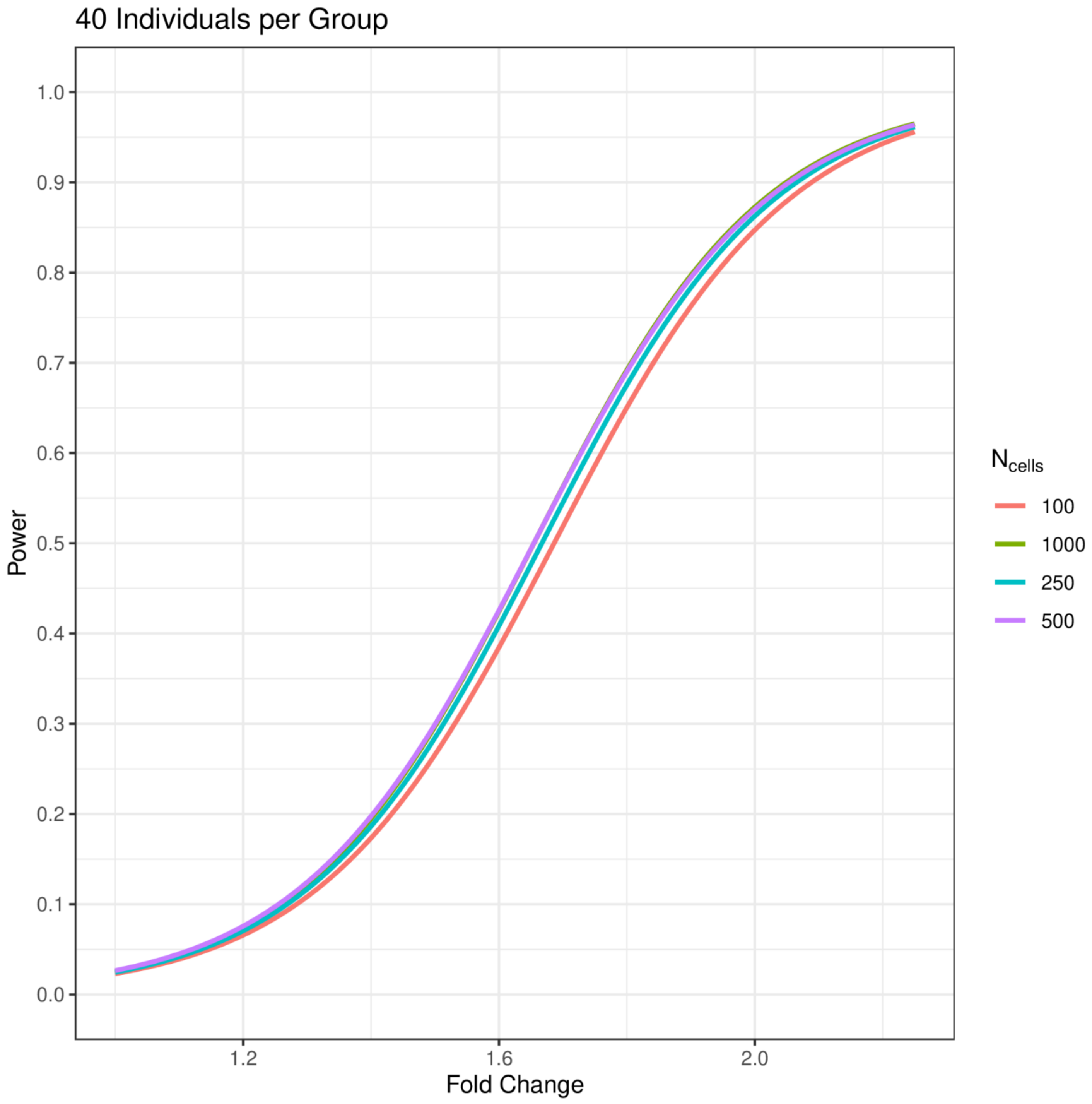
40 Individuals per Group
Power
Fold Change
1.0
0.9
0.8
0.7
0.6
0.5
0.4
0.3
0.2
0.1
0.0
1.2
1.6
2.0
$N_{cells}$
100
1000
250
500

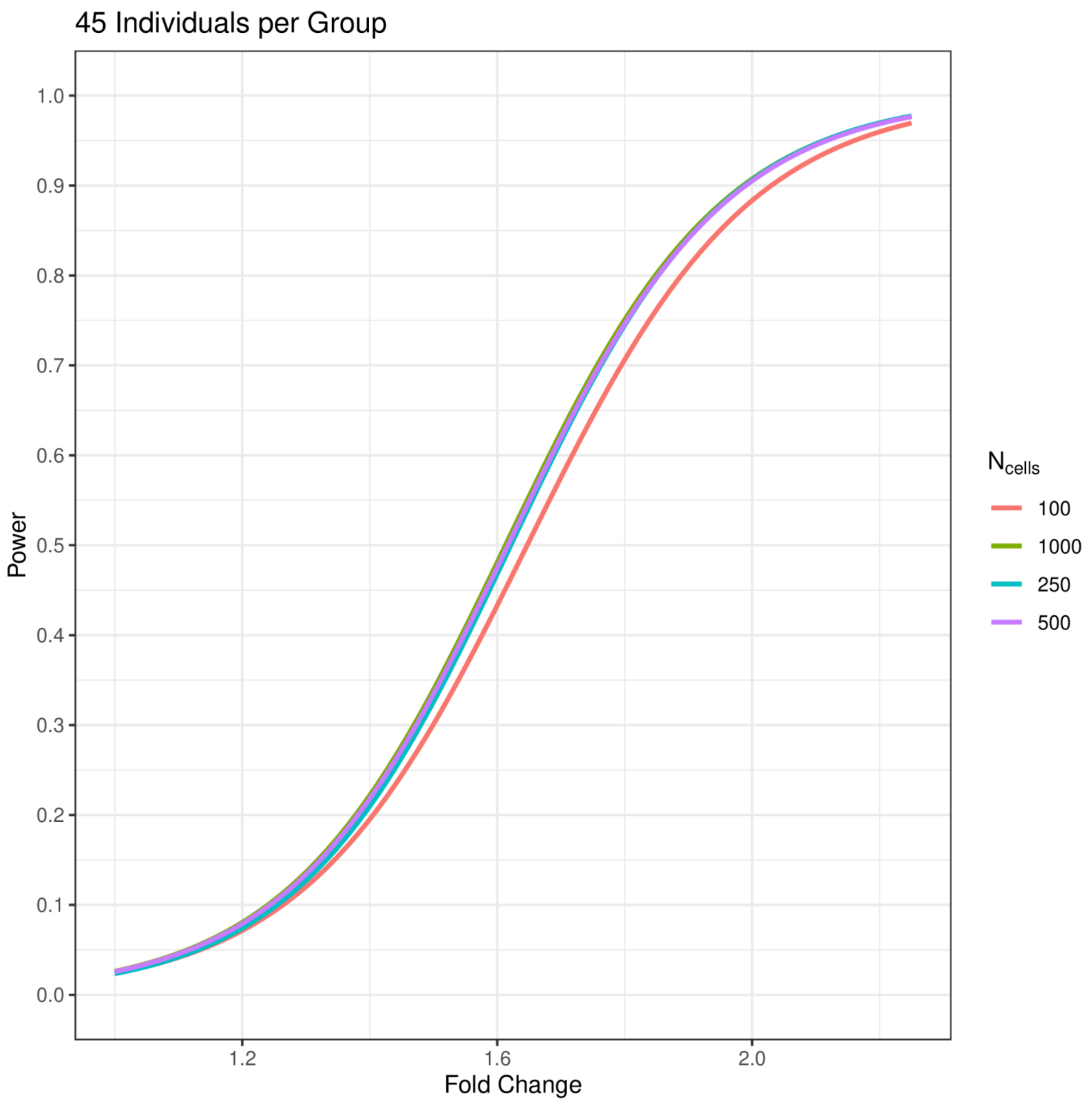

45 Individuals per Group
Power
Fold Change
1.0
0.9
0.8
0.7
0.6
0.5
0.4
0.3
0.2
0.1
0.0
1.2
1.6
2.0
N cells
100
1000
250
500

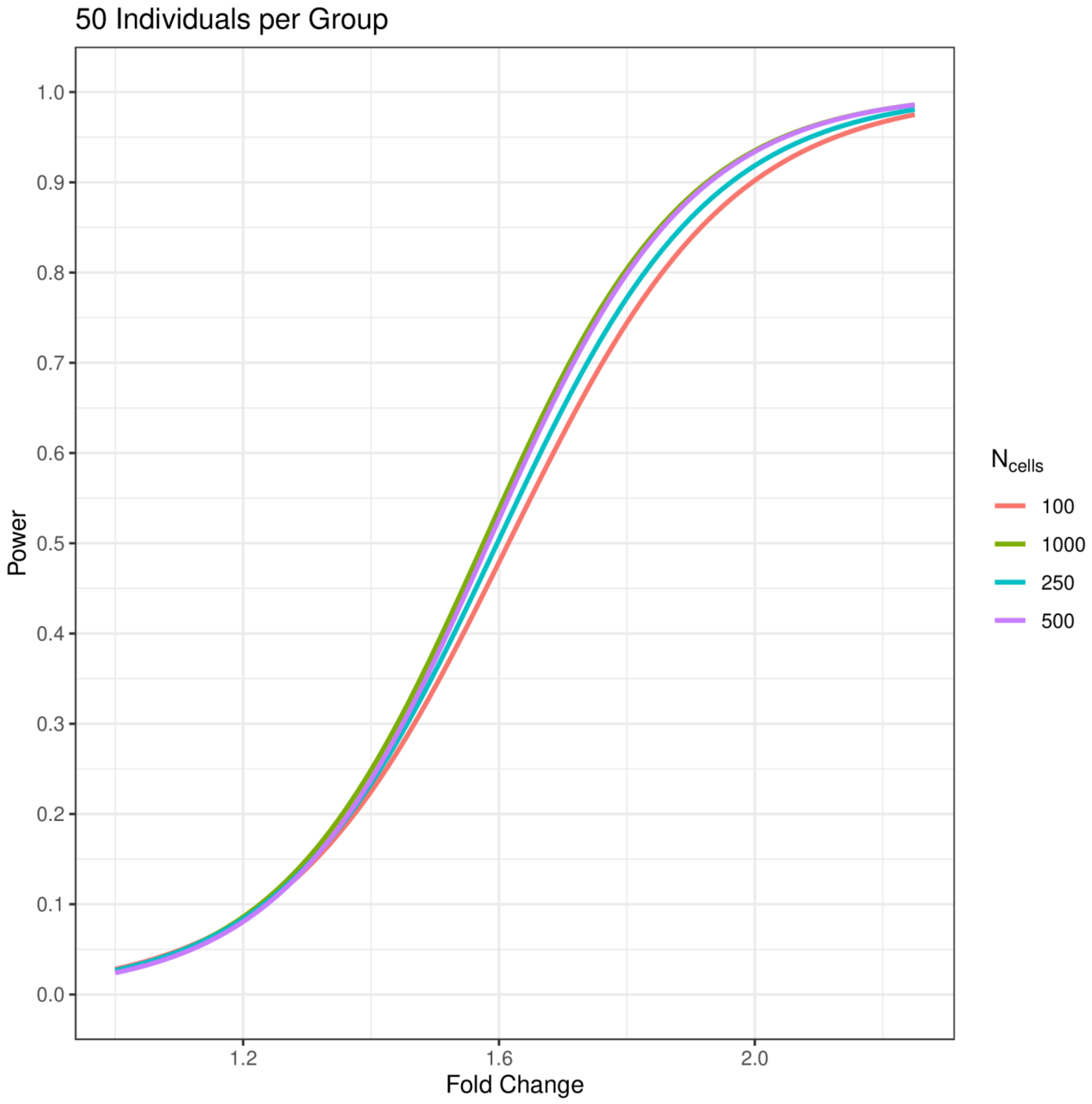
50 Individuals per Group
1.0
0.9
0.8
0.7
0.6
0.5
0.4
0.3
0.2
0.1
0.0
Power
1.2
1.6
2.0
Fold Change
$N_{cells}$
100
1000
250
500

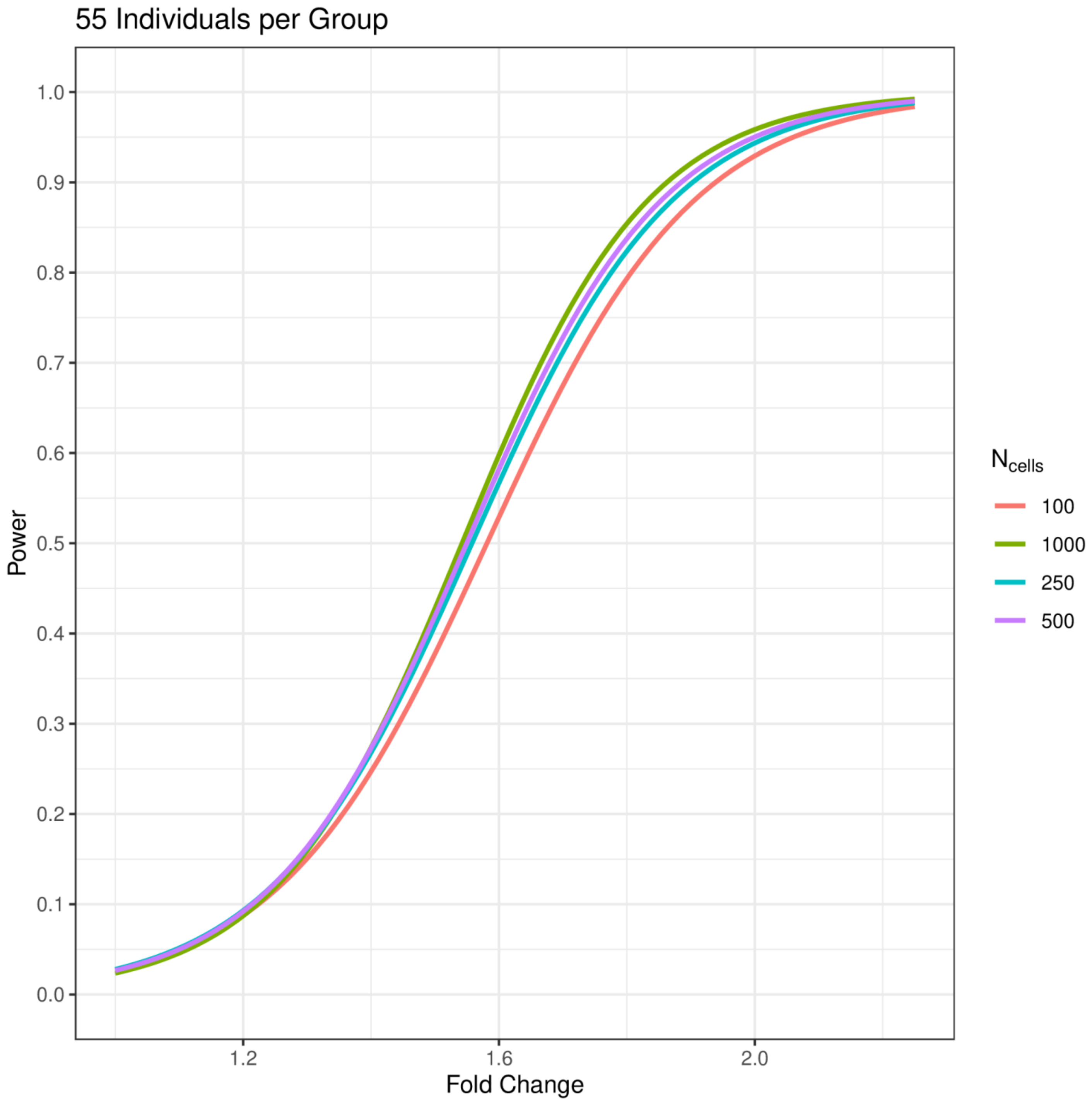
55 Individuals per Group
Power
1.0
0.9
0.8
0.7
0.6
0.5
0.4
0.3
0.2
0.1
0.0
1.2
1.6
2.0
Fold Change
N cells
100
1000
250
500

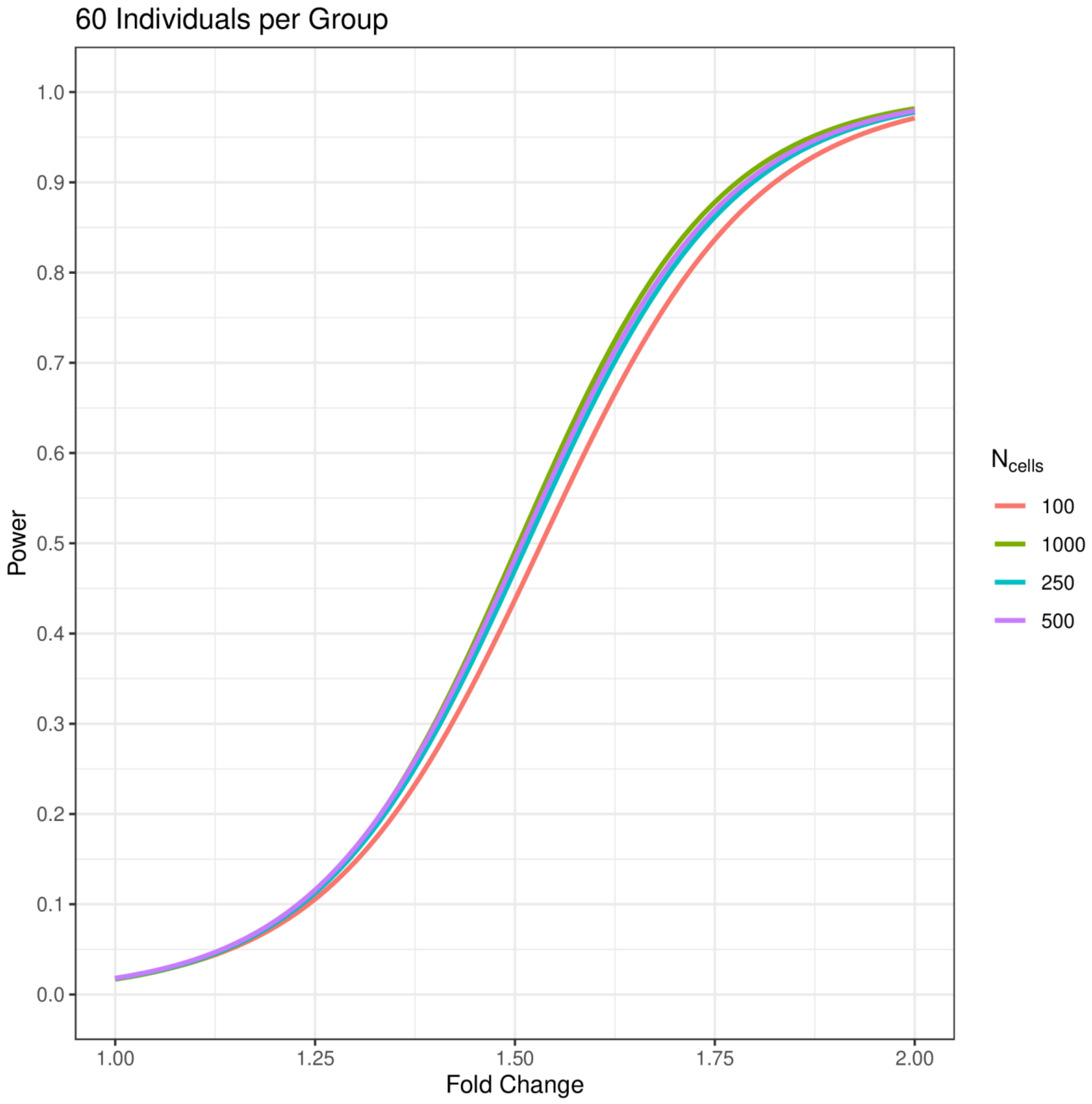
60 Individuals per Group
Power
1.0
0.9
0.8
0.7
0.6
0.5
0.4
0.3
0.2
0.1
0.0
1.00
1.25
1.50
1.75
2.00
Fold Change
$N_{cells}$
100
1000
250
500

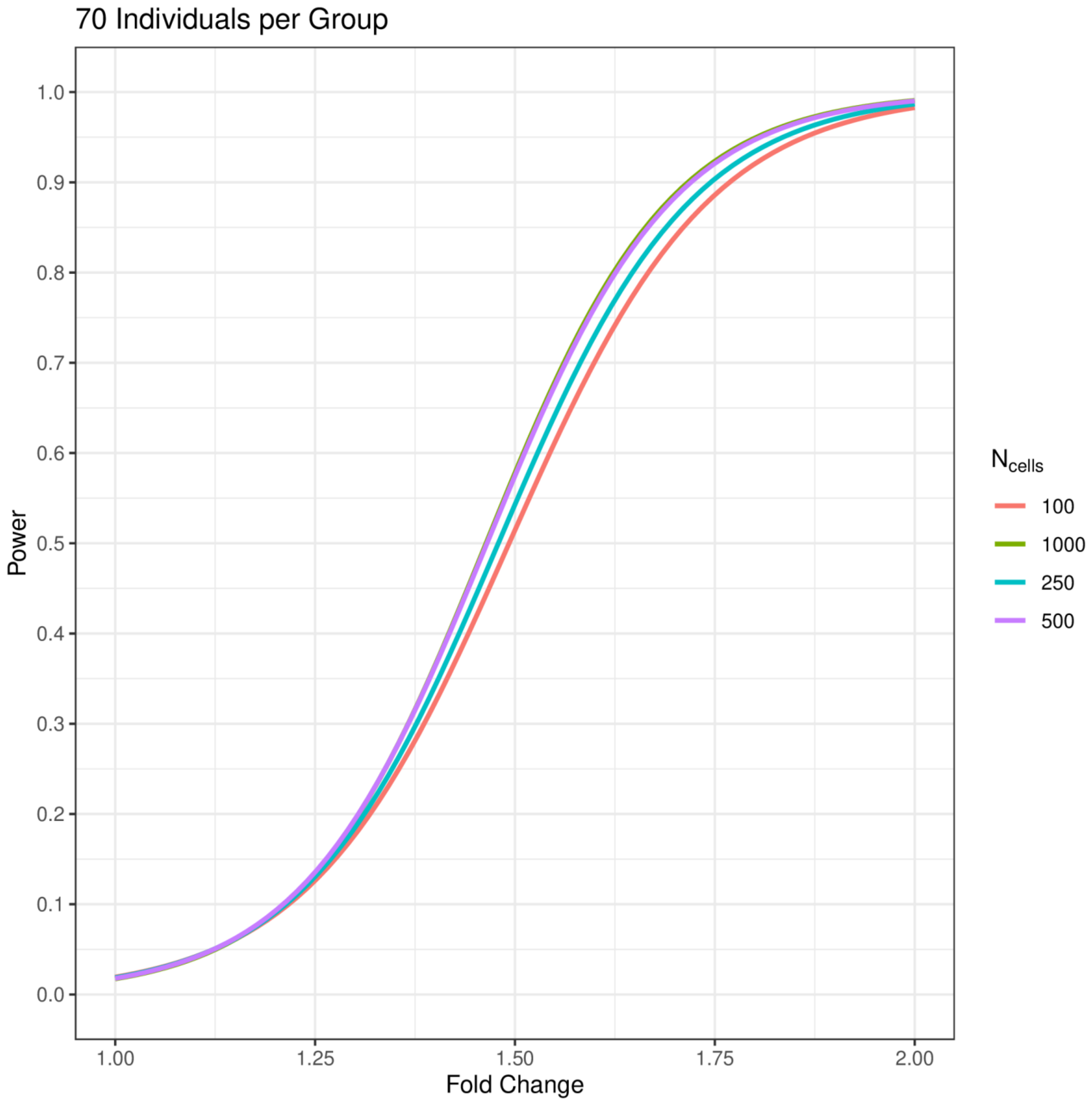
70 Individuals per Group
Power
Fold Change
1.0
0.9
0.8
0.7
0.6
0.5
0.4
0.3
0.2
0.1
0.0
1.00
1.25
1.50
1.75
2.00
$N_{cells}$
100
1000
250
500

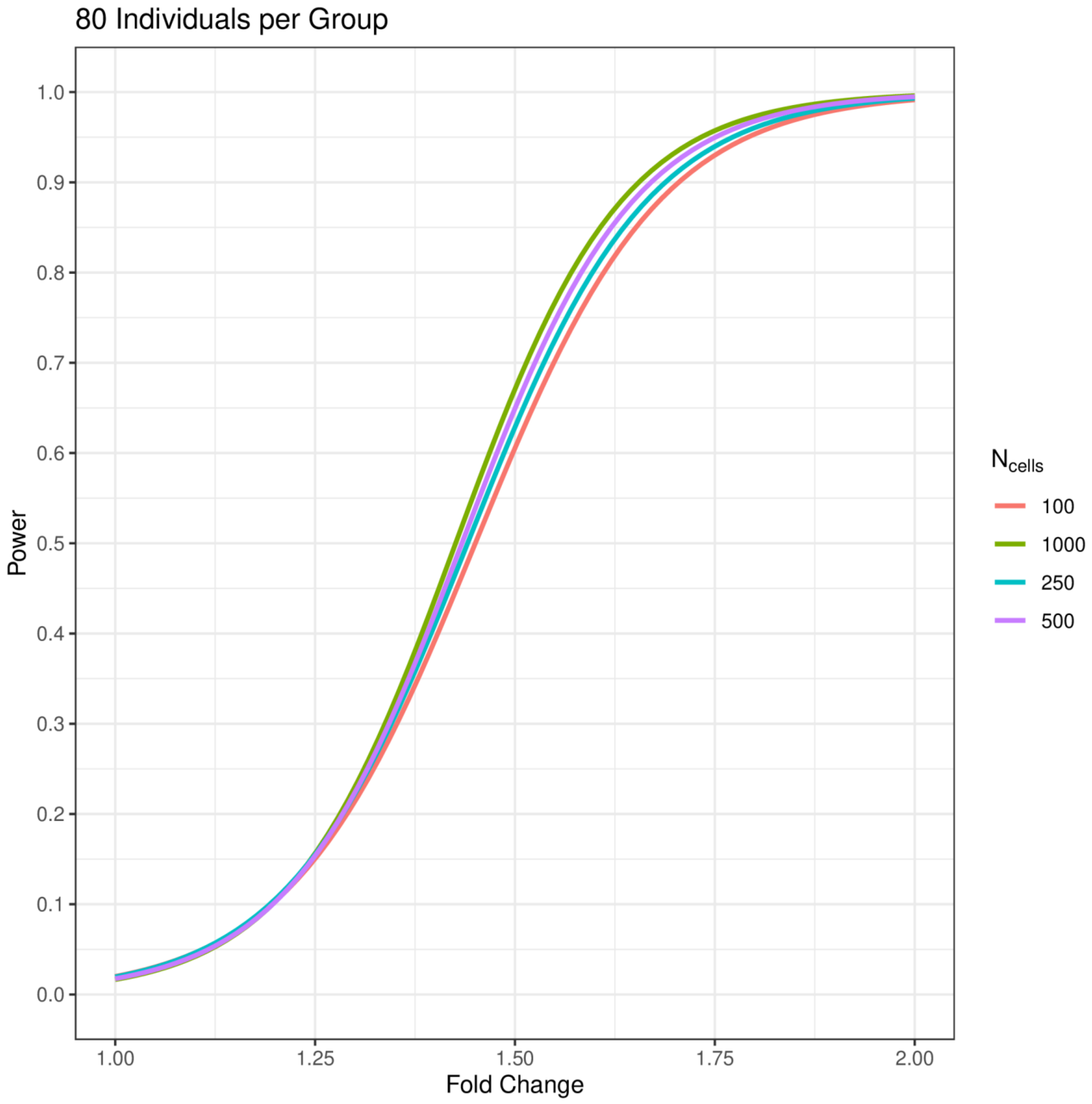
80 Individuals per Group
Power
1.0
0.9
0.8
0.7
0.6
0.5
0.4
0.3
0.2
0.1
0.0
1.00
1.25
1.50
1.75
2.00
Fold Change
$N_{cells}$
100
1000
250
500

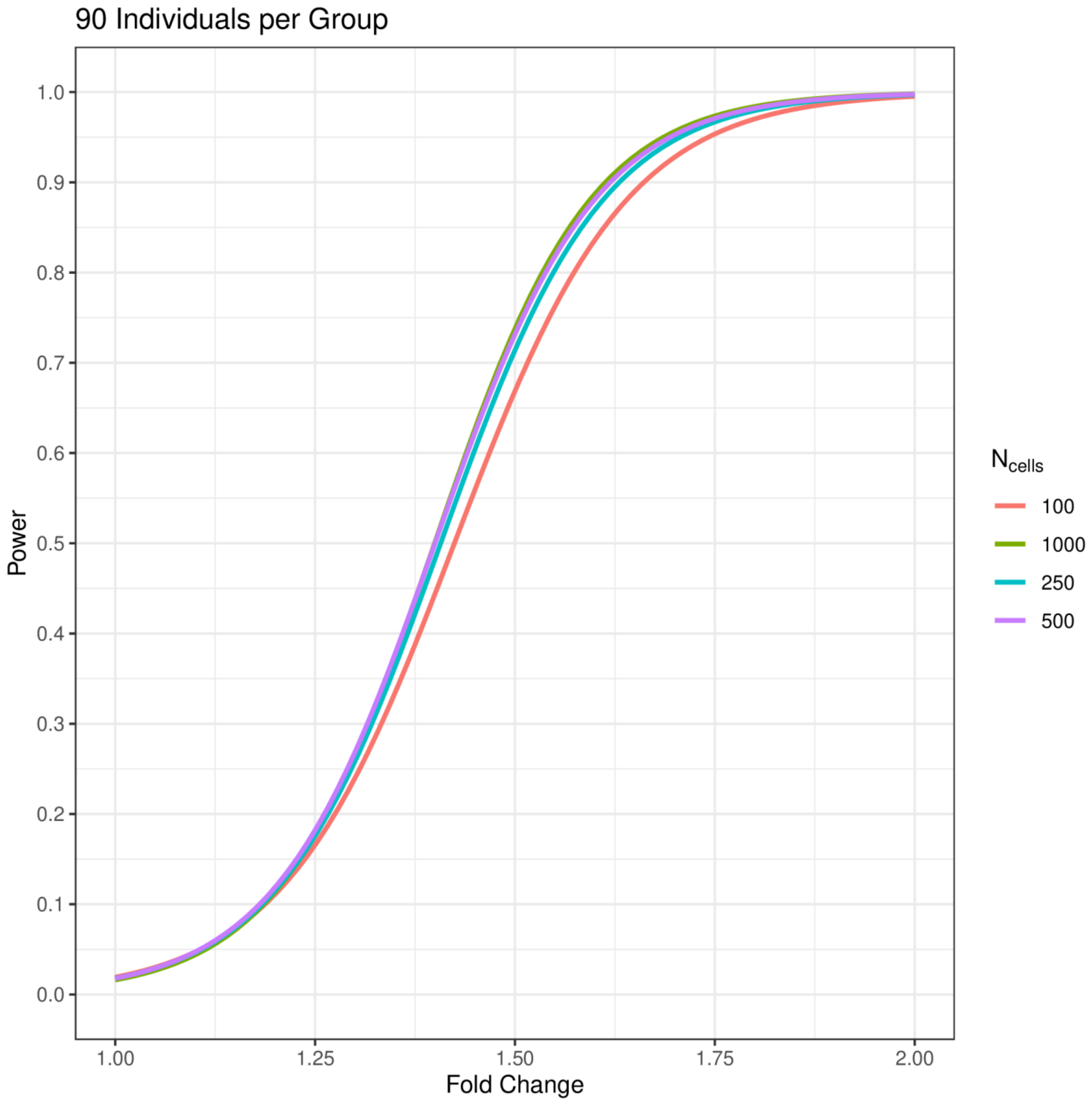

90 Individuals per Group
Power
1.0
0.9
0.8
0.7
0.6
0.5
0.4
0.3
0.2
0.1
0.0
1.00
1.25
1.50
1.75
2.00
Fold Change
$N_{cells}$
100
1000
250
500

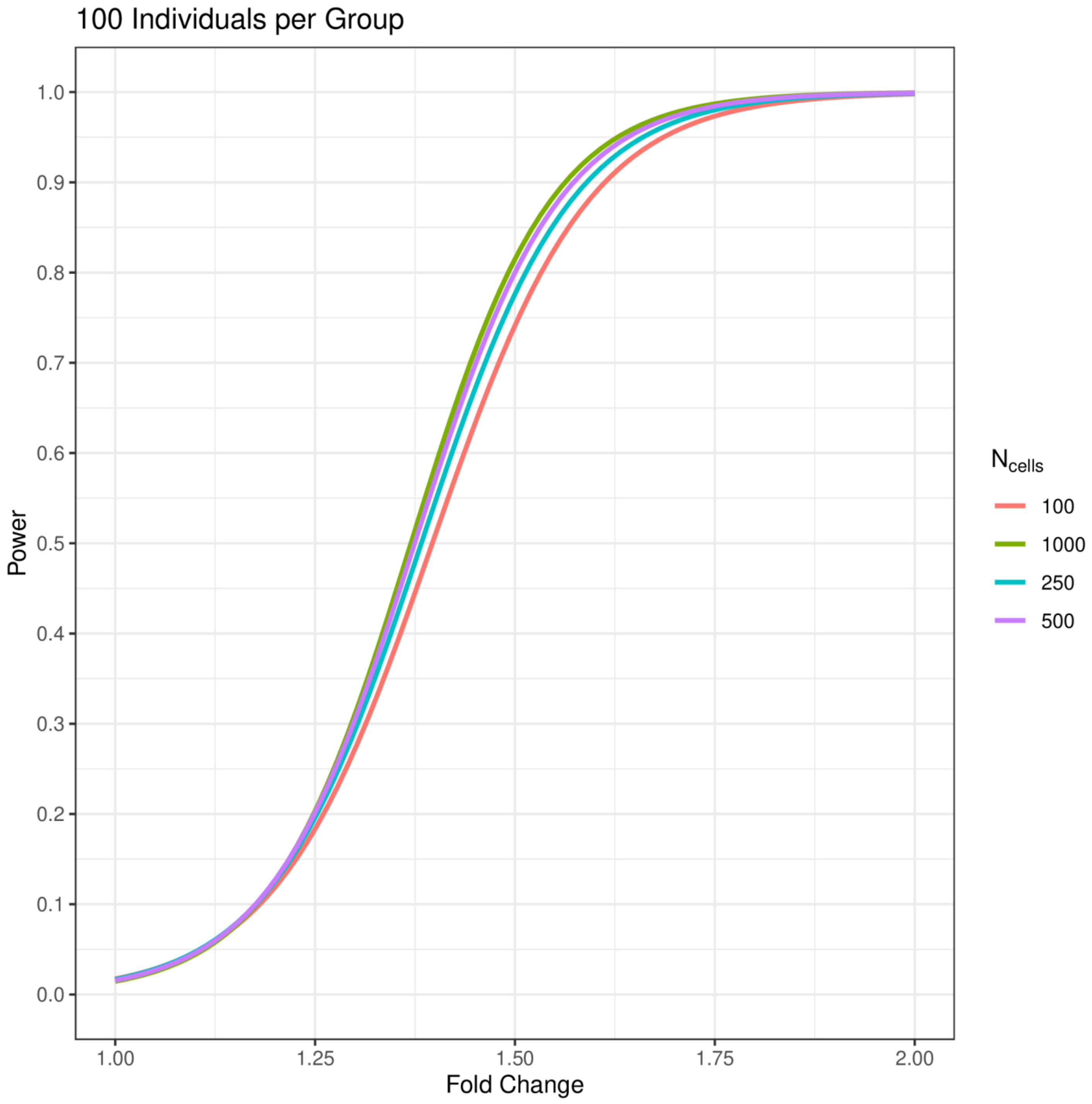
100 Individuals per Group
Power
1.0
0.9
0.8
0.7
0.6
0.5
0.4
0.3
0.2
0.1
0.0
1.00
1.25
1.50
1.75
2.00
Fold Change
$N_{cells}$
100
1000
250
500

**Table S1.** Type I error rates of some of the currently applied tools in single-cell analysis. Type I error rates of ten different methods under twenty different conditions and a significance threshold of $p<0.05$. 250,000 iterations were computed to obtain an error rate for each method. The inflated type I error rates computed with mixed models at the lower numbers of individuals per group are a consequence of the two-part hurdle model simultaneously testing two hypotheses and an overabundance of sub-sampling with small sample sizes. Type I error rates are well controlled for with mixed models and pseudobulk methods, while type I error rates inflate with other methods as additional independent samples or more cells are added. Pseudobulk methods are overly conservative.

| | | Two-part Hurdle | | | Tweedie | | | Pseudobulk | | | |
|---|---|---|---|---|---|---|---|---|---|---|---|
| $N_{ind}$ | $N_{cells}$ | Default | Corrected | RE | GLMM | GLM | GEE1 | Mean | Sum | Tobit | Modified *t* |
| 5 | 50 | 0.561 | 0.637 | 0.069 | 0.082 | 0.340 | 0.114 | 0.023 | 0.035 | 0.353 | 0.400 |
| | 100 | 0.677 | 0.719 | 0.064 | 0.084 | 0.463 | 0.110 | 0.022 | 0.032 | 0.471 | 0.510 |
| | 250 | 0.798 | 0.778 | 0.066 | 0.083 | 0.609 | 0.103 | 0.023 | 0.028 | 0.628 | 0.644 |
| | 500 | 0.862 | 0.803 | 0.065 | 0.081 | 0.705 | 0.104 | 0.023 | 0.026 | 0.725 | 0.718 |
| 10 | 50 | 0.563 | 0.611 | 0.055 | 0.064 | 0.350 | 0.076 | 0.024 | 0.021 | 0.345 | 0.397 |
| | 100 | 0.689 | 0.718 | 0.053 | 0.065 | 0.462 | 0.077 | 0.024 | 0.020 | 0.470 | 0.502 |
| | 250 | 0.810 | 0.793 | 0.049 | 0.064 | 0.610 | 0.074 | 0.022 | 0.019 | 0.624 | 0.635 |
| | 500 | 0.875 | 0.827 | 0.049 | 0.061 | 0.705 | 0.073 | 0.021 | 0.018 | 0.722 | 0.717 |
| 20 | 50 | 0.562 | 0.606 | 0.051 | 0.056 | 0.344 | 0.063 | 0.024 | 0.016 | 0.343 | 0.393 |
| | 100 | 0.687 | 0.705 | 0.048 | 0.056 | 0.459 | 0.064 | 0.024 | 0.014 | 0.466 | 0.503 |
| | 250 | 0.817 | 0.805 | 0.042 | 0.058 | 0.610 | 0.060 | 0.022 | 0.011 | 0.619 | 0.637 |
| | 500 | 0.884 | 0.844 | 0.042 | 0.055 | 0.705 | 0.062 | 0.021 | 0.010 | 0.720 | 0.716 |
| 30 | 50 | 0.563 | 0.604 | 0.053 | 0.054 | 0.341 | 0.058 | 0.025 | 0.013 | 0.344 | 0.395 |
| | 100 | 0.691 | 0.698 | 0.049 | 0.056 | 0.463 | 0.058 | 0.025 | 0.012 | 0.469 | 0.504 |
| | 250 | 0.818 | 0.803 | 0.044 | 0.055 | 0.608 | 0.057 | 0.022 | 0.010 | 0.624 | 0.636 |
| | 500 | 0.886 | 0.853 | 0.041 | 0.055 | 0.707 | 0.058 | 0.022 | 0.009 | 0.719 | 0.706 |
| 40 | 50 | 0.561 | 0.602 | 0.051 | 0.054 | 0.345 | 0.055 | 0.025 | 0.013 | 0.340 | 0.393 |
| | 100 | 0.689 | 0.699 | 0.049 | 0.053 | 0.455 | 0.055 | 0.026 | 0.012 | 0.467 | 0.502 |
| | 250 | 0.820 | 0.803 | 0.044 | 0.053 | 0.607 | 0.053 | 0.022 | 0.010 | 0.622 | 0.639 |
| | 500 | 0.888 | 0.856 | 0.042 | 0.053 | 0.704 | 0.054 | 0.022 | 0.008 | 0.721 | 0.713 |

*Default denotes MAST was implemented without random-effects, RE denotes random-effects, Corrected denotes data was batch-corrected for individual prior to analysis without using individual as a random-effect, GLM denotes generalized linear model, and GLMM denotes generalized linear mixed-effects model.

**Two-part Hurdle model as implemented in MAST, Tweedie distribution as implemented in 'glmmTMB', GEE1 as implemented in 'geepack', Pseudobulk averaged or summed across cells within an individual and was implemented in DESeq2, Modified t as implemented in ROTS, and Tobit as implemented in Monocle.

**Table S2.** Type I error rates of some of the currently applied tools in single-cell analysis. Type I error rates of ten different methods under twenty different conditions and a significance threshold of $p<0.01$. 250,000 iterations were computed to obtain an error rate for each method. The inflated type I error rates computed with mixed models at the lower numbers of individuals per group are a consequence of the two-part hurdle model simultaneously testing two hypotheses and an overabundance of sub-sampling with small sample sizes. Type I error rates are well controlled for with mixed models and pseudobulk methods, while type I error rates inflate with other methods as additional independent samples or more cells are added. Pseudobulk methods are overly conservative.

| | | **Two-part Hurdle** | | | **Tweedie** | | | **Pseudobulk** | | | |
|---|---|---|---|---|---|---|---|---|---|---|---|
| **$N_{ind}$** | **$N_{cells}$** | **Default** | **Corrected** | **RE** | **GLMM** | **GLM** | **GEE1** | **Mean** | **Sum** | **Tobit** | **Modified *t*** |
| 5 | 50 | 0.447 | 0.567 | 0.016 | 0.028 | 0.232 | 0.046 | 0.005 | 0.011 | 0.233 | 0.286 |
| | 100 | 0.584 | 0.674 | 0.015 | 0.029 | 0.351 | 0.042 | 0.004 | 0.009 | 0.353 | 0.397 |
| | 250 | 0.736 | 0.748 | 0.015 | 0.026 | 0.516 | 0.037 | 0.005 | 0.008 | 0.528 | 0.547 |
| | 500 | 0.820 | 0.776 | 0.015 | 0.024 | 0.626 | 0.037 | 0.005 | 0.007 | 0.645 | 0.630 |
| 10 | 50 | 0.447 | 0.532 | 0.010 | 0.016 | 0.234 | 0.022 | 0.004 | 0.005 | 0.229 | 0.282 |
| | 100 | 0.593 | 0.659 | 0.010 | 0.015 | 0.348 | 0.020 | 0.005 | 0.004 | 0.350 | 0.390 |
| | 250 | 0.744 | 0.763 | 0.009 | 0.013 | 0.512 | 0.019 | 0.004 | 0.004 | 0.524 | 0.538 |
| | 500 | 0.830 | 0.805 | 0.009 | 0.012 | 0.626 | 0.018 | 0.004 | 0.004 | 0.642 | 0.628 |
| 20 | 50 | 0.447 | 0.528 | 0.009 | 0.012 | 0.230 | 0.015 | 0.004 | 0.003 | 0.225 | 0.277 |
| | 100 | 0.591 | 0.642 | 0.008 | 0.012 | 0.345 | 0.014 | 0.004 | 0.002 | 0.342 | 0.391 |
| | 250 | 0.752 | 0.767 | 0.007 | 0.011 | 0.512 | 0.013 | 0.004 | 0.002 | 0.517 | 0.539 |
| | 500 | 0.838 | 0.820 | 0.007 | 0.010 | 0.624 | 0.012 | 0.003 | 0.001 | 0.639 | 0.628 |
| 30 | 50 | 0.447 | 0.527 | 0.010 | 0.011 | 0.227 | 0.013 | 0.004 | 0.002 | 0.223 | 0.278 |
| | 100 | 0.593 | 0.637 | 0.009 | 0.011 | 0.348 | 0.013 | 0.004 | 0.001 | 0.348 | 0.391 |
| | 250 | 0.752 | 0.759 | 0.007 | 0.010 | 0.512 | 0.012 | 0.004 | 0.001 | 0.523 | 0.541 |
| | 500 | 0.842 | 0.827 | 0.007 | 0.010 | 0.626 | 0.013 | 0.003 | 0.001 | 0.637 | 0.615 |
| 40 | 50 | 0.445 | 0.525 | 0.009 | 0.011 | 0.230 | 0.012 | 0.004 | 0.002 | 0.221 | 0.277 |
| | 100 | 0.591 | 0.638 | 0.009 | 0.011 | 0.342 | 0.012 | 0.004 | 0.001 | 0.345 | 0.390 |
| | 250 | 0.751 | 0.758 | 0.007 | 0.010 | 0.508 | 0.009 | 0.004 | 0.001 | 0.519 | 0.541 |
| | 500 | 0.844 | 0.828 | 0.006 | 0.009 | 0.622 | 0.011 | 0.004 | 0.001 | 0.639 | 0.624 |

*Default denotes MAST was implemented without random-effects, RE denotes random-effects, Corrected denotes data was batch-corrected for individual prior to analysis without using individual as a random-effect, GLM denotes generalized linear model, and GLMM denotes generalized linear mixed-effects model.

**Two-part Hurdle model as implemented in MAST, Tweedie distribution as implemented in 'glmmTMB', GEE1 as implemented in 'geepack', Pseudobulk averaged or summed across cells within an individual and was implemented in DESeq2, Modified t as implemented in ROTS, and Tobit as implemented in Monocle.

**Table S3.** Type I error rates of some of the currently applied tools in single-cell analysis. Type I error rates of ten different methods under twenty different conditions and a significance threshold of $p<0.001$. 250,000 iterations were computed to obtain an error rate for each method. The inflated type I error rates computed with mixed models at the lower numbers of individuals per group are a consequence of the two-part hurdle model simultaneously testing two hypotheses and an overabundance of sub-sampling with small sample sizes. Type I error rates are well controlled for with mixed models and pseudobulk methods, while type I error rates inflate with other methods as additional independent samples or more cells are added. Pseudobulk methods are overly conservative.

| | | Two-part Hurdle | | | Tweedie | | | Pseudobulk | | | |
|---|---|---|---|---|---|---|---|---|---|---|---|
| $N_{ind}$ | $N_{cells}$ | **Default** | **Corrected** | **RE** | **GLMM** | **GLM** | **GEE1** | **Mean** | **Sum** | **Tobit** | **Modified *t*** |
| 5 | 50 | 3.40x10$^{-1}$ | 4.84x10$^{-1}$ | 2.27x10$^{-3}$ | 8.79x10$^{-3}$ | 1.48x10$^{-1}$ | 1.65x10$^{-2}$ | 8.43x10$^{-4}$ | 3.53x10$^{-3}$ | 1.39x10$^{-1}$ | 1.94x10$^{-1}$ |
| | 100 | 4.87x10$^{-1}$ | 6.22x10$^{-1}$ | 1.84x10$^{-3}$ | 7.98x10$^{-3}$ | 2.53x10$^{-1}$ | 1.46x10$^{-2}$ | 4.96x10$^{-4}$ | 2.66x10$^{-3}$ | 2.45x10$^{-1}$ | 2.99x10$^{-1}$ |
| | 250 | 6.66x10$^{-1}$ | 7.18x10$^{-1}$ | 1.96x10$^{-3}$ | 7.55x10$^{-3}$ | 4.23x10$^{-1}$ | 1.20x10$^{-2}$ | 7.36x10$^{-4}$ | 1.76x10$^{-3}$ | 4.25x10$^{-1}$ | 4.52x10$^{-1}$ |
| | 500 | 7.71x10$^{-1}$ | 7.52x10$^{-1}$ | 2.19x10$^{-3}$ | 6.85x10$^{-3}$ | 5.46x10$^{-1}$ | 1.22x10$^{-2}$ | 6.97x10$^{-4}$ | 1.56x10$^{-3}$ | 5.59x10$^{-1}$ | 5.40x10$^{-1}$ |
| 10 | 50 | 3.39x10$^{-1}$ | 4.53x10$^{-1}$ | 1.03x10$^{-3}$ | 2.81x10$^{-3}$ | 1.46x10$^{-1}$ | 4.81x10$^{-3}$ | 5.10x10$^{-4}$ | 1.35x10$^{-3}$ | 1.36x10$^{-1}$ | 1.88x10$^{-1}$ |
| | 100 | 4.92x10$^{-1}$ | 5.87x10$^{-1}$ | 9.39x10$^{-4}$ | 2.58x10$^{-3}$ | 2.49x10$^{-1}$ | 3.92x10$^{-3}$ | 6.48x10$^{-4}$ | 8.32x10$^{-4}$ | 2.41x10$^{-1}$ | 2.93x10$^{-1}$ |
| | 250 | 6.71x10$^{-1}$ | 7.29x10$^{-1}$ | 8.41x10$^{-4}$ | 1.97x10$^{-3}$ | 4.18x10$^{-1}$ | 3.57x10$^{-3}$ | 3.41x10$^{-4}$ | 6.46x10$^{-4}$ | 4.21x10$^{-1}$ | 4.41x10$^{-1}$ |
| | 500 | 7.78x10$^{-1}$ | 7.82x10$^{-1}$ | 9.53x10$^{-4}$ | 1.68x10$^{-3}$ | 5.42x10$^{-1}$ | 2.86x10$^{-3}$ | 3.14x10$^{-4}$ | 6.46x10$^{-4}$ | 5.54x10$^{-1}$ | 5.39x10$^{-1}$ |
| 20 | 50 | 3.39x10$^{-1}$ | 4.50x10$^{-1}$ | 9.28x10$^{-4}$ | 1.47x10$^{-3}$ | 1.43x10$^{-1}$ | 2.11x10$^{-3}$ | 3.79x10$^{-4}$ | 6.48x10$^{-4}$ | 1.34x10$^{-1}$ | 1.83x10$^{-1}$ |
| | 100 | 4.92x10$^{-1}$ | 5.78x10$^{-1}$ | 6.90x10$^{-4}$ | 1.35x10$^{-3}$ | 2.48x10$^{-1}$ | 2.02x10$^{-3}$ | 5.05x10$^{-4}$ | 3.11x10$^{-4}$ | 2.36x10$^{-1}$ | 2.90x10$^{-1}$ |
| | 250 | 6.76x10$^{-1}$ | 7.18x10$^{-1}$ | 5.88x10$^{-4}$ | 9.21x10$^{-4}$ | 4.17x10$^{-1}$ | 1.31x10$^{-3}$ | 3.53x10$^{-4}$ | 1.94x10$^{-4}$ | 4.15x10$^{-1}$ | 4.45x10$^{-1}$ |
| | 500 | 7.84x10$^{-1}$ | 7.93x10$^{-1}$ | 7.33x10$^{-4}$ | 9.09x10$^{-4}$ | 5.40x10$^{-1}$ | 1.15x10$^{-3}$ | 2.18x10$^{-4}$ | 1.29x10$^{-4}$ | 5.51x10$^{-1}$ | 5.37x10$^{-1}$ |
| 30 | 50 | 3.37x10$^{-1}$ | 4.46x10$^{-1}$ | 7.98x10$^{-4}$ | 8.64x10$^{-4}$ | 1.39x10$^{-1}$ | 1.49x10$^{-3}$ | 3.58x10$^{-4}$ | 2.27x10$^{-4}$ | 1.31x10$^{-1}$ | 1.86x10$^{-1}$ |
| | 100 | 4.92x10$^{-1}$ | 5.70x10$^{-1}$ | 5.93x10$^{-4}$ | 8.62x10$^{-4}$ | 2.47x10$^{-1}$ | 1.22x10$^{-3}$ | 4.04x10$^{-4}$ | 1.55x10$^{-4}$ | 2.40x10$^{-1}$ | 2.91x10$^{-1}$ |
| | 250 | 6.76x10$^{-1}$ | 7.06x10$^{-1}$ | 7.01x10$^{-4}$ | 9.36x10$^{-4}$ | 4.16x10$^{-1}$ | 9.74x10$^{-4}$ | 3.66x10$^{-4}$ | 8.40x10$^{-5}$ | 4.21x10$^{-1}$ | 4.44x10$^{-1}$ |
| | 500 | 7.88x10$^{-1}$ | 7.96x10$^{-1}$ | 6.14x10$^{-4}$ | 8.30x10$^{-4}$ | 5.42x10$^{-1}$ | 1.64x10$^{-3}$ | 2.46x10$^{-4}$ | 2.58x10$^{-5}$ | 5.50x10$^{-1}$ | 5.23x10$^{-1}$ |
| 40 | 50 | 3.37x10$^{-1}$ | 4.45x10$^{-1}$ | 7.15x10$^{-4}$ | 1.19x10$^{-3}$ | 1.42x10$^{-1}$ | 1.54x10$^{-3}$ | 3.90x10$^{-4}$ | 1.42x10$^{-4}$ | 1.30x10$^{-1}$ | 1.84x10$^{-1}$ |
| | 100 | 4.90x10$^{-1}$ | 5.71x10$^{-1}$ | 8.31x10$^{-4}$ | 1.20x10$^{-3}$ | 2.43x10$^{-1}$ | 1.20x10$^{-3}$ | 3.46x10$^{-4}$ | 1.36x10$^{-4}$ | 2.37x10$^{-1}$ | 2.92x10$^{-1}$ |
| | 250 | 6.76x10$^{-1}$ | 7.08x10$^{-1}$ | 4.94x10$^{-4}$ | 8.58x10$^{-4}$ | 4.13x10$^{-1}$ | 1.20x10$^{-3}$ | 2.84x10$^{-4}$ | 9.04x10$^{-5}$ | 4.14x10$^{-1}$ | 4.44x10$^{-1}$ |
| | 500 | 7.90x10$^{-1}$ | 7.90x10$^{-1}$ | 4.46x10$^{-4}$ | 5.21x10$^{-4}$ | 5.40x10$^{-1}$ | 8.39x10$^{-4}$ | 3.40x10$^{-4}$ | 3.90x10$^{-5}$ | 5.51x10$^{-1}$ | 5.33x10$^{-1}$ |

*Default denotes MAST was implemented without random-effects, RE denotes random-effects, Corrected denotes data was batch-corrected for individual prior to analysis without using individual as a random-effect, GLM denotes generalized linear model, and GLMM denotes generalized linear mixed-effects model.
**Two-part Hurdle model as implemented in MAST, Tweedie distribution as implemented in 'glmmTMB', GEE1 as implemented in 'geepack', Pseudobulk averaged or summed across cells within an individual and was implemented in DESeq2, Modified t as implemented in ROTS, and Tobit as implemented in Monocle.

**Table S4.** Type I error rates of some of the currently applied tools in single-cell analysis. Type I error rates of ten different methods under twenty different conditions and a significance threshold of $p<0.0001$. 250,000 iterations were computed to obtain an error rate for each method. The inflated type I error rates computed with mixed models at the lower numbers of individuals per group are a consequence of the two-part hurdle model simultaneously testing two hypotheses and an overabundance of sub-sampling with small sample sizes. Type I error rates are well controlled for with mixed models and pseudobulk methods, while type I error rates inflate with other methods as additional independent samples or more cells are added. Pseudobulk methods are overly conservative.

| | | Two-part Hurdle | | | Tweedie | | | Pseudobulk | | | |
|---|---|---|---|---|---|---|---|---|---|---|---|
| $N_{ind}$ | $N_{cells}$ | Default | Corrected | RE | GLMM | GLM | GEE1 | Mean | Sum | Tobit | Modified *t* |
| 5 | 50 | 2.68x10$^{-1}$ | 4.18x10$^{-1}$ | 4.35x10$^{-4}$ | 3.68x10$^{-3}$ | 9.93x10$^{-2}$ | 7.71x10$^{-3}$ | 1.67x10$^{-4}$ | 1.76x10$^{-3}$ | 8.79x10$^{-2}$ | 1.40x10$^{-1}$ |
| | 100 | 4.16x10$^{-1}$ | 5.74x10$^{-1}$ | 2.65x10$^{-4}$ | 2.79x10$^{-3}$ | 1.95x10$^{-1}$ | 6.28x10$^{-3}$ | 8.72x10$^{-5}$ | 1.15x10$^{-3}$ | 1.79x10$^{-1}$ | 2.36x10$^{-1}$ |
| | 250 | 6.11x10$^{-1}$ | 6.94x10$^{-1}$ | 3.44x10$^{-4}$ | 3.18x10$^{-3}$ | 3.55x10$^{-1}$ | 4.83x10$^{-3}$ | 1.91x10$^{-4}$ | 6.55x10$^{-4}$ | 3.53x10$^{-1}$ | 3.86x10$^{-1}$ |
| | 500 | 7.28x10$^{-1}$ | 7.34x10$^{-1}$ | 3.60x10$^{-4}$ | 2.48x10$^{-3}$ | 4.85x10$^{-1}$ | 4.98x10$^{-3}$ | 1.41x10$^{-4}$ | 5.61x10$^{-4}$ | 4.95x10$^{-1}$ | 4.75x10$^{-1}$ |
| 10 | 50 | 2.67x10$^{-1}$ | 3.96x10$^{-1}$ | 1.12x10$^{-4}$ | 7.80x10$^{-4}$ | 9.80x10$^{-2}$ | 1.23x10$^{-3}$ | 6.45x10$^{-5}$ | 7.84x10$^{-4}$ | 8.69x10$^{-2}$ | 1.35x10$^{-1}$ |
| | 100 | 4.20x10$^{-1}$ | 5.31x10$^{-1}$ | 9.76x10$^{-5}$ | 6.11x10$^{-4}$ | 1.88x10$^{-1}$ | 9.31x10$^{-4}$ | 9.31x10$^{-5}$ | 3.97x10$^{-4}$ | 1.74x10$^{-1}$ | 2.30x10$^{-1}$ |
| | 250 | 6.11x10$^{-1}$ | 6.97x10$^{-1}$ | 9.89x10$^{-5}$ | 3.94x10$^{-4}$ | 3.51x10$^{-1}$ | 9.24x10$^{-4}$ | 8.00x10$^{-5}$ | 2.38x10$^{-4}$ | 3.46x10$^{-1}$ | 3.74x10$^{-1}$ |
| | 500 | 7.34x10$^{-1}$ | 7.65x10$^{-1}$ | 6.78x10$^{-5}$ | 4.12x10$^{-4}$ | 4.79x10$^{-1}$ | 6.09x10$^{-4}$ | 5.18x10$^{-5}$ | 1.81x10$^{-4}$ | 4.88x10$^{-1}$ | 4.72x10$^{-1}$ |
| 20 | 50 | 2.67x10$^{-1}$ | 3.91x10$^{-1}$ | 5.75x10$^{-5}$ | 2.00x10$^{-4}$ | 9.65x10$^{-2}$ | 3.29x10$^{-4}$ | 5.59x10$^{-5}$ | 4.28x10$^{-4}$ | 8.44x10$^{-2}$ | 1.32x10$^{-1}$ |
| | 100 | 4.19x10$^{-1}$ | 5.28x10$^{-1}$ | 4.19x10$^{-5}$ | 1.58x10$^{-4}$ | 1.85x10$^{-1}$ | 3.00x10$^{-4}$ | 8.52x10$^{-5}$ | 1.50x10$^{-4}$ | 1.69x10$^{-1}$ | 2.26x10$^{-1}$ |
| | 250 | 6.16x10$^{-1}$ | 6.73x10$^{-1}$ | 6.82x10$^{-5}$ | 1.95x10$^{-4}$ | 3.49x10$^{-1}$ | 1.51x10$^{-4}$ | 3.62x10$^{-5}$ | 4.53x10$^{-5}$ | 3.41x10$^{-1}$ | 3.77x10$^{-1}$ |
| | 500 | 7.39x10$^{-1}$ | 7.68x10$^{-1}$ | 6.81x10$^{-5}$ | 1.15x10$^{-4}$ | 4.77x10$^{-1}$ | 1.49x10$^{-4}$ | 2.87x10$^{-5}$ | 2.59x10$^{-5}$ | 4.83x10$^{-1}$ | 4.73x10$^{-1}$ |
| 30 | 50 | 2.67x10$^{-1}$ | 3.87x10$^{-1}$ | 1.10x10$^{-4}$ | 1.14x10$^{-4}$ | 9.27x10$^{-2}$ | 1.64x10$^{-4}$ | 4.21x10$^{-5}$ | 1.10x10$^{-4}$ | 8.35x10$^{-2}$ | 1.33x10$^{-1}$ |
| | 100 | 4.18x10$^{-1}$ | 5.20x10$^{-1}$ | 6.85x10$^{-5}$ | 1.14x10$^{-4}$ | 1.85x10$^{-1}$ | 1.49x10$^{-4}$ | 9.20x10$^{-5}$ | 8.44x10$^{-5}$ | 1.74x10$^{-1}$ | 2.26x10$^{-1}$ |
| | 250 | 6.15x10$^{-1}$ | 6.66x10$^{-1}$ | 1.10x10$^{-4}$ | 1.24x10$^{-4}$ | 3.49x10$^{-1}$ | 7.95x10$^{-5}$ | 1.44x10$^{-5}$ | 1.94x10$^{-5}$ | 3.45x10$^{-1}$ | 3.77x10$^{-1}$ |
| | 500 | 7.42x10$^{-1}$ | 7.65x10$^{-1}$ | 2.73x10$^{-5}$ | 2.34x10$^{-5}$ | 4.80x10$^{-1}$ | - | 2.88x10$^{-5}$ | 6.46x10$^{-6}$ | 4.82x10$^{-1}$ | 4.57x10$^{-1}$ |
| 40 | 50 | 2.64x10$^{-1}$ | 3.84x10$^{-1}$ | 1.48x10$^{-4}$ | 1.53x10$^{-4}$ | 9.25x10$^{-2}$ | 1.51x10$^{-4}$ | 4.18x10$^{-5}$ | 7.78x10$^{-5}$ | 8.06x10$^{-2}$ | 1.31x10$^{-1}$ |
| | 100 | 4.17x10$^{-1}$ | 5.19x10$^{-1}$ | 5.38x10$^{-5}$ | 1.76x10$^{-4}$ | 1.82x10$^{-1}$ | 1.89x10$^{-4}$ | 3.53x10$^{-5}$ | 3.21x10$^{-5}$ | 1.71x10$^{-1}$ | 2.29x10$^{-1}$ |
| | 250 | 6.16x10$^{-1}$ | 6.70x10$^{-1}$ | 6.87x10$^{-5}$ | 8.00x10$^{-5}$ | 3.45x10$^{-1}$ | - | 2.83x10$^{-5}$ | 1.30x10$^{-5}$ | 3.40x10$^{-1}$ | 3.76x10$^{-1}$ |
| | 500 | 7.44x10$^{-1}$ | 7.58x10$^{-1}$ | 4.15x10$^{-5}$ | 1.26x10$^{-4}$ | 4.78x10$^{-1}$ | - | 4.36x10$^{-5}$ | 1.32x10$^{-5}$ | 4.85x10$^{-1}$ | 4.66x10$^{-1}$ |

*Default denotes MAST was implemented without random-effects, RE denotes random-effects, Corrected denotes data was batch-corrected for individual prior to analysis without using individual as a random-effect, GLM denotes generalized linear model, and GLMM denotes generalized linear mixed-effects model.

**Two-part Hurdle model as implemented in MAST, Tweedie distribution as implemented in ‘glmmTMB’, GEE1 as implemented in ‘geepack’, Pseudobulk averaged or summed across cells within an individual and was implemented in DESeq2, Modified t as implemented in ROTS, and Tobit as implemented in Monocle.
